# PFKFB3 Depletion Activates β-Cell Replication by Cell Competitive Culling of Compromised β-Cells Under Stress

**DOI:** 10.1101/2021.04.07.438857

**Authors:** Jie Min, Feiyang Ma, Matteo Pellegrini, Oppel Greeff, Salvador Moncada, Slavica Tudzarova

## Abstract

Highly conserved hypoxia-inducible factor 1 alpha (HIF1α) and its target 6-phosphofructo-2-kinase/fructose-2,6-biphosphatase 3 (PFKFB3) play a critical role in the survival of damaged β-cells in type 2 diabetes (T2D) while rendering β-cells non-responsive to glucose stimulation by mitochondrial suppression. HIF1α-PFKFB3 is activated in 30-50% of all β-cells in diabetic islets, leaving an open question of whether targeting this pathway may adjust β-cell mass and function to the specific metabolic demands during diabetogenic stress.

Our previous studies of β-cells under amyloidogenic stress by human islet amyloid polypeptide (hIAPP) revealed that PFKFB3 is a metabolic execution arm of the HIF1α pathway with potent implications on Ca^2+^ homeostasis, metabolome, and mitochondrial form and function.

To discriminate the role of PFKFB3 from HIF1α *in vivo*, we generated mice with conditional β-cell specific disruption of the *Pfkfb3* gene on a hIAPP^+/-^ background and a high-fat diet (HFD) [PFKFB3^βKO^ + diabetogenic stress (DS)].

PFKFB3 disruption in β-cells under diabetogenic stress led to selective purging of hIAPP-damaged β-cells and the disappearance of bihormonal insulin- and glucagon-positive cells, thus compromised β-cells. At the same time, PFKFB3 disruption led to a three-fold increase in β-cell replication resembling control levels as measured with minichromosome maintenance 2 protein (MCM2). PFKFB3 disruption depleted bihormonal cells while increased β-cell replication that was reflected in the increased β-/α-cell ratio and maintained β-cell mass. Analysis of metabolic performance indicated comparable glucose intolerance and reduced plasma insulin levels in PFKFB3^βKO^ DS relative to PFKFB3^WT^ DS mice. In the PFKFB3^βKO^ DS group, plasma glucagon levels were reduced compared to PFKFB3^WT^ DS mice and were in line with increased insulin sensitivity. Glucose intolerance in PFKFB3^βKO^ DS mice could be explained by the compensatory expression of HIF1α after disruption of PFKFB3. Our data strongly suggest that the replication and functional recovery of β-cells under diabetogenic stress depend on selective purification of HIF1α and PFKFB3-positive β-cells. Thus, HIF1α-PFKFB3-dependent activation of cell competition and purging of compromised β-cells may yield functional competent β-cell mass in diabetes.

## INTRODUCTION

Regenerative growth that exceeds physiological constraints is triggered by the increase in metabolic demands, and it is a feature of adaptive processes whose outcome determines the development of various diseases including diabetes [1, 2]. Regenerative β-cell growth is typically triggered during pregnancy to compensate for the increased metabolic load of a developing fetus [3], or in nondiabetic obese individuals in response to insulin resistance [4]. Diabetogenic stress, such as the accumulation of the toxic oligomers of human islet amyloid polypeptide (hIAPP) [5-7] poses a particular challenge for long-lived and highly specialized pancreatic β-cells. With progressive accumulation of hIAPP toxic oligomers, β-cells have to work harder to consolidate β-cell mass with the specialized function in order to maintain euglycemia [2]. This balance typically falls short in type 2 diabetes (T2D) [8] but it cannot be explained by the deficit in the β-cell mass alone since β-cell mass is rather preserved (from 35% to 76%) [6, 9-12]. In contrast to β- cell mass, β-cell glucose responsiveness declines even prior to the onset of T2D. This suggests that, although viable, most of the β-cells in T2D fail to adapt to the growing metabolic demands under stress and are dysfunctional.

We demonstrated previously that damaged β-cells activate a highly conserved hypoxia-inducible factor 1 alpha (HIF1α) metabolic pathway and become entrapped via high glycolysis disengaged from the TCA cycle that ensures the survival of damaged and dysfunctional β-cells [13]. The impact on glucose metabolism by HIF1α in β-cells was executed via 6-phosphofructo-2-kinase/fructose-2,6-biphosphatase 3 (PFKFB3), a key regulator of β-cell aerobic glycolysis under stress [13, 14]. We reported that activation of HIF1α-PFKFB3 pathway in prediabetes predicted chronic glycolytic energy production that further exacerbates the slip of β-cells into the decompensation and diabetes progression [13, 15, 16]. Dysfunctional β-cells that can sustain damage with activation of HIF1α-PFKFB3 metabolic remodeling constituted one-third to one-fifth of all β-cells in humans with T2D [13]. They closely resembled suboptimal or unfit (compromised) cells that, by increasing aerobic glycolysis, have escaped the fitness quality control of the evolutionarily conserved cell competition [17-19]. Since the survival of damaged β-cells in T2D depends on the glycolytic HIF1α-PFKFB3 pathway [13], targeting this pathway may differentially affect the fate of healthy versus damaged β-cells.

Based on the new evidence, we propose that the relationship between heterogeneous β- cell subpopulations that result from some β-cells being subjected to injury is operative in β-cell replenishment by cellular replication. In our study, conditional knockout of PFKFB3 in adult β-cells under high diabetogenic stress (hIAPP and high-fat diet) (PFKFB3^βKO^ DS) led to selective elimination or purging of damaged β-cells and double-insulin and glucagon positive (bihormonal) cells and increased the replication of remnant healthy β- cells. Despite the β-cell mass being maintained, PFKFB3^βKO^ DS mice still showed impaired glucose tolerance with independent expression of HIF1α.

We analyzed β-cells with HIF1α (lactate dehydrogenase positive) signature in humans and compared them to bihormonal β-cells in our mouse model of diabetes. The loss of PFKFB3 during diabetogenic stress induced the clearance of bihormonal and damaged β-cells but it was insufficient to suppress the HIF1α expression in β-cells. β-cells with HIF1α signature in the absence of PFKFB3 were probably responsible for impaired glucose tolerance.

In this study we sought to dissect the roles of HIF1α and PFKFB3 in β-cell regeneration under diabetogenic stress.

## MATERIALS AND METHODS

### Animals

Homozygous hIAPP^+/+^ mice were a gift from Dr Peter Butler’s laboratory and were previously described [20]. We have generated a β-cell specific inducible PFKFB3 knockout mouse model (RIP-CreERT:PFKFB3^fl/fl^) by crossing mice that carry the floxed *Pfkfb3* gene (JAX Laboratories) with mice that express Cre recombinase under the control of the rat insulin promoter (RIP-CreERT). We have crossed mice on a homozygous hIAPP^+/+^ background with either PFKFB3^fl/f^ or RIP-CreERT mice and then crossed PFKFB3^fl/fl^ hIAPP^+/-^ and PFKFB3^fl/fl^ RIP-CreERT mice together to generate the three experimental genotypes: RIP-CreERT PFKFB3^fl/fl^ hIAPP^-/-^, RIP-CreERT PFKFB3^wt/wt^ hIAPP^+/-^ and RIP-CreERT PFKFB3^fl/fl^ hIAPP^+/-^ hereto referred as PFKFB3^WT^ hIAPP^-/-^ (WT), PFKFB3^WT^ hIAPP^+/-^ (PFKFB3^WT^ diabetogenic stress, DS), and PFKFB3^βKO^ hIAPP^+/-^ (PFKFB3^βKO^ DS). All experimental groups were subjected to high fat diet. Cre-loxP recombination of the floxed sites in *Pfkfb3* was induced by intraperitoneal tamoxifen injection at the age of 20-27 weeks. The mice were given a chow diet for 10 weeks post tamoxifen injection and then all mice were exposed to a high-fat diet for another 13 weeks (HFD, Research Diets Inc, New Brunswick, NJ, USA) to induce diabetes in combination with hIAPP^+/-^, since only male mice homozygous for hIAPP (hIAPP^+/+^) develop diabetes spontaneously [20]. The mice were maintained on a 12 hours day/night cycle at the UCLA Institutional Animal Care and Use Committee (ARC) approved mice colony facility. At 30-37 weeks of age, all mice were assigned to receive a diet containing high amounts of fat (35% w/w or 60% calories from fat;; D12492). The fat composition of the high-fat diet was 32.2% saturated, 35.9% monounsaturated, and 31.9% polyunsaturated fats. Mice had *ad libitum* access to diet and water for the duration of the study. Bodyweight and fasting blood glucose levels were assessed weekly, with additional measurements being made on days that included glucose and insulin tolerance tests (IP-GTT and ITT, respectively).

### Insulin and glucose tolerance tests

An intraperitoneal glucose tolerance test (IP-GTT) was performed at 9 and 12 weeks after HFD (19 and 22 weeks post tamoxifen injection). Tail vein blood glucose was collected prior to and 15, 30, 60, 90, 120 minutes post glucose bolus injection. Retro-orbital bleeding was used to collect the blood for the second IP-GTT prior to and 30 minutes after glucose bolus injection. The mice were anaesthetised by brief exposure to isoflurane (10 seconds). The blood was collected in an EDTA-coated microcentrifuge tube and the plasma was obtained by centrifuging the samples for 10 minutes (5000 RCF, 10 min, 4°C).

### Glucose and insulin assays

Fasted blood glucose was measured weekly after overnight fasting for 18 hours (after the regular change of cages and bedding and withdrawal of food while providing water *ad libitum*) from a tail drawn blood using a freestyle blood glucose meter (Abbott Diabetes Care Inc, Alameda, CA, USA). When blood glucose exceeded the detection range of the blood glucose meter, plasma glucose was determined using the glucose oxidase method and analyzed with YSI 2300 STAT PLUS Glucose and L-Lactate Analyzer.

Insulin, C-peptide, and glucagon levels in plasma were determined using ultrasensitive ELISA for mouse insulin (Mercodia 10-1247-01, Uppsala, Sweden), mouse C-peptide (Crystal Chem 90050, IL, USA), and mouse glucagon (Mercodia 10-1281-01, Uppsala, Sweden).

Ten weeks after HFD (19 weeks after tamoxifen injection), an intraperitoneal insulin tolerance test (0.75 IU/kg) (Lilly insulin Lispro, LLC, Indianapolis, USA) was performed in conscious mice fasted for 6 hours. Tail vein blood was collected prior to and at 0, 20, 40, 60 minutes post insulin administration for glucose measurements.

### Pancreas perfusion and isolation

One week following IP-GTT and ITT, mice were euthanised by cervical dislocation. A medial cut was used to open the abdomen and chest cavities, while a cut of the right ventricle was followed with a poke of the left ventricle with a needle to inject 10 ml cold phosphate buffered saline (PBS) slowly for perfusion of the pancreas. After perfusion, the pancreas was placed in cold PBS and separated from other tissue including the surrounding fat. The pancreas was then weighed after absorbing the excess PBS with tissue.

### Histological assessments

After excision of smaller pieces, the pancreas was fixed in 4% paraformaldehyde (Electron Microscopy Sciences 19202, Hatfield, PA, USA) overnight at 4°C, paraffin-embedded, and sectioned at 4  μm thickness. For β-cell area, peroxidase and haematoxylin staining were performed on deparaffinised sections that were sequentially incubated with rabbit anti-insulin antibody (Cell Signaling Technology C27C9, Danvers, MA, USA, 1:400), then with F(ab’)_2_ conjugates with Biotin-SP (Jackson ImmunoResearch 711-066-152, West Grove, PA, USA, 1:100 for IHC), after which steps the VECTASTAIN ABC Kit (HRP) (Vector Laboratories PK-4000, Burlingame, CA, USA), the DAB substrate Kit (HRP) (Vector Laboratories SK-4100, Burlingame, CA, USA), and Harris Haematoxylin were applied prior to mounting the sections with Permount (Fisher SP15-100, Hampton, NH, USA). Morphometric analyses were performed using Image-Pro Plus 5.1 software on the Olympus IX70 inverted tissue culture microscope (Olympus, Center Valley, PA, USA). Imaging and data analysis were performed by two observers in a blinded fashion for the experimental mouse genotype of each section. The islet edges were manually circumscribed using a multichannel image. Insulin- and haematoxylin-positive areas were determined for each islet using pixel thresholding. The β-cell area was then calculated as insulin-positive areas/haematoxylin-positive areas * 100%.

Immunofluorescence analysis was performed in Openlab 5.5.0 software on the Leica DM6000 B research microscope. The following antibodies were used: rabbit anti-PFKFB3 (Abcam ab181861, Cambridge, MA, USA, 1:100);; mouse anti-MCM2 (BD Transduction Laboratories 610700, San Diego, CA, USA, 1:100);; rabbit anti-cleaved caspase-3 (Cell Signaling Technology 9664S, Danvers, MA, USA, 1:400);; guinea pig anti-insulin (Abcam ab195956, Cambridge, MA, USA, 1:400);; mouse anti-glucagon (Sigma-Aldrich G2654, St.Louis, MO, USA,1:1000), mouse anti-c-Myc (Santa Cruz Biotechnology Inc 9E10 sc-40, Dallas, Texas, USA, 1:100);; mouse anti-HIF1α (Novus Biologicals NB100-105, Centennial, CO, USA, 1:50). The anti-PFKFB3 staining was performed as described in [21] with some modifications. The presence of any of above stated markers in the islets was evaluated in the pancreatic sections using images of 20 – 25 islets per sample taken with a Leica DM6000 fluorescent microscope (Wetzlar, Germany). Islet was considered as a group of cells consisting of 4 or more β-cells. We used the following secondary antibodies: F(ab’)2 conjugates with FITC Donkey Anti-Guinea Pig IgG (H+L) (Jackson ImmunoResearch 706-096-148, West Grove, PA, USA, 1:200 for IF);; F(ab’)2 conjugates with Cy3 Donkey Anti-Rabbit IgG (H+L) (Jackson ImmunoResearch 711-166-152, West Grove, PA, USA, 1:200 for IF);; F(ab’)2 conjugates with Cy3 Donkey Anti-Mouse IgG (H+L) (Jackson ImmunoResearch 711-165-151, West Grove, PA, USA, 1:200 for IF) and F(ab’)2 conjugates with Alexa 647 Donkey Anti-Mouse IgG (H+L) (Jackson ImmunoResearch 715-606-150, West Grove, PA, USA, 1:100 for IF). The In Situ Cell Death Detection Kit (Roche Diagnostics Corporation 12156792910, Indianapolis, IN, USA) was used for the determination of cell death by TUNEL assay. Vectashield with DAPI (Vector Laboratories H1200, Burlingame, CA, USA) was used to mount the slides.

### Single-cell RNA sequencing data analysis

The single-cell RNA-seq dataset was obtained from GEO accession GSE124742, in which pancreatic cells from non-diabetics and type 2 diabetes (T2D) were used. To identify different cell types and find signature genes for each cell type, the R package Seurat (version 3.1.2) was used to analyze the expression matrix. Cells with less than 100 genes and 500 UMIs detected were removed from further analysis. The Seurat function NormalizeData was used to normalize the raw counts. Variable genes were identified using the FindVariableGenes function. The Seurat ScaleData function was used to scale and center expression values in the dataset for dimensionality reduction. Default parameters were used in the Seurat functions above. Principal component analysis (PCA) and uniform manifold approximation and projection (UMAP) were used to reduce the dimensions of the data, and the first two dimensions were used in plots. The FindClusters function was later used to cluster the cells. The FindAllMarkers function was used to determine the marker genes for each cluster, which were then used to define the cell types. Differential expression analysis between two groups of cells was carried out using the FindMarkers function. The Wilcoxon Rank-Sum test was performed in the differential analysis, and the Benjamini-Hochberg Procedure was applied to adjust the false discovery rate.

Data were analyzed through the use of IPA (QIAGEN Inc., https://www.qiagenbioinformatics.com/products/ingenuitypathway-analysis) [22] and Enrich***r***, the comprehensive resource for curated gene sets and a search engine available at http://amp.pharm.mssm.edu/Enrich [23-25].

### Statistical analyses

Data are presented as an error of the mean (standard error, SEM) for the number of mice indicated. For the IP-GTT and ITT, areas under the curve (AUC) for glucose, insulin, C-peptide and glucagon were calculated using the trapezoidal rule. Mean data were compared between groups by analysis using Student’s t-test. P values less than 0.05 were considered significant.

## RESULTS

### β-cells in PFKFB3^βKO^ mice demonstrate ongoing immunopositivity for HIF1α and impaired glucose tolerance

To study and dissect the role of PFKFB3 from HIF1α in the survival of damaged β-cells under diabetogenic stress *in vivo*, we generated mice with β-cell specific conditional disruption of the *Pfkfb3* gene on a *hIAPP^+/-^* background and exposed them to a high-fat diet for 13 weeks (PFKFB3^βKO^ DS). Diabetogenic stress was deemed high since it involved insulin resistance (obesity) and exposure to misfolded proteins through hIAPP^+/-^ expression, altogether with advanced age (44 to 50 weeks), known as cumulative risk factors in diabetes [26-28].

PFKFB3^fl/fl^ hIAPP^+/-^mice were born at the expected Mendelian ratio. From one week before the monitoring of mice up to the end of the experiment, there was no difference in the bodyweight among different experimental groups (Supplementary Figure 1A-D). No difference was observed in the pancreas weight, but both spleen and liver showed lower weight in PFKFB3^WT^ DS mice and PFKFB3^βKO^ DS mice compared to WT controls although not reaching a significant difference (Supplementary Figure 2A-C).

Efficient disruption of PFKFB3 expression was confirmed by PFKFB3 immunostaining of the pancreatic sections of the PFKFB3^βKO^ DS mice using PFKFB3^WT^ hIAPP^+/+^ as a positive control (Figure 1A and Supplemental Data). Diabetogenic stress led to 33.9 ± 6.4% PFKFB3 immunolabelling of β-cells in PFKFB3^WT^ DS mice (p<0.05), similar to that previously reported for humans with T2D [28]. PFKFB3 immunolabelling of β-cells from WT mice was 3.7 ± 1.9%, while in PFKFB3^βKO^ DS mice it was successfully abolished and accounted for 1.0 ± 0.8% (Figure 1A and B).

**Figure 1.**
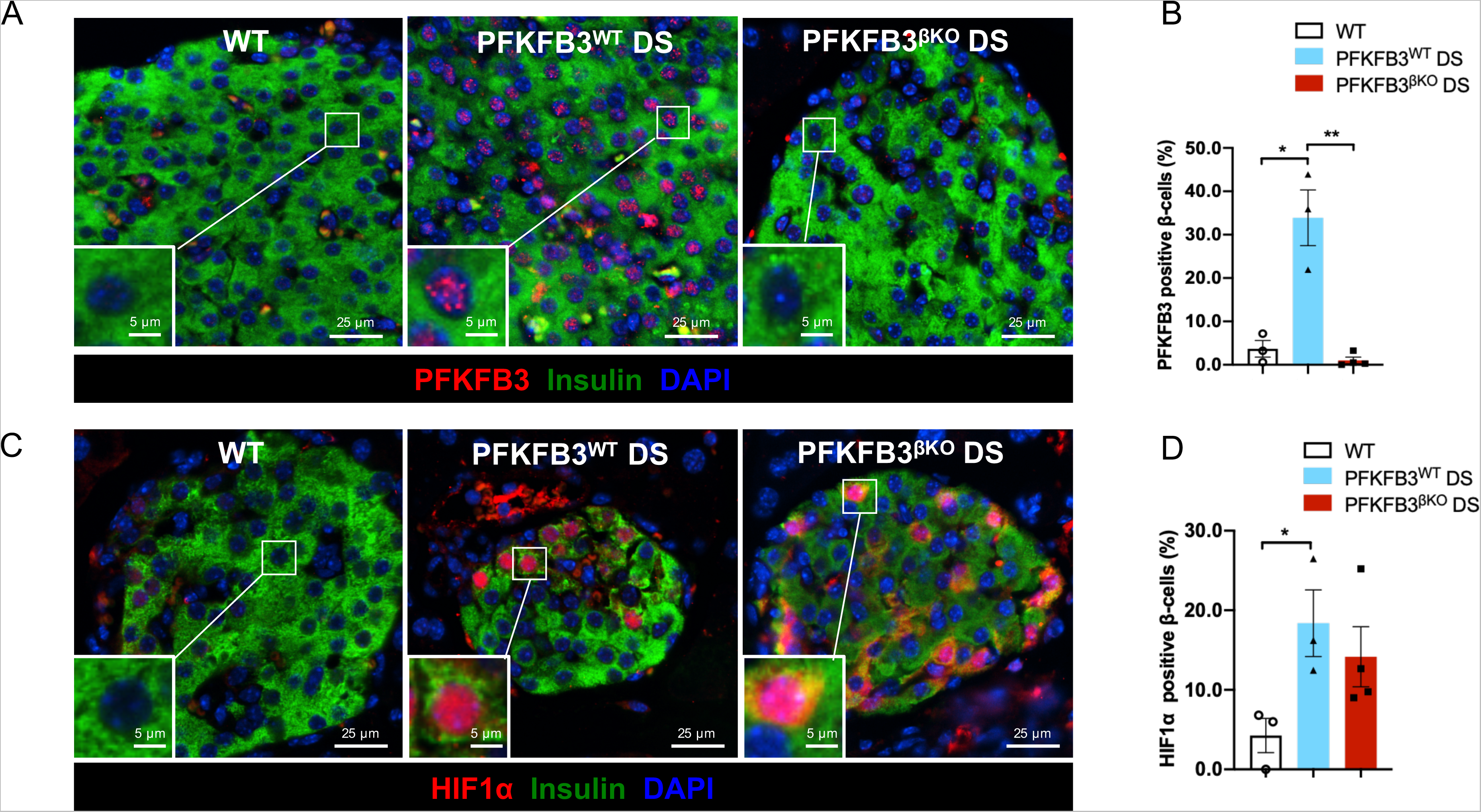
HIF1α is upregulated in PFKFB3^βKO^ DS mice. (**A**) Representative immunofluorescence images of islets from WT+HFD, PFKFB3^WT^ DS and PFKFB3^βKO^ DS mice immunostained for PFKFB3 (red), insulin (green) and nuclei (blue). (**B**) Quantification of images in A. (**C**) Representative immunofluorescence images of islets from WT+HFD, PFKFB3^WT^ and PFKFB3^βKO-^ mice on diabetogenic stress (DS) immunostained for HIF1α (red), insulin (green) and nuclei (blue). (**D**) Quantification of images in C (n=3, n=4 for PFKFB3^βKO^ DS SEM *p<0.05). HFD – high fat diet.

To establish if in this model PFKFB3-is linked to HIF1α expression, we immunostained pancreatic sections from all experimental groups with HIF1α antibody. HIF1α expression increased to 18.4 ± 4.2% in β-cells from PFKFB3^WT^ DS mice compared to WT control (*p<0.05). In PFKFB3^βKO^ DS mice, 14.2 ± 3.8% of all β-cells (Figure 1C and D) showed HIF1α immunopositive cytoplasm and nucleus (Supplemental data). This result indicated that PFKFB3 knockout triggered a compensatory HIF1α expression in response to stress.

Analysis of the metabolic performance of PFKFB3^βKO^ DS mice revealed glucose intolerance at both 9 (*p<0.05, n=4) and 12 weeks (not shown) after onset of the high-fat diet (Figure 2A, B). Insulin tolerance test indicated higher insulin sensitivity in PFKFB3^βKO^ DS mice that was found significant in younger mice (Figure 2C, D and Supplementary Figure 3). Plasma insulin levels mirrored C-peptide levels (not shown) and were lower in PFKFB3^βKO^ DS and PFKFB3^WT^ DS compared to WT controls (p<0.05) (Figure 2E). Interestingly, although plasma insulin was low for PFKFB3^βKO^ DS and PFKFB3^WT^ DS mice, the later had much higher plasma glucagon levels, while PFKFB3^βKO^ DS mice demonstrated a sharp reduction in glucagon levels in comparison to PFKFB3^WT^ DS mice and the same levels as seen in WT controls (Figure 2F). These results, together with increased insulin sensitivity, suggested impaired insulin secretion in PFKFB3^βKO^ DS mice. Next, we asked whether impaired insulin secretion in PFKFB3^βKO^ DS mice is due to a failure to expand the β-cell mass in the absence of PFKFB3.

**Figure 2.**
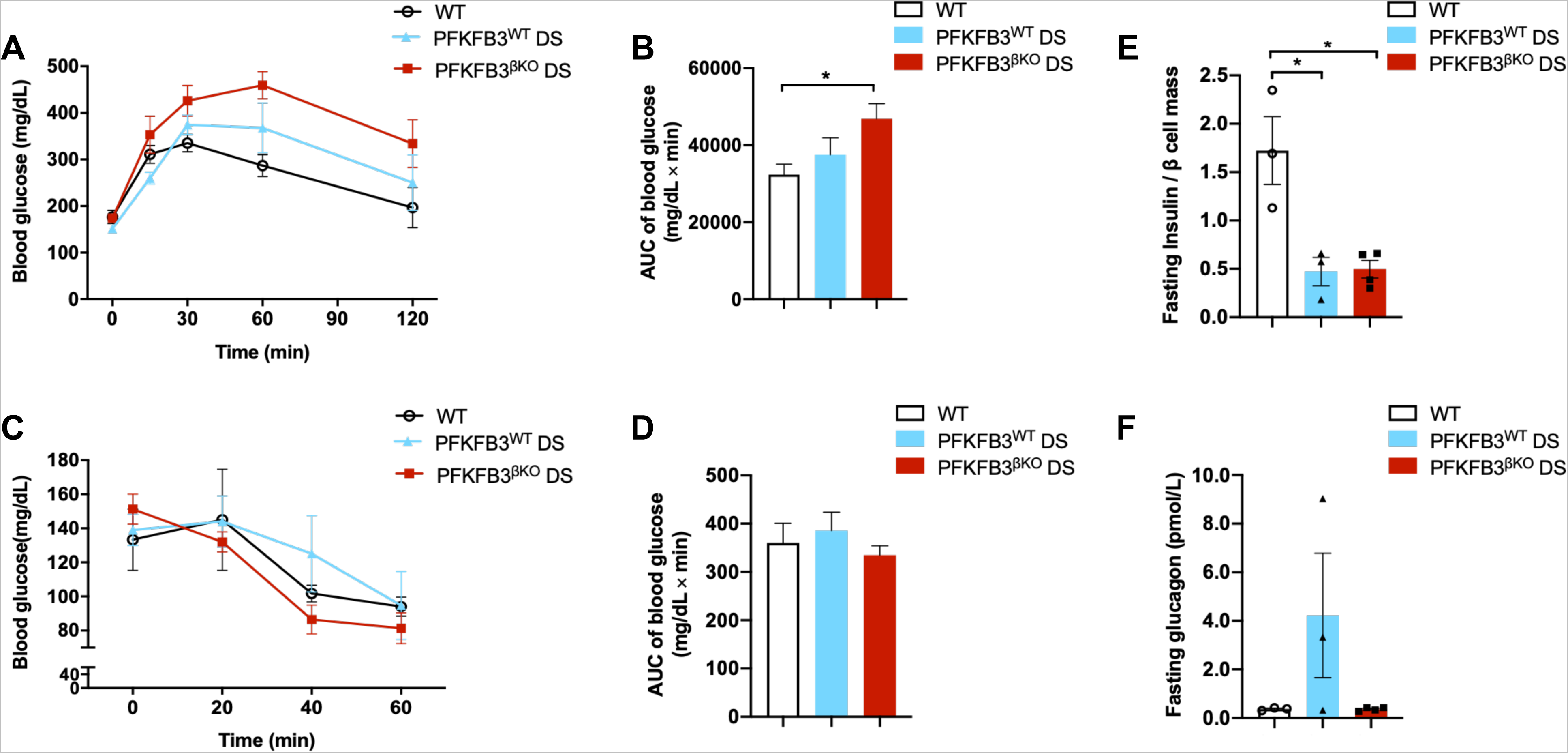
PFKFB3^βKO^ DS mice demonstrate similar impaired glucose tolerance and insulin-but reduced glucagon plasma levels relative to PFKFB3^WT^ DS mice. (**A**) Intraperitoneal glucose tolerance test (IP-GTT) at 9 weeks post-onset of high-fat diet (HFD). (**B**) Quantification of the area under the curve (AUC) as mg/dL x min in the experimental groups in A. (**C**). Insulin tolerance test at 10 weeks after onset of HFD. (**D**) Quantification of the area under the curve (AUC) as mg/dL x min in the experimental groups in E. (**E**) Fasting plasma insulin and (**F**) Fasting plasma glucagon (12 weeks post onset of HFD) (n=3, n=4 for PFKFB3^βKO^ DS, SEM *p<0.05).

### β-cells in PFKFB3^βKO^ DS mice demonstrate increased cell turn-over and replication

β-cell fractional area and mass were unaltered among the experimental groups (Figure 3A-B). To investigate growth dynamics that ultimately led to comparable β-cell mass between PFKFB3^βKO^ DS and PFKFB3^WT^ DS mice, we performed TUNEL staining as a measurement of past cell dying (Figure 3C and Supplemental data) and cleaved caspase 3 immunostaining as a measurement of active β-cell death [29] (Figure 3E-F). According to the TUNEL analysis, past β-cell death was increased in the PFKFB3^βKO^ DS mice relative to PFKFB3^WT^ DS mice (Figure 3C and Supplemental data) while no difference was measured in the ongoing cell death by cleaved caspase 3 (Figure F) compared to the other groups. To our surprise, the β-/α-cell ratio was increased in PFKFB3^βKO^ DS relative to PFKFB3^WT^ DS mice (Figure 3D) and this posed a question of its origin.

**Figure 3.**
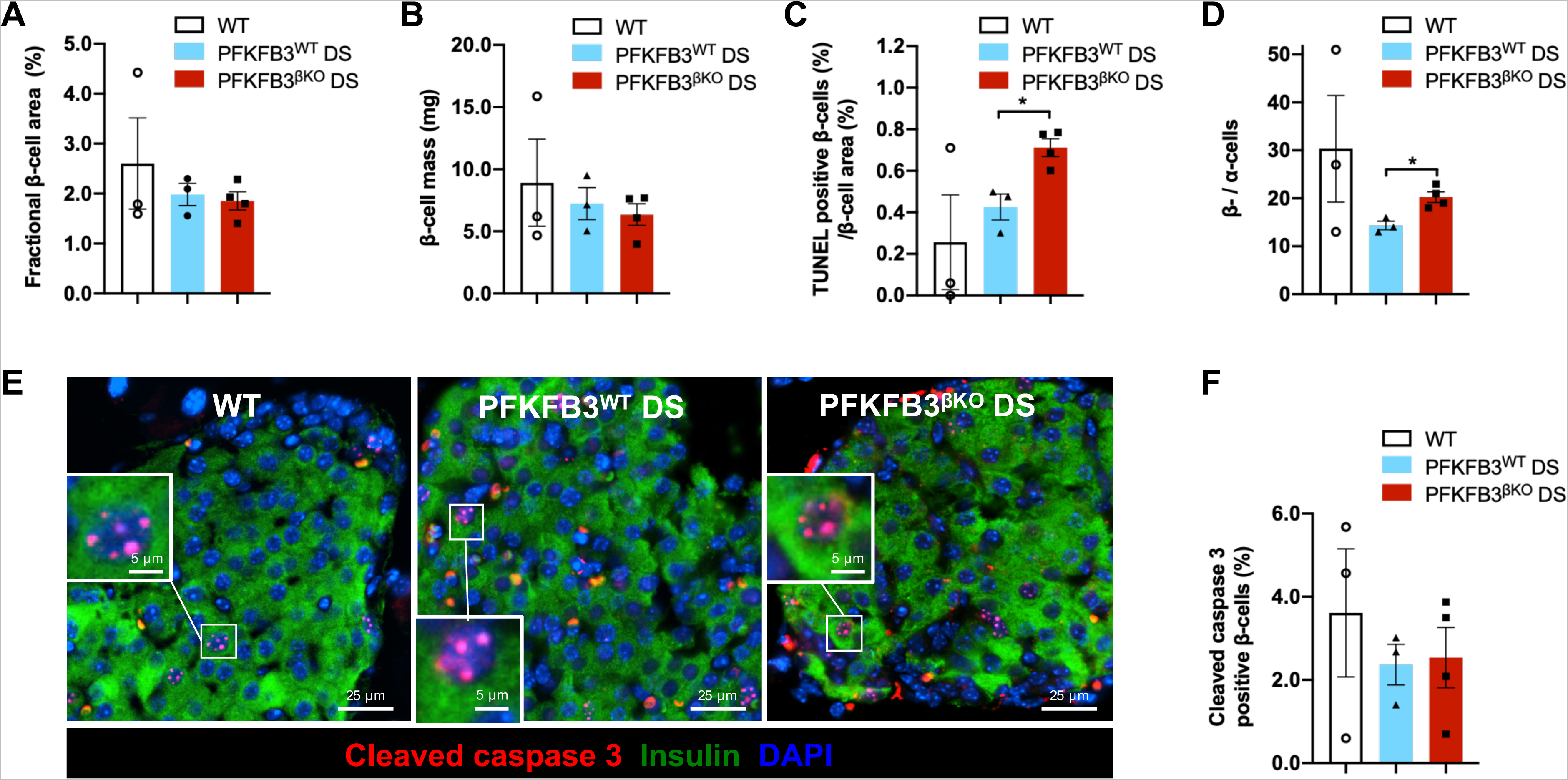
PFKFB3^βKO^ DS mice show increased β-/α-cell ratio in spite of the increase in the cell death relative to PFKFB3^WT^ DS mice. (**A**) Quantification of fractional β-cell area (%). (**B**) Quantification of β-cell mass (mg). (**C**) Quantification of β-cell death as measured by labelling with TUNEL assay (%) represented relative to fractional β-cell area. (**D**) Quantification of β-cell-relative to α-cell number in indicated experimental groups. **(E)** Representative immunofluorescence images of islets from WT+HFD, PFKFB3^WT^ DS and PFKFB3^βKO^ DS mice immunostained for cleaved caspase-3 (red), insulin (green) and nuclei (blue). **(F)** Quantification of images under E (n=3, n=4 for PFKFB3^βKO^ DS, SEM *p<0.05).

To further elucidate if the increase in the β-/α-cell ratio relied on the increased generation of β-cells, we performed immunolabelling with an early replication initiation marker, minichromosome maintenance protein 2 (MCM2) [30, 31]. Our data showed that β-cells from PFKFB3^βKO^ DS mice exhibited a three-fold increase in MCM2 labelling (5.3 ± 0.8%, *p<0.05), indicating increased β-cell replication compared to PFKFB3^WT^ DS mice (1.9 ± 0.04%) and similar to WT controls (7.0 ± 1.3%, Figure 4A-B). Despite the increase in both cell death and β-cell replication in PFKFB3^βKO^ DS mice, β-cell fractional area was comparable in all three groups (Figure 3A). Thus, the increment in β-cell replication in the PFKFB3^βKO^ DS mice appeared to maintain β-cell mass despite the increased cell death and initial loss of β-cells (measured by TUNEL assay) in the absence of PFKFB3.

**Figure 4.**
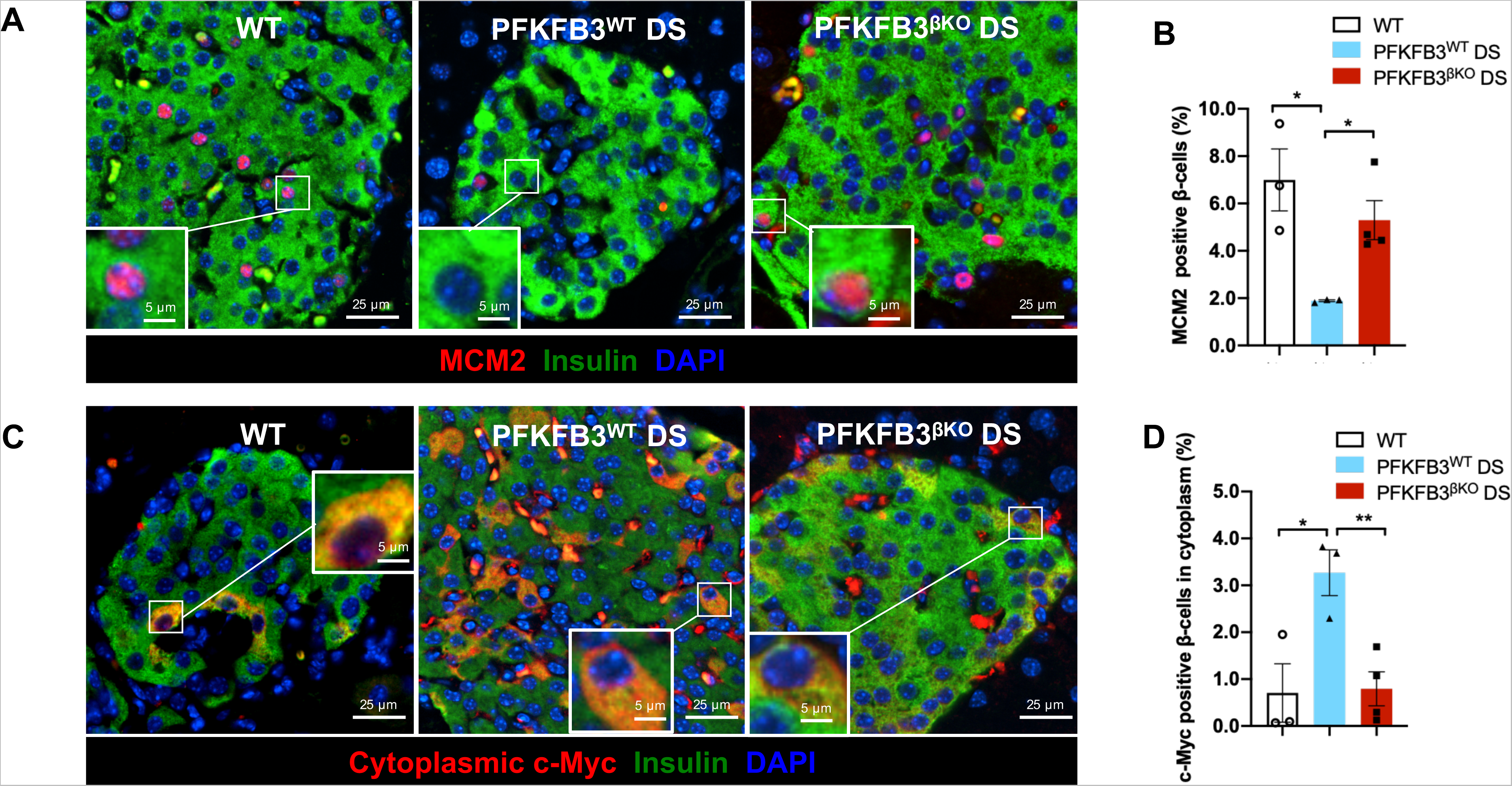
PFKFB3^βKO^ DS mice show increased replication of healthy β-cells compared to PFKFB3^WT^ DS mice. **(A)** Representative immunofluorescence images of islets from WT+HFD, PFKFB3^WT^ DS and PFKFB3^βKO^ DS immunostained for MCM2 (red), insulin (green) and nuclei (blue). (**B**) Quantification of images under A. **(C)** Representative immunofluorescence images of islets from WT+HFD, PFKFB3^WT^ DS and PFKFB3^βKO^ DS immunostained for c-Myc (red), insulin (green) and nuclei (blue). (**D**) Quantification of cytoplasmic c-Myc indicating cells undergoing hIAPP-induced calpain activation (damage) as revealed by immunostaining in C (n=3, n=4 for PFKFB3^βKO^ DS, SEM *p<0.05).

To clarify if replicating β-cells possessed any residual injury, we made use of the specific marker of hIAPP - incurred damage in β-cells – the cytoplasmic accumulation of the calpain-mediated truncation of c-Myc [32]. Our analysis revealed an increase of cytoplasmic c-Myc truncation in PFKFB3^WT^ DS (3.3 ± 0.5%, *p<0.05) but reversal to WT control levels in PFKFB3^βKO^ DS mice (0.8 ± 0.4% and 0.7 ± 0.6%, respectively, Figure 4C-D). These results indicated that healthy β-cells contributed to the increment in replication, probably after increased clearance of β-cells with hIAPP misfolding stress (damaged β-cells).

### Mouse β-cells with residual HIF1α signature resemble a subpopulation of β-cells in humans with T2D

HIF1α immunostaining in the pancreatic sections of PFKFB3^βKO^ DS mice affected about 14% of all β-cells, and although it coincided with reduced misfolding protein stress measured by cytoplasmic c-Myc, this potentially indicated a reason for non-recovered metabolic function in replicating β-cells (Figure 1C-D). To characterize the potential contribution of these HIF1α-positive β-cells to loss of function, we used single-cell RNA sequencing (sc RNA-seq) data from humans with T2D in comparison to nondiabetics available in a public repository [33]. First, we analyzed the quality and validated sc RNA-seq data (Supplementary Figures 4 and 5. Pancreatic cells from healthy and T2D donors were reclustered (umap_cluster) and annotated to the specific cell types based on gene markers such as insulin (INS) for β-cells (umap_celltype, Figure 5A-D). β-cells were then separated into those from healthy and T2D conditions (umap_disease, Figure 5A and Supplementary Figure 6). Nine different cell types (clusters) were identified and the gene expression that delineated their differences are presented in Figure 5C and Supplementary Figures 7 and 8. Using INS as a β-cell marker we identified two clusters of β-cells, cluster 1 and cluster 7, while clusters 2-6, 8 and 9 referred to other pancreatic cell types (Figure 5B, C). Composition in clusters 6 (α-cells), 7 (β-cell subpopulation) and 8 (δ−cells) differed the most between healthy and T2D donors (Supplementary Figure 6). Since HIF1α is mainly regulated in a posttranslational manner, we distinguished β-cells based on the expression or not of lactate dehydrogenase A (LDHA), a HIF1α transcriptional target from aerobic glycolysis (Figure 5D). For each condition, and based on LDHA expression, cells were split into LDHA-positive and LDHA-negative subpopulation and differential expression analysis was performed between the two groups (healthy vs T2D) (Figure 6A-D) [34, 35]. LDHA-positive β-cells completely overlapped with cluster 7 β-cells (Figure 5C,D), and, independent of the disease state, were associated with genes relevant to metabolism, HIF1α signaling, insulin secretion and ghrelin effect on insulin secretion (Table 1 [differentially expressed genes in cluster 1 vs 7], Supplemental Table 1 and Table 2 [differentially expressed genes in LDHA-positive vs negative β-cells] summarizing Enrich*r* analysis). Genes that were upregulated in cluster 7 and LDHA-positive β-cells accounted apart from LDHA in relation to metabolism, for aryl hydrocarbon nuclear translocator 2 (ARNT2, i.e., HIF1β), glucokinase (GK), phosphofructokinase 1 platelet type (PFKFP), pyruvate dehydrogenase kinase 4 (PDK4), genes relevant for insulin secretion such as FBP1 via phosphoenolpyruvate pool, or identity such as glucagon (GCG, pro-α-cell identity) and Aristaless Related Homeobox (ARX, pro-α-cell identity) and INS (lower expression, β-cell identity). LDHA-positive or cluster 7 β-cells showed not only altered (β-cell and pro-α-cell identity) but also an immature phenotype in line with upregulation of genes such as aldehyde dehydrogenase 1A1 (ALDH1A1) [36]. Interestingly, although these cells expressed more pro-apoptotic BID, they expressed at the same time the inhibitor of apoptosis SPOCK3, a key to self-protection of cell subpopulations with lower survival features (Figure 6B,D) [17].

**Figure 5.**
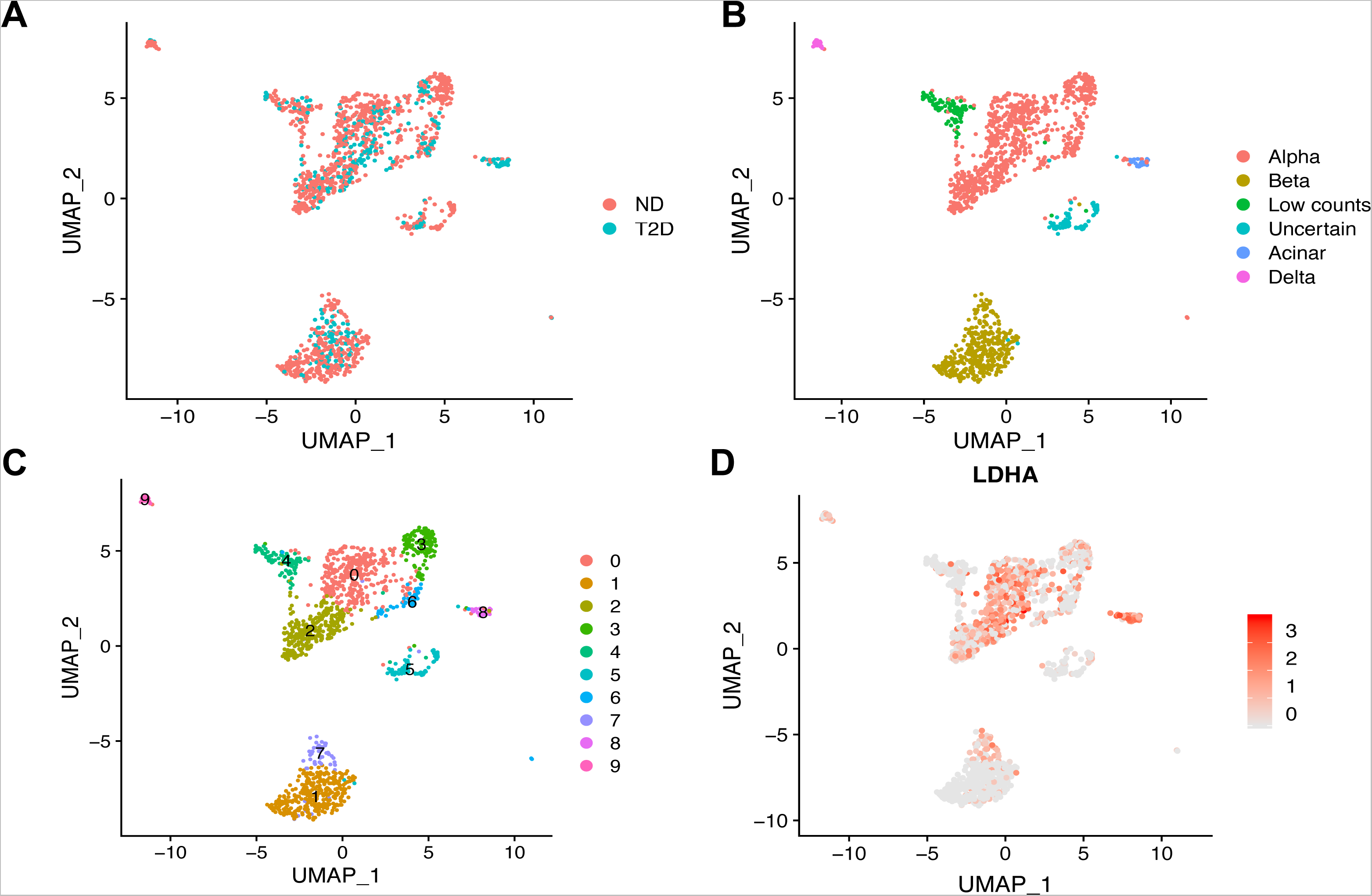
Cluster 7 β-cell subpopulation overlaps with LDHA-positive β-cells with HIF1α signature. (**A**) UMAP plot of cluster distribution of 1,482 pancreatic cells colored by disease conditions (non-diabetics [ND] and T2D). (**B**) UMAP plot of cluster distribution and colored by cell type annotations (α-cells (alpha), β-cells (beta), cells with low counts, cells with non-verified identity, acinar cells and ductal cells). (**C**) UMAP plot colored by unsupervised clusters. (**D**) UMAP plot showing LDHA expression. The color scale represents normalized expression level of the genes. Published sc RNA-Seq in [33] was presented in a quantitative way to compare nine identified pancreatic cell subpopulations in the top differentially expressed genes.

**Figure 6.**
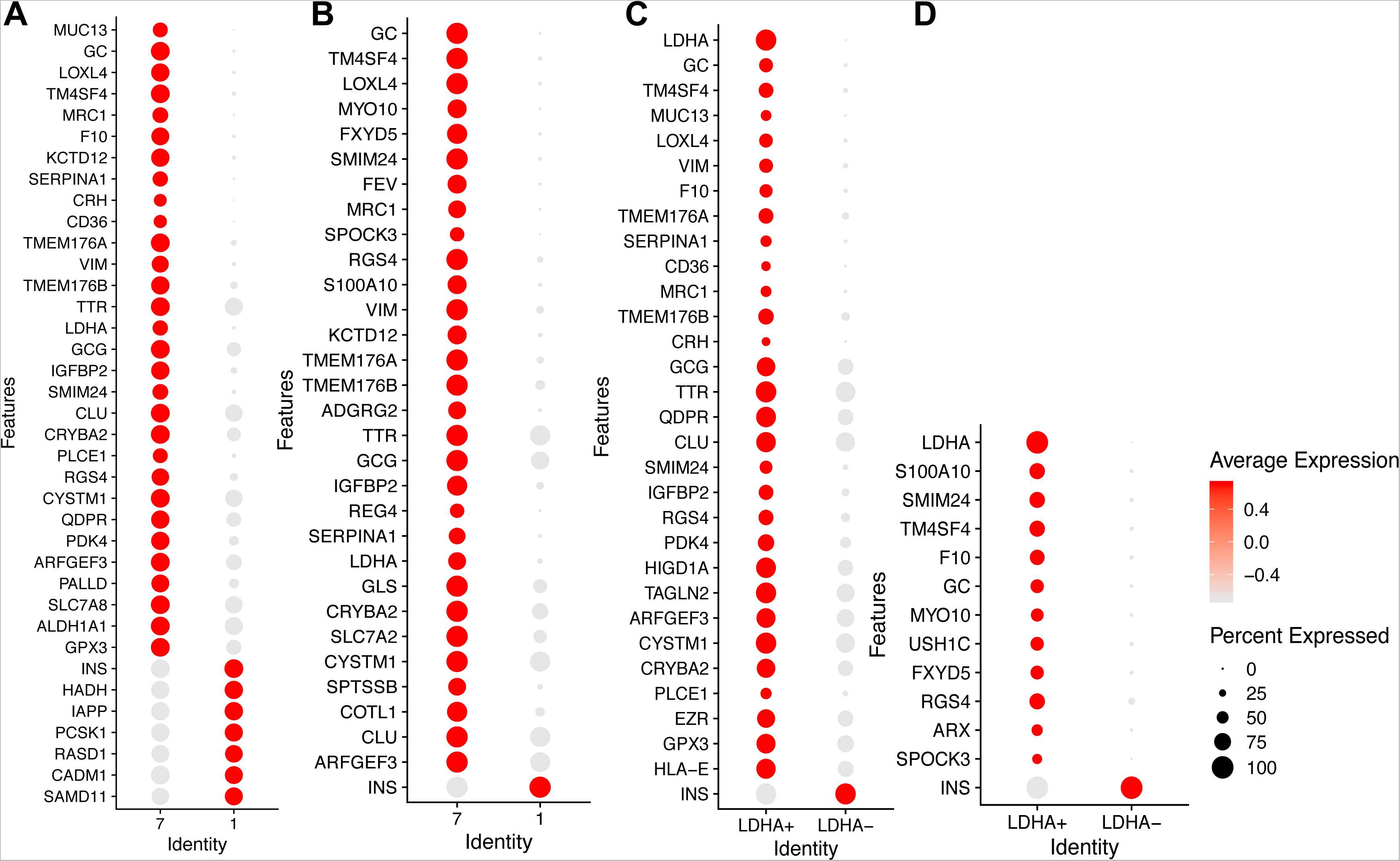
Differential expression analysis comparing LDHA-positive and LDHA-negative β-cells from non-diabetic and T2D donors. (**A**) Dotplot showing the differentially expressed genes between β-cells from cluster 7 and cluster 1 in non-diabetic donors. The size of the dots represents percentage of cells the gene was detected in. The color scale represents the scaled expression of the gene in the two groups. The Wilcoxon Rank-Sum test was performed in the differential expression analysis, and the Benjamini-Hochberg procedure was applied to adjust the false discovery rate. Genes with adjusted p value less than 0.05 were considered significantly differentially expressed. (**B**) Dotplot showing the differentially expressed genes between β-cells from cluster 7 and cluster 1 from T2D donors. (**C**) Dotplot showing the differentially expressed genes between LDHA positive- and LDHA negative β-cells from non-diabetic donors. (**D**) Dotplot showing the differentially expressed genes between LDHA positive- and LDHA negative β-cells from T2D donors.

**Table 1.** β-cell cluster analysis with Enrichr. Top enriched pathways (adjusted p<0.05) in differentially expressed genes in Cluster 1 versus Cluster 7 β-cells in healthy-(top) and T2D donors (bottom) according to curated libraries in Enrich**r**.

| BioPlanet 2019 | Wiki Pathways 2019 | KEGG 2019 human | Elsevier Pathway Collection | Reactome 2016 | TRRUST |
| --- | --- | --- | --- | --- | --- |
| Metabolism | Cori Cycle | Circadian entrainment | L-cell: GCG, PYY, 5-HT release | Peptide hormone metabolism | HIF1α human |
|  | Amino acid metabolism |  |  | Metabolism |  |

T2D
| BioPlanet 2019 | Wiki Pathways 2019 | KEGG 2019 human | Elsevier Pathway Collection | Reactome 2016 | TRRUST |
| --- | --- | --- | --- | --- | --- |
| Ghrelin synthesis and secretion | Coagulation | Coagulation | Ghrelin effect on insulin secretion | Synthesis, secretion and de-acetylation of ghrelin | JunB |
| Diabetes pathways | Cori Cycle | Insulin secretion |  | Peptide hormone metabolism | SP1 |

**Table 2.** LDHA positive-versus negative β-cell analysis with Enrichr. Top enriched pathways (adjusted p<0.05) in differentially expressed genes in LDHA positive versus LDHA negative β-cells in non-diabetic-(top) and T2D donors (bottom) according to curated libraries in Enrich**r**.

| BioPlanet 2019 | Wiki Pathways 2019 | KEGG 2019 human | Elsevier Pathway Collection | Reactome 2016 | TRRUST |
| --- | --- | --- | --- | --- | --- |
| TGFβ | Cori Cycle | Amino acid metabolism | Ghrelin effect on insulin secretion | Peptide hormone metabolism | HIF1α human |
| Metabolism | Amino acid metabolism | Insulin secretion | GCG and PPY regulation |  | SP4 human |

T2D
| BioPlanet 2019 | Wiki Pathways 2019 | KEGG 2019 human | Elsevier Pathway Collection | Reactome 2016 | TRRUST |
| --- | --- | --- | --- | --- | --- |
| Insulin receptor substrate | Cori Cycle | HIF1α signaling | $\beta$ - to α-cell conversion | Coagulation | DLX2 |
|  |  | Insulin secretion | Hyperglycemia and hyperlipidemia | IRS activation | ISL1 |

These results suggested that in humans with or without T2D, a fraction of β-cells (LDHA-positive cells overlapping with cluster 7 β-cells) possess a genetic signature with reduced INS and increased GCG and ARX expression. Insight into overrepresentation by Ingenuity Pathway Analysis (IPA) revealed that the difference between the significantly altered genes in clusters 1 and 7, and between LDHA-positive and negative cells is recapitulated by liver-X/retinoid-X receptor (LXR/RXR) (Table 3) [37, 38], indicating this upstream regulator as a part of the epistatic HIF1α non-canonical metabolic pathway. Moreover, the STRING analysis indicated that while differences between clusters 1 and 7, as well as LDHA-negative and positive β-cells, were well preserved in healthy individuals, these differences were strongly reduced in T2D. These data suggested that in T2D, the clusters of β-cells begin to resemble each other (Supplementary Figure 9A,B and 10A,B). When the β-cell clusters were compared in either T2D or healthy individuals, differentially expressed genes in cluster 1 and LDHA-negative β-cells showed a significant overlap. While a difference was observed in cluster 1 (Supplementary Figure 11A,B), no difference was observed in cluster 7 or LDHA-negative β-cells when health was compared to T2D. Collectively, given that in T2D, clusters 7 and 1 became more similar, and cluster 7 didn’t change, we concluded that cluster 1 underwent changes to become more similar to cluster 7. All differentially expressed genes between cluster 1 and 7 and LDHA-positive versus negative β-cells in ether health or T2D, as well as cluster 1 differences relative to health versus T2D are presented in Supplemental Tables 2 - 7

**Table 3.** Upstream regulators in Cluster 1 vs 7 and LDHA-positive vs negative β-cells. Summary of top upstream regulators identified in Cluster 1 versus Cluster 7 and LDHA-positive versus LDHA-negative β-cells in non-diabetics and T2D according to Ingenuity Pathway Analysis.

| Cluster 1 vs 7<br>in health | Cluster 1 vs 7<br>in T2D | LDHA pos vs neg<br>in health | LDHA pos vs neg<br>in T2D |
| --- | --- | --- | --- |
| LXR/RXR<br>activation | LXR/RXR<br>activation | LXR/RXR<br>activation | FXR/RXR<br>activation |
| ILK signaling | FXR/RXR<br>activation | FXR/RXR<br>activation | Pyruvate to lactate |
| Cytotoxic T cell<br>mediated apoptosis | Macro-pinocytosis | GPCR-mediated<br>nutrient sensing | Extrinsic pro-<br>thrombin activation |
| Opioid signaling | Insulin secretion | Neuropathic pain<br>signaling | G protein signaling |
| Dopamine receptor<br>signaling | Production of NOS<br>and ROS in<br>macrophages | Super-pathway of<br>citruline<br>metabolism | Coagulation |

### Bihormonal cells in diabetic mice resemble human β-cells with HIF1α signature

To find a complementary β-cell population (low INS and co-expression of GCG and ARX) to cluster 7 or LDHA-positive cells, we double stained pancreatic sections from our experimental mice with specific insulin and glucagon antibodies.

Diabetogenic stress increased twice the number of double-positive (bihormonal) cells in PFKFB3^WT^ DS compared to WT controls (5.7 ± 2.8% relative to 2.7 ± 1.8%, respectively, Figure 7A). PFKFB3 knockout led to a reduction of cells with concomitant insulin and glucagon immunopositivity when compared to PFKFB3^WT^ DS mice (0.8 ± 0.3% relative to 5.7 ± 2.8%, Figure 7A). Not only was the fraction of bihormonal cells abolished in PFKFB3^βKO^ DS mice, but also it correlated with a significant increase in the β-cells (*p<0.05, Figure 7B). This indicated that PFKFB3 disruption led to specific culling of β- cells with double insulin and glucagon identity and/or the replication was stimulated from β-cells. The opposite was true for α-cells relative to all α-, β- and bihormonal cells together (Figure 7C). This ratio reached control levels and was reduced in PFKFB3^βKO^ DS compared to PFKFB3^WT^ DS mice (*p<0.05). Of note, the use of a high-fat diet versus a chow diet led to a decrease in α-cell count in line with α-cell hypotrophy reported in obesity [39]. In addition, the bihormonal cells were present in our experimental mice exposed to the high-fat diet and were not detected in WT^chow^ controls or hIAPP^+/+^ (hTG) on a chow diet used as negative controls, neither in prediabetic nor in diabetic mice (Figure 7D-E).

**Figure 7.**
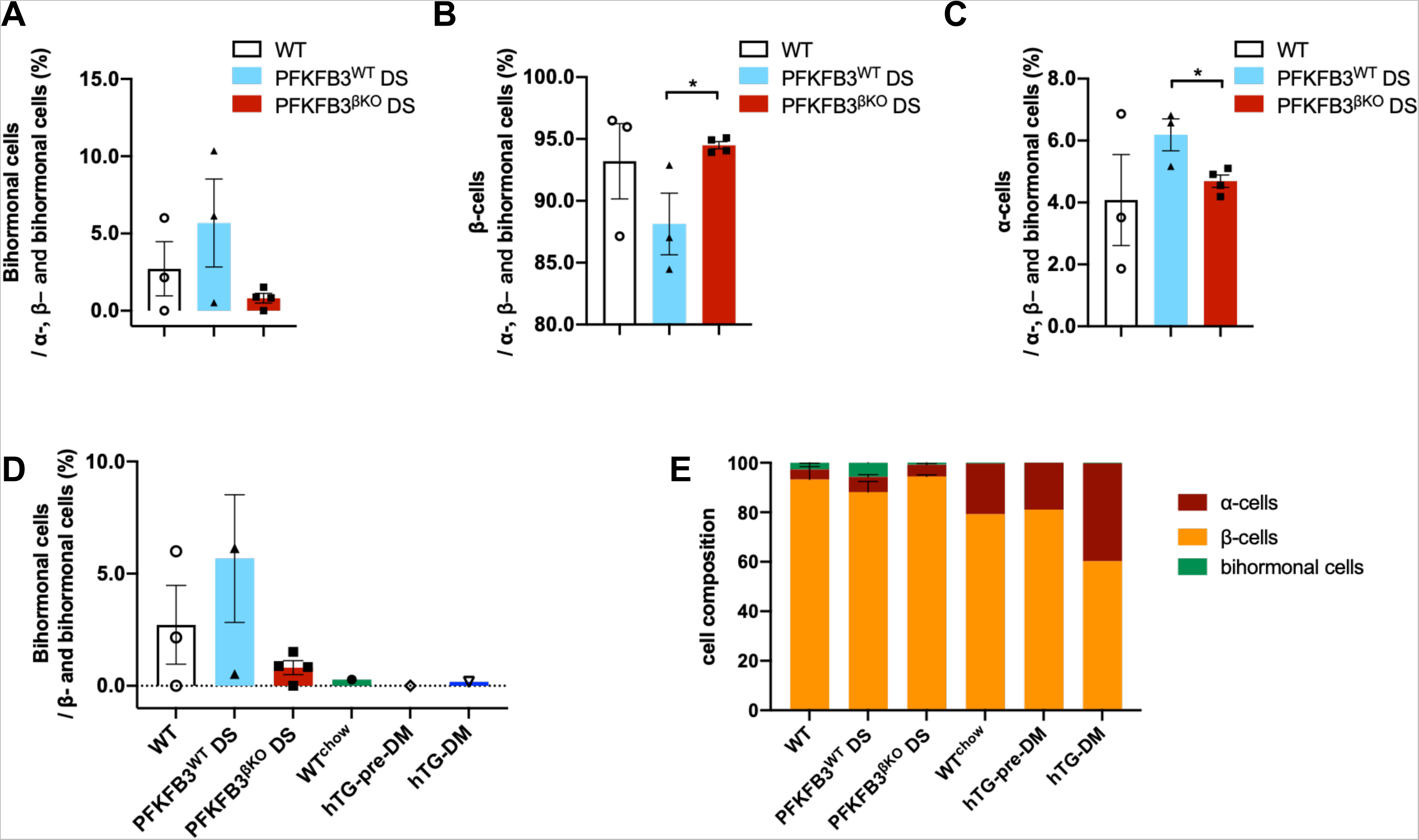
PFKFB3^βKO^ DS mice show a depletion of double-insulin and glucagon positive (bihormonal) cells. (**A**) Quantification of the ratio between bihormonal (insulin- and glucagon positive) cells relative to all α-, β- and bihormonal cells together (%). (**B**) Quantification of the ratio between β-cells relative to all α-, β- and bihormonal cells together (%). (**C**) Quantification of the ratio between α-cells relative to all α-, β- and bihormonal cells together (%). (**D**) Quantification of the ratio between bihormonal cells relative to all insulin positive cells (β- and bihormonal cells together) (%) in indicated experimental groups. WT and homozygous hIAPP^+/+^mice on chow diet with prediabetes (pre-DM) and diabetes (DM) (WT^chow^, hTG-preDM and hTG-DM) were used for comparison to the study experimental groups. (**E**) Cell composition of β-, α-cells and bihormonal cells in indicated experimental groups (n=3, n=4 for PFKFB3^βKO^ DS, SEM *p<0.05).

PFKFB3 knockout leading to depletion of cells with double insulin and glucagon identity indicated that those cells depend on PFKFB3-mediated survival or that the double identity (pro α and pro β-cell identity) represented an adaptation via PFKFB3.

## DISCUSSION

In this study, we demonstrate that the specific β-cell disruption of the *Pfkfb3* gene in adult mice under high diabetogenic stress leads to partial islet regeneration. This is achieved via culling of damaged and bihormonal (insulin- and glucagon-positive) cells and replication of remnant healthy β-cells.

Culling probably affects a substantial number of β-cells since previously we reported that 30-50% of all β-cells depend on high PFKFB3 expression to sustain the injury in T2D [13]. PFKFB3 implication in remodeled metabolism explains its impact on survival [13], but also poses a question related to the preservation of β-cell function. Metabolic remodeling by HIF1α-PFKFB3 pathway in misfolded protein stress (hIAPP) recapitulates the consequences of HIF1α expression after conditional inactivation of von Hippel Lindau gene (*Vhl*) [13, 40]. In the presence or absence of diabetogenic stress, HIF1α activation led to diminished glucose-stimulated changes in cytoplasmic Ca^2+^ concentrations, electric activity and insulin secretion culminating in impaired systemic glucose tolerance [13, 40].

HIF1α was expressed in a significant number of β-cells from PFKFB3^βKO^ DS mice (∼ 14%). This indicated that in this mouse model of diabetes, pertained HIF1α response is independent of PFKFB3. HIF1α expression levels in PFKFB3^βKO^ DS and PFKFB3^WT^ DS mice (14% and 18%, respectively) paralleled the glucose intolerance and the lower plasma insulin- and C-peptide levels in comparison to WT controls (p<0.05).

To investigate the role of HIF1α in the molecular basis for β-cell dysfunction, we analyzed scRNA Seq data from humans with obese-T2D and nondiabetics [33]. We made use of the distinction of LDHA-positive versus negative β-cells since LDHA is a bona fide target, serving as a substitute marker for HIF1α. HIF1α is regulated mainly posttranslationally with no changes in the transcript levels. LDHA-positive β-cells overlapped with cluster 7 β-cell subpopulations and were represented by HIF1α− (ARNT2, GK, PFKFP, PDK4) and bihormonal signature (α− and β-cell identity) (GCG, ARX and INS) and some markers of immaturity (ALDH1A1). Bihormonal cells in our mouse model resembled LDHA-positive or cluster 7 β-cells and could originate from β-cells subjected to dedifferentiation or transdifferentiation [41].

Ingenuity Pathway Analysis (IPA) used for comparison between LDHA-positive (cluster 7) and LDHA-negative (cluster 1) β-cells in both non-diabetics and T2D, indicated the existence of a master upstream regulator, the liver-X receptor, a type of retinoid receptor (LXR/RXR) [42], potentially a part of the epistatic HIF1α metabolic pathway. Activation of this receptor may independently increase aerobic glycolysis in response to high fat diet via transcriptional upregulation of hexokinases 1 and 3 (HK1 and 3) and SLC2A1 (GLUT1)) and interplay with HIF1α [37, 38].

Interestingly, the bihormonal cell subpopulation was present in the obese nondiabetics and in our WT controls under HFD, indicating that the formation of bihormonal cells may well be an adaptive response to increased lipogenesis and high-fat diet. This is in line with previous discussed studies suggesting that altered identities of β-cells including bihormonal cells likely serve as a compensatory response to enhance function/expand cell numbers such as in obesity or a cell protection from ongoing stress [43, 44].

As such, bihormonal cells were not detected in WT controls or hIAPP^+/+^ in the absence of HFD (used as negative controls), neither in prediabetic nor in diabetic mice. In both non-diabetics and T2D, the difference between bihormonal β-cell cluster 7 (LDHA-positive β- cells) and cluster 1 implicated LXR/RXR. Previous reports indicated that, unlike acute activation that is an adaptive response to increased demand on insulin secretion, the chronic activation of LXR may contribute to β-cell dysfunction by the accumulation of free fatty acids and triglycerides [42].

In addition, the STRING analysis indicated that, while differences between clusters 1 and 7 as well as LDHA-negative and positive β-cells were well preserved in non-diabetics, these differences were reduced in T2D. Preserved features of bihormonal cells can be relevant in the context of cell fitness competition, where differences could constitute a basis for bihormonal cells’ recognition and homeostatic control.

Cell fitness competition is an important extrinsic cell quality control based on the distinction of cell population with inferior versus cell population with superior fitness (survival) characteristics. This distinction is key in triggering selective culling of the cell population with inferior fitness characteristics (“losers”) and the expansion of the cell population with superior fitness characteristics (“winners”). Replacement of the “losers” with the “winners” allows for the maintenance of homeostatic tissue. By implication, the reversal of cell competitive tissue makeup such as we see in T2D (clusters 1 and 7) may lead to inhibition of cell competition followed by tissue dysfunction over time.

In our mouse model of diabetes, PFKFB3 knockout in adult β-cells led to a strong reduction of damaged β-cells and bihormonal cells and a concomitant increase in healthy β-cell replication. We monitored damaged β-cells by measuring the extent of the calpain (hIAPP)-mediated truncation of the cytoplasmic c-Myc [32]. Calpain was previously reported to directly reflect hIAPP misfolded protein toxicity [45, 46]. In PFKFB3^βKO^ DS mice, cytoplasmic c-Myc was reversed to the barely detectable levels as measured in WT controls, respectively. These results indicated that the increment in replication was contributed to by healthy β-cells and was conceivably facilitated by the excelled loss of β- cells with hIAPP injury. Therefore, our results suggested a cell competition-dependent β- cell regeneration by the culling of damaged β-cells after PFKFB3 knockout (Figure 8). In addition, in PFKFB3^βKO^ DS mice, the increase in β-/α-cell ratio and the reduction in glucagon levels in comparison to PFKFB3^WT^ DS may have accounted for the observed trend of higher insulin sensitivity [47-49].

**Figure 8.**
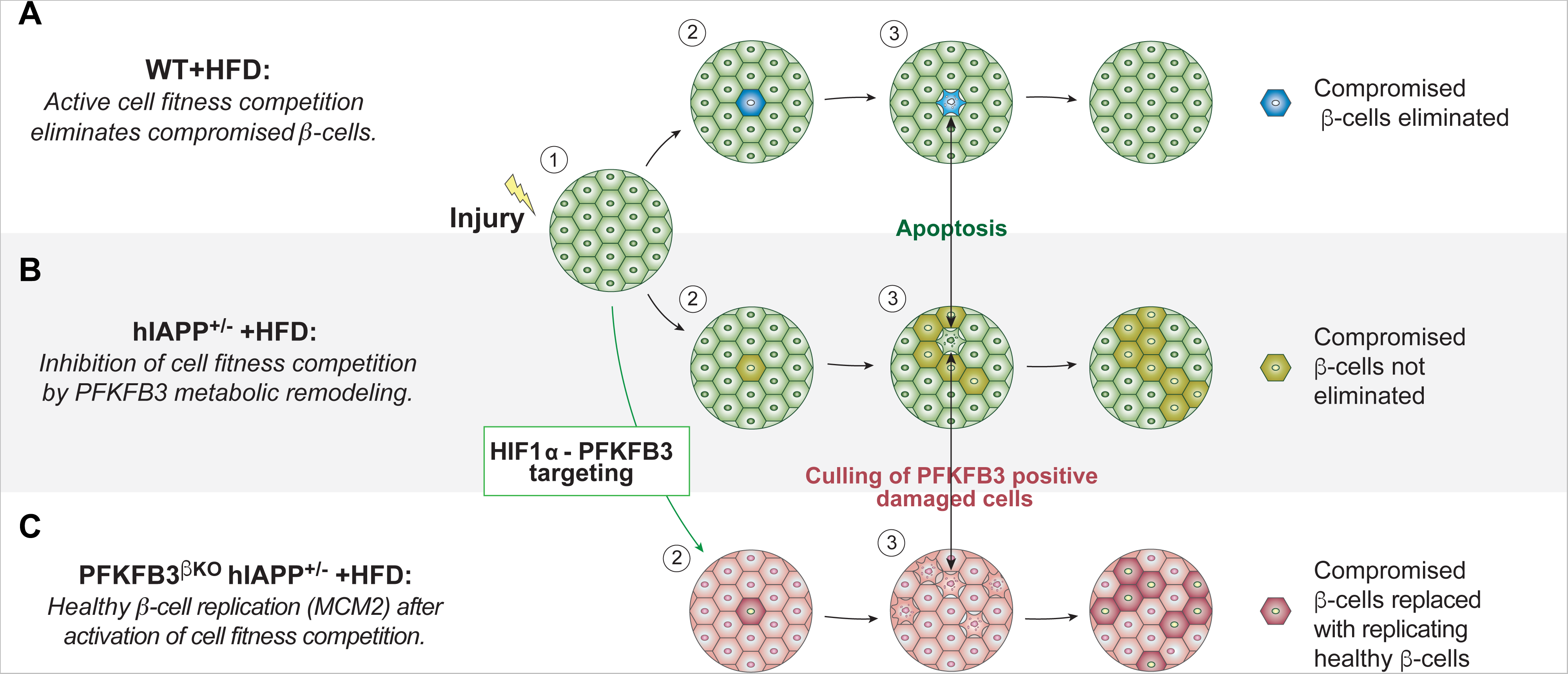
Model of the proposed role of β-cell fitness comparison in β-cell replenishment under stress. An initial insult or injury can change the fitness level of some β-cells within β-cell population and trigger cell competition. (**A**) After metabolic stress such as a high-fat diet (HFD) in non-diabetic or wild-type (WT) mice, suboptimal cells (blue) are eliminated from the tissue by competition with healthy ***β***-cells (green), which replicate to regenerate the lost tissue. (**B**) In a cell competition in which injury is sustained (T2D) or in PFKFB3^WT^ DS mice, damaged β-cells (light green) survive in spite of reduced fitness and cannot be purged from the tissue because of metabolic remodeling by the HIF1α-PFKFB3 pathway. These damaged β-cells may impede healthy β-cell replenishment (replication). (**C**) Under conditions described in (**B**), the targeting of the pro-survival PFKFB3 leads to activation of cell competition and elimination of suboptimal (damaged) β-cells (light pink). Elimination of suboptimal (damaged) β-cells leads to replication of the remaining healthy β-cells (MCM2-positive cells, dark pink).

In conclusion, it appears that the preservation of the β-cell mass and increase in the β-/α-ratio in the PFKFB3^βKO^ DS mice may stem cumulatively from β-cell replication that overcomes the initial loss of damaged β-cells and reduction in bihormonal cells.

While previously differentiation was discussed to restore β-cell mass, here we demonstrate how selective purification of compromised cells via PFKFB3 disruption can serve the same goal [50]. One advantage of the latter strategy would rely on the compensatory hypertrophic/hyperplastic growth after selective elimination of β-cells with lower fitness following the rules of cell competition. For example, in the model of Alzheimer’s disease in Drosophila, activation of cell competition that purged damaged neurons was sufficient to stimulate restoration of cognitive function [50]. It is tempting to propose that PFKFB3 plays a key role in cell competition activation in β-cells under diabetogenic stress. This could be a powerful physiological tool to induce replication in post-mitotic cells under stress such as β-cells in T2D. Dissection of the roles of HIF1α and PFKFB3 in this context indicates that while PFKFB3 targeting can control the β-cell mass dynamic under stress, it is probably a compensatory HIF1α positive response that, linked to LXR/RXR or other pathway, determines the β-cell function under stress. When addressed both in the activation of the physiological cell competition under stress, they might provide a powerful tool to restore a functionally competent β-cell mass in T2D.

## AKNOWLEDGEMENT

This work was supported by funding from the Larry Hillblom Foundation (Start-up Grant #2017-D-002-SUP). J.M. was supported by Department of Endocrinology, Union Hospital of Tongji Medical College Huazhong University of Science and Technology, Wuhan, Hubei, China.

We are very grateful to Madeline Rosenberger and Dr. Tatyana Gurlo for their help advise in the animal breeding and technical issues in the experiments. We thank Dr. Andrea Mattern and the Division of Laboratory Medicine at UCLA for their exceptional support in mice experiments. We also thank Dr. Lulu Chen and Dr. Tianshu Zeng for the critical reading of the manuscript. We thank Dr. Katrien De Bock and Dr. Peter Carmeliet for their kind sharing of the protocol for immunostaining of PFKFB3 in mouse tissue sections.

