## Supplementary Figures, Supplemental data and supplemental tables for "PFKFB3 Depletion Activates β-Cell Replication by Cell Competitive Culling of Compromised β-Cells Under Stress"

#### Supplementary Figure 1

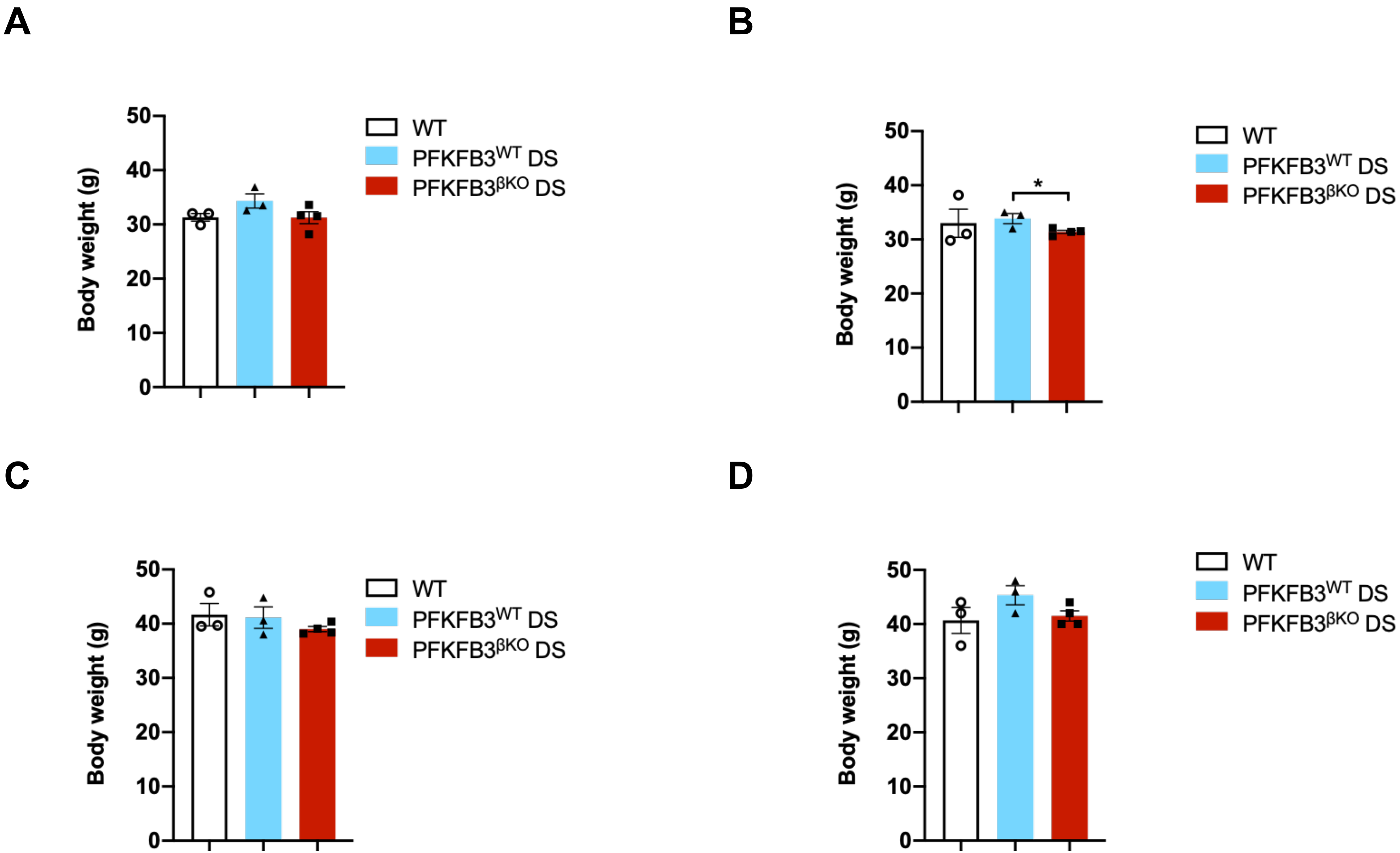

### Supplementary Figure 2

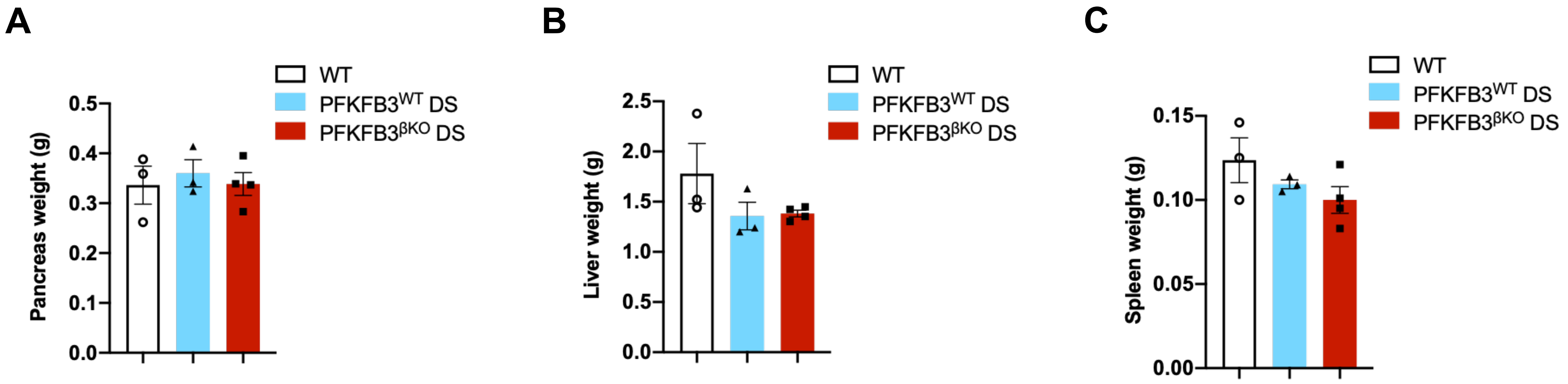

### Supplementary Figure 3

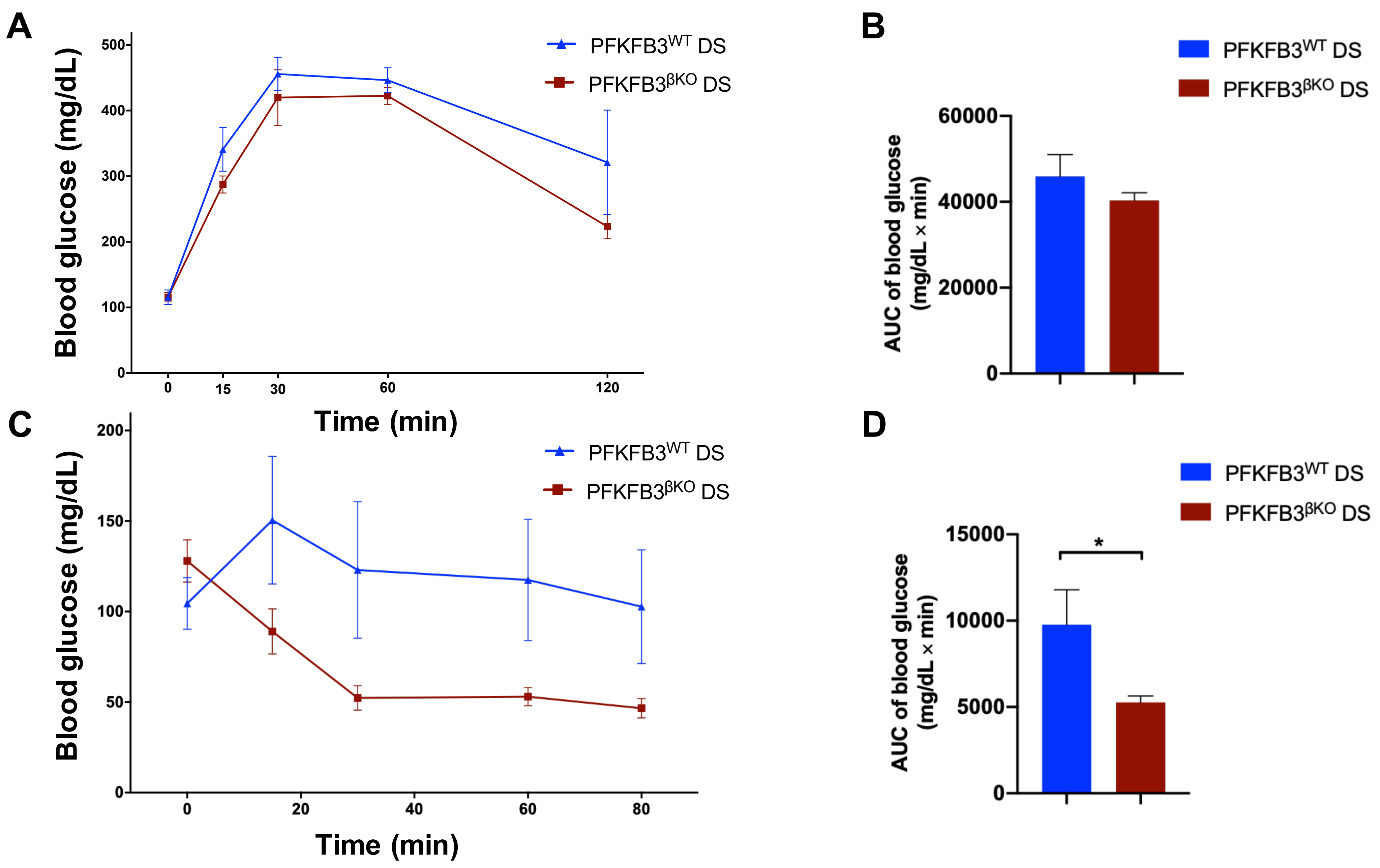

### Supplementary Figure 4

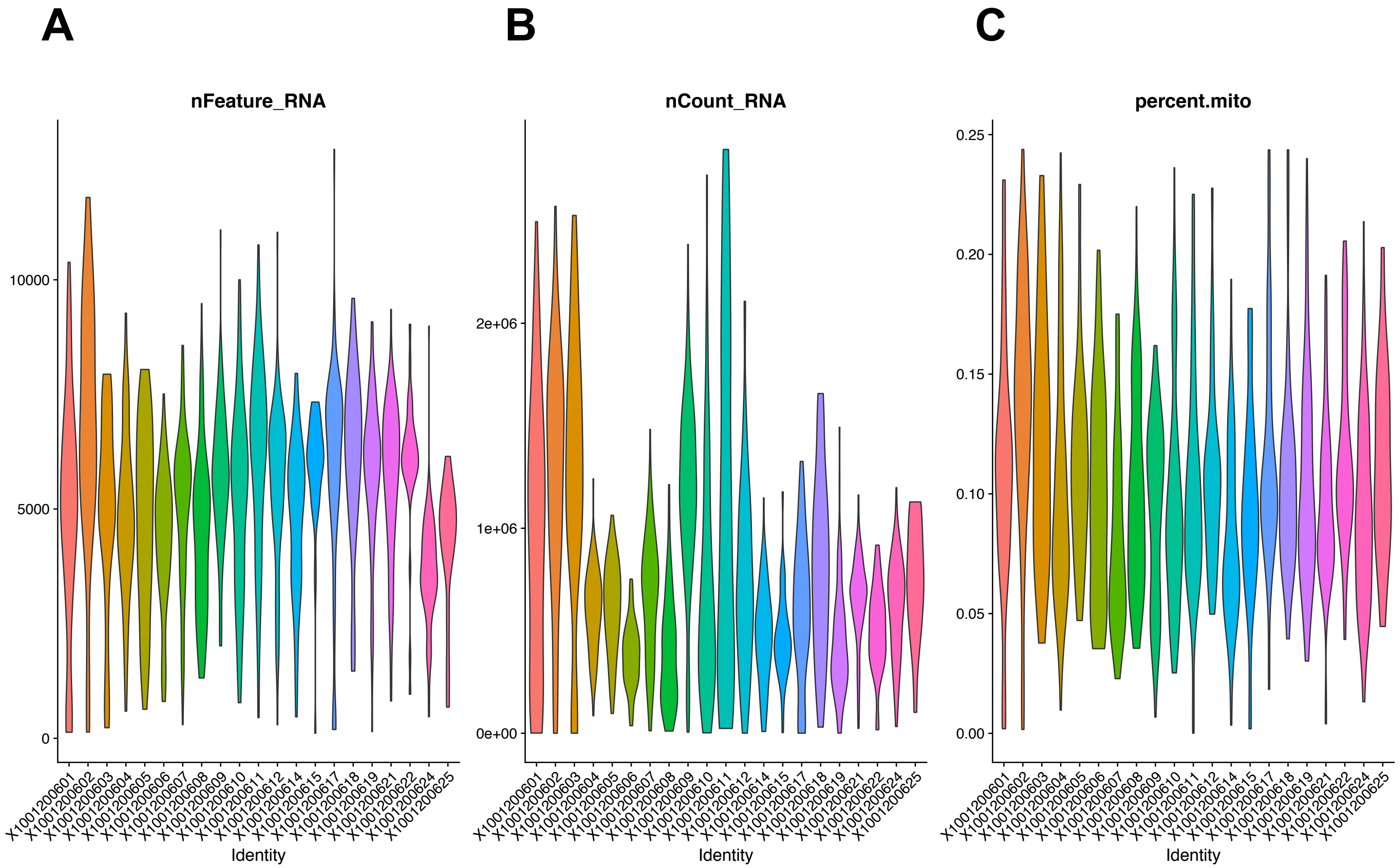

### Supplementary Figure 5

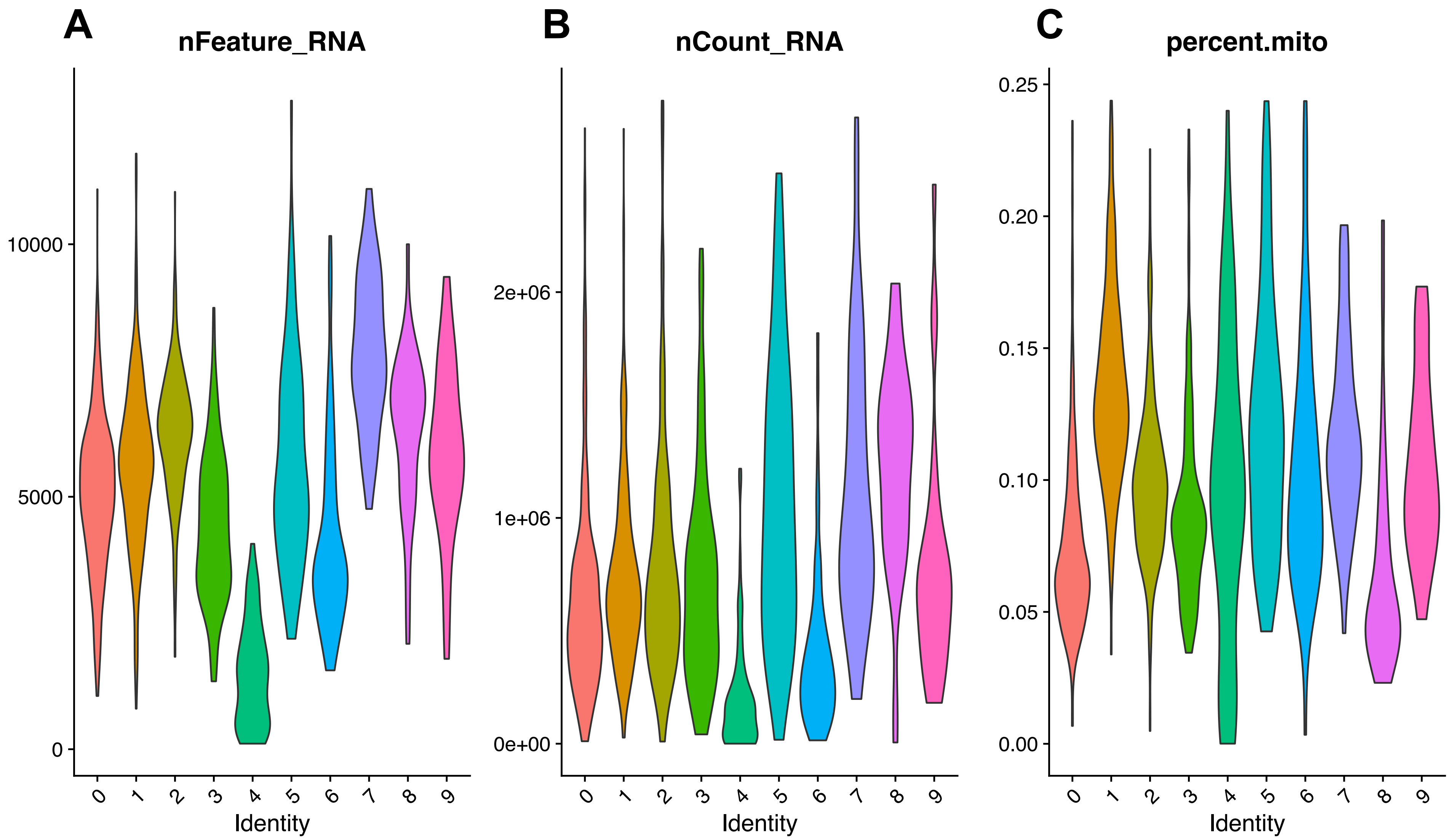

**Supplementary Figure 6**

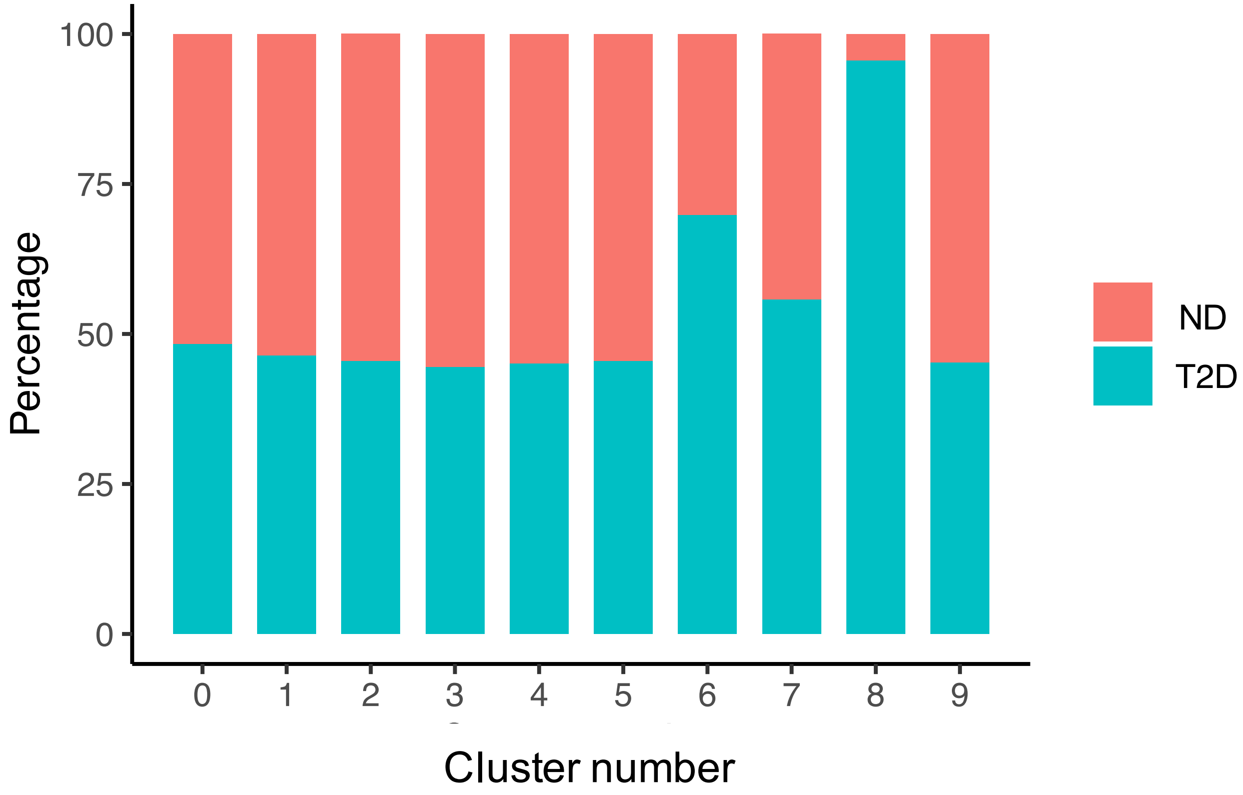

### Supplementary Figure 7

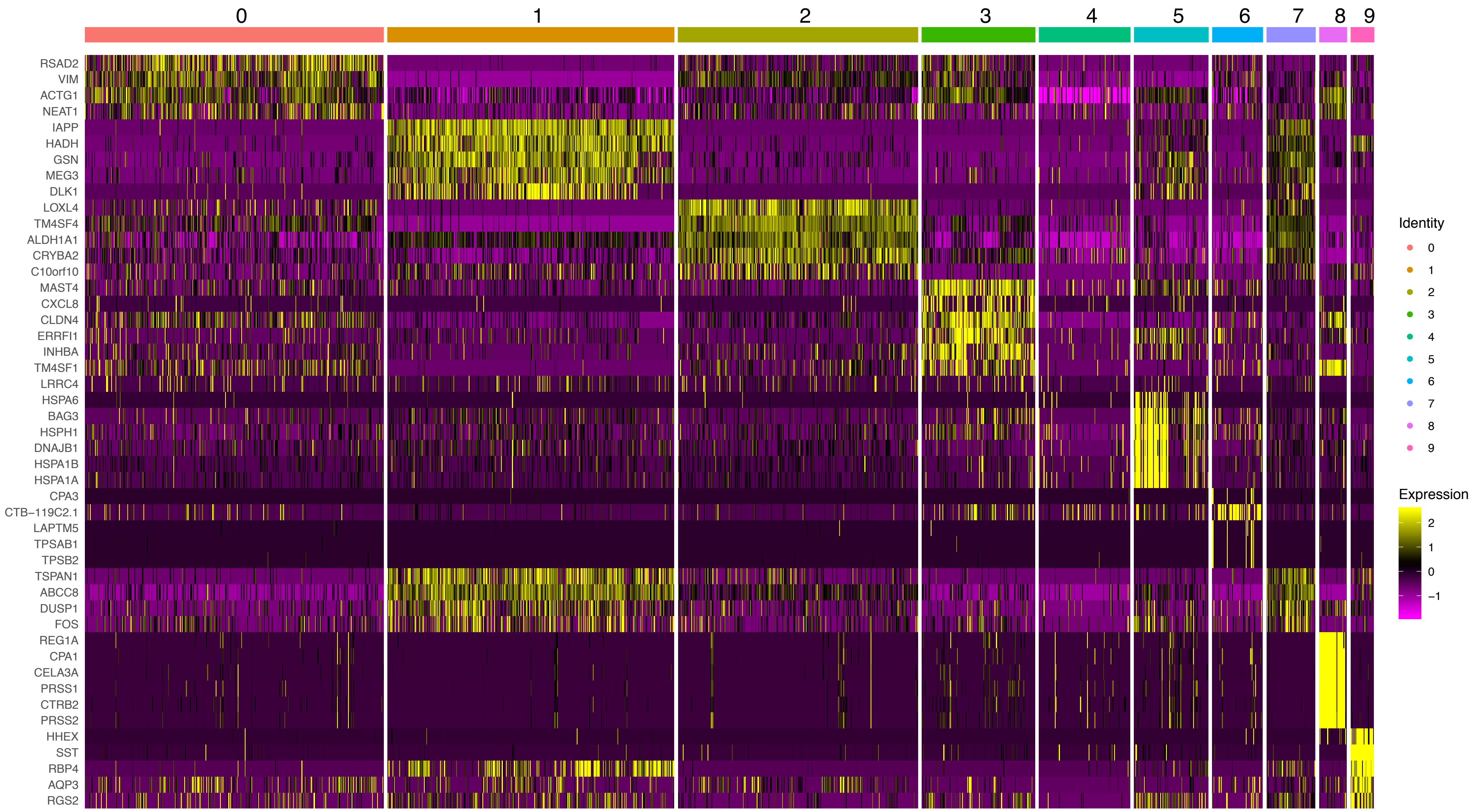

### Supplementary Figure 8

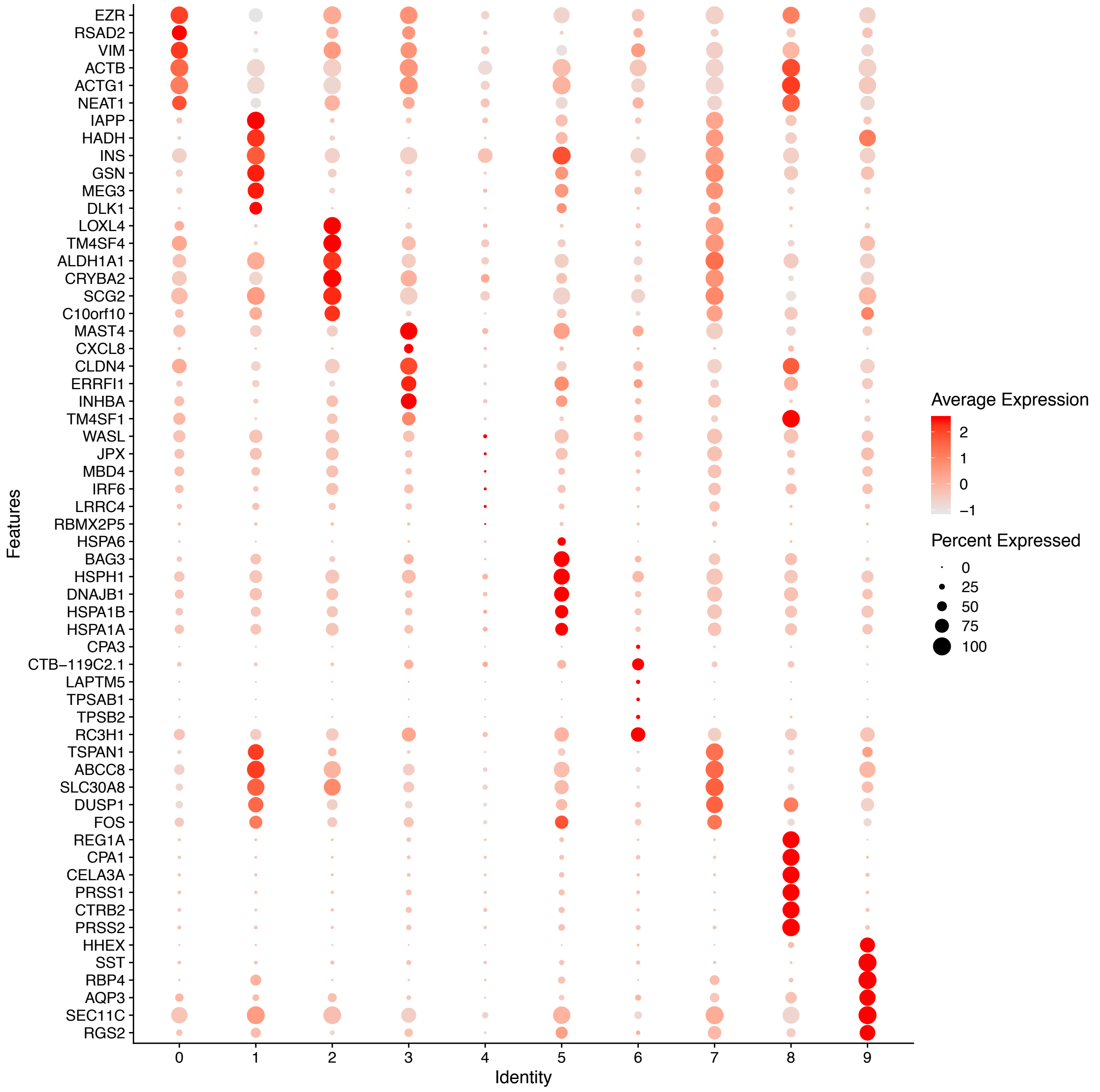

### Supplementary Figure 9

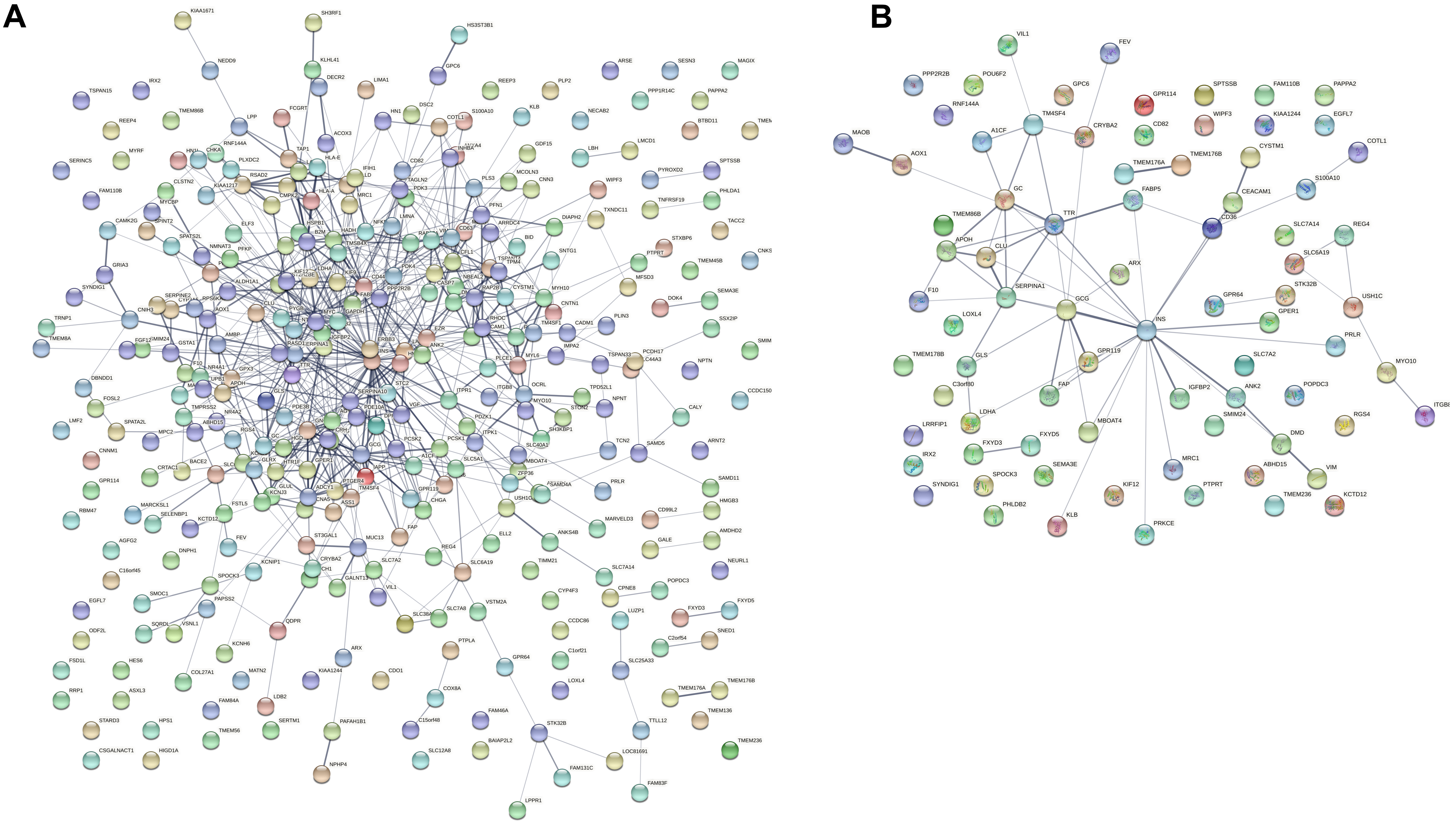

### Supplementary Figure 10

A

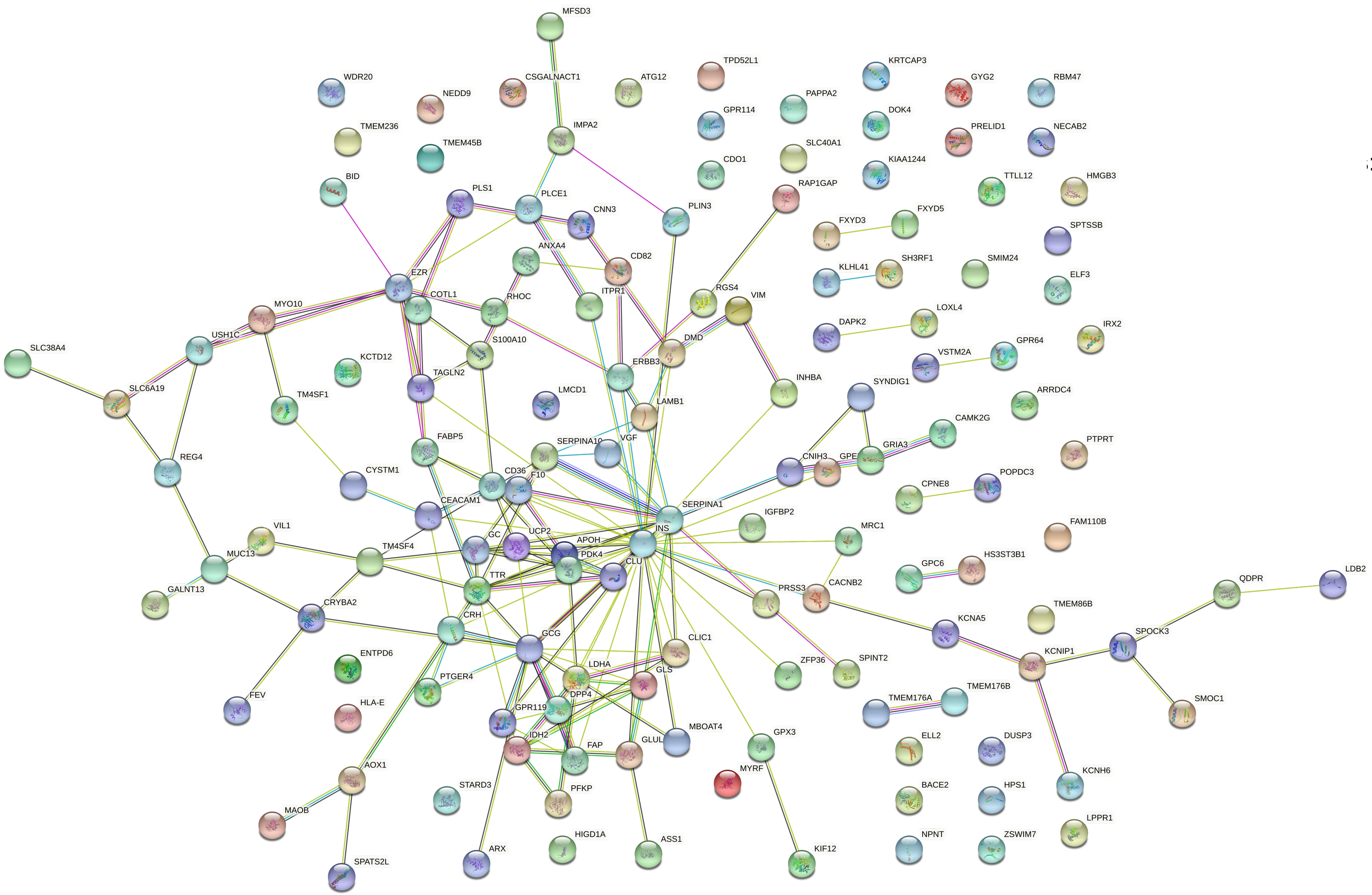


B

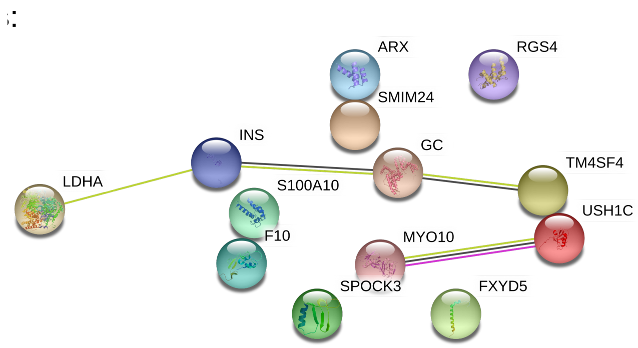

**Supplementary Figure 11**

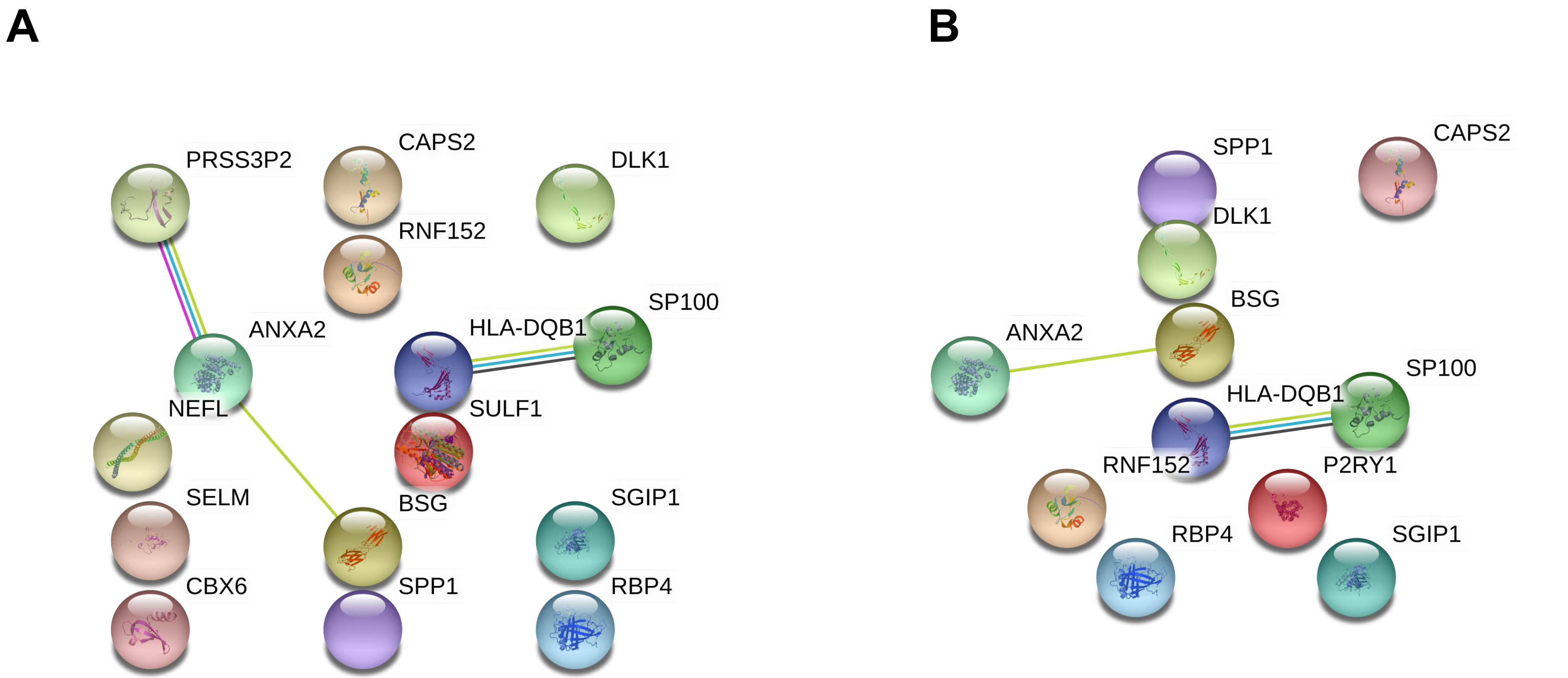

### Supplemental Data

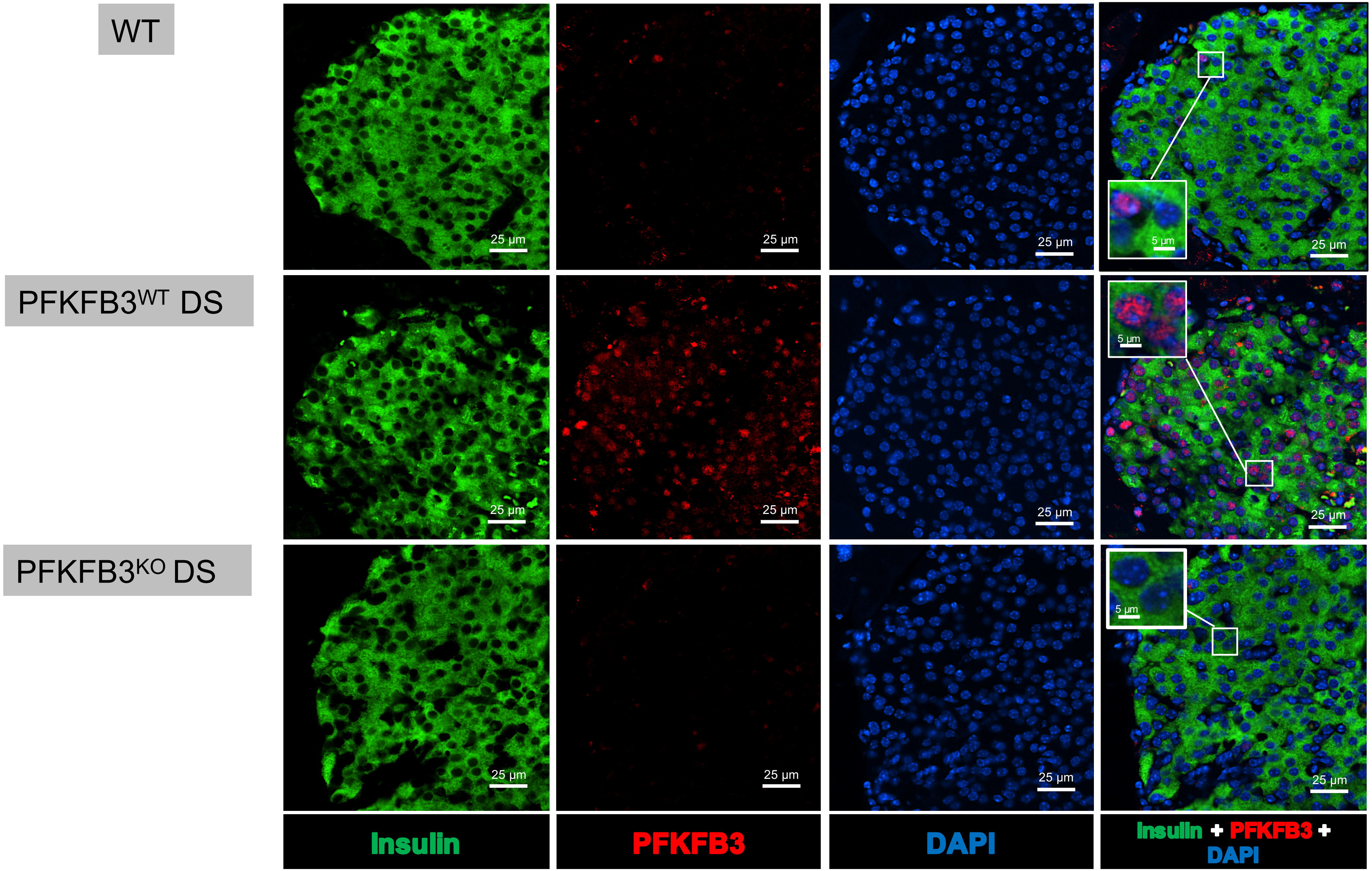

### Supplemental Data

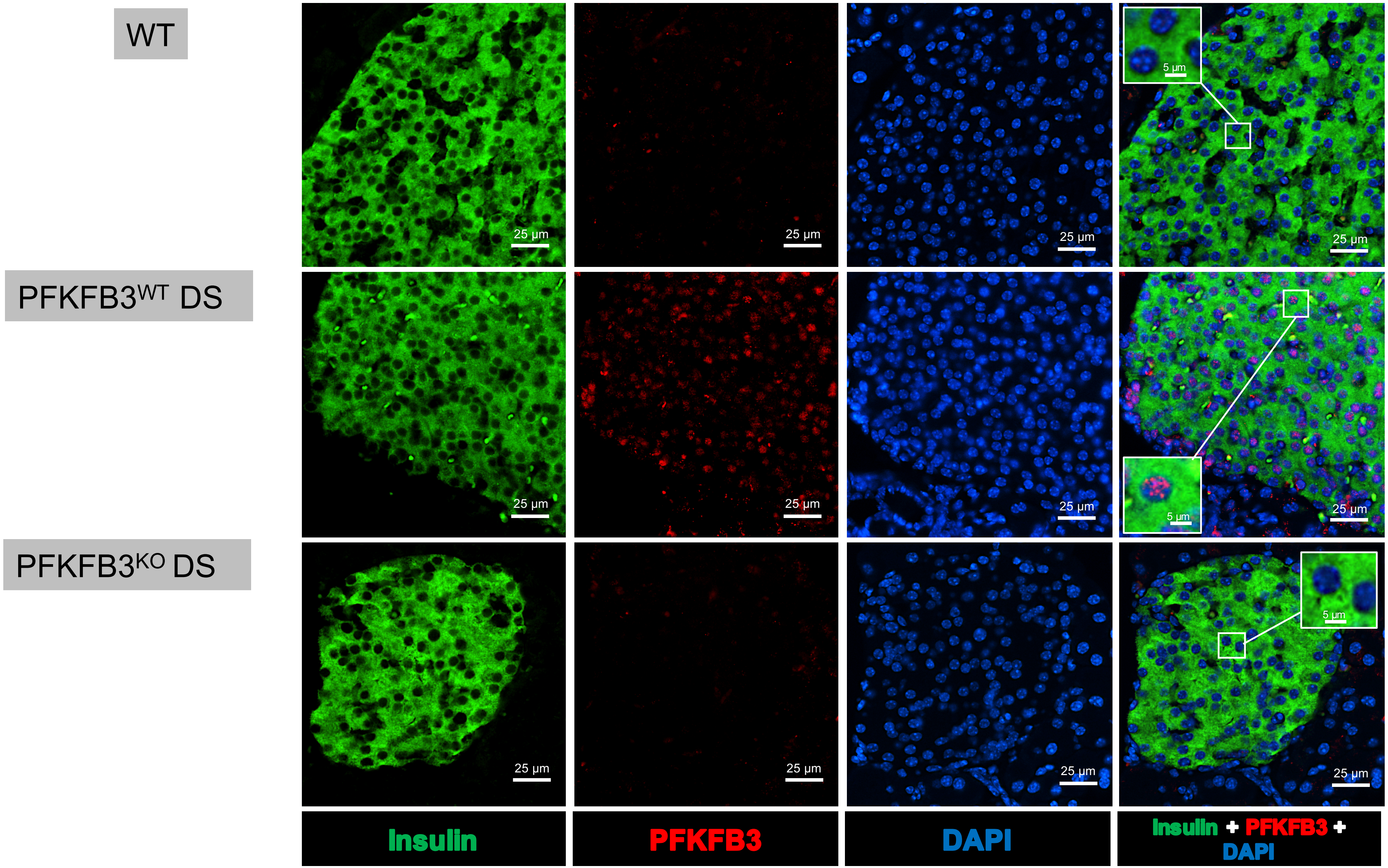

### Supplemental Data

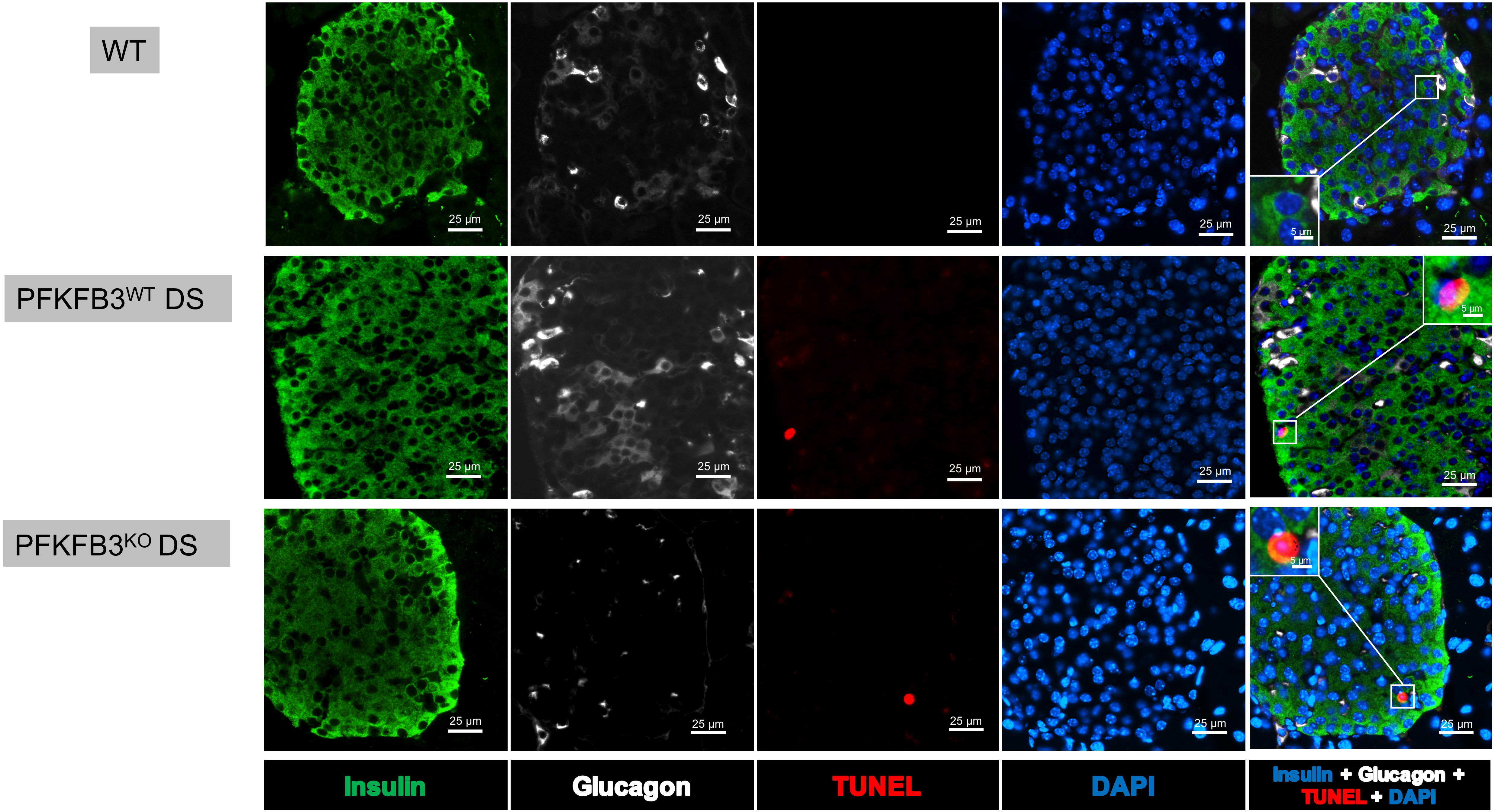

### Supplemental Data

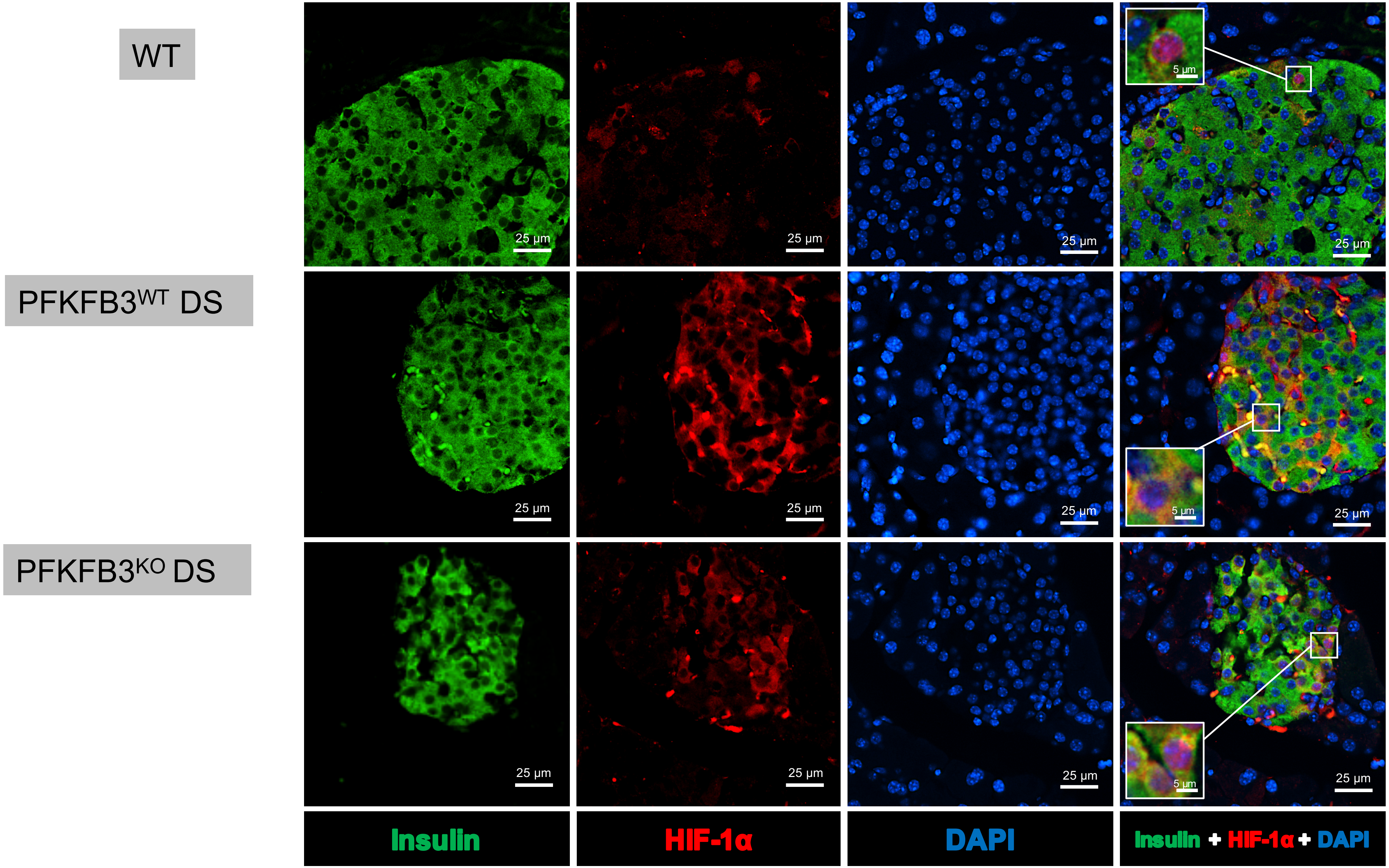

### Supplemental Data

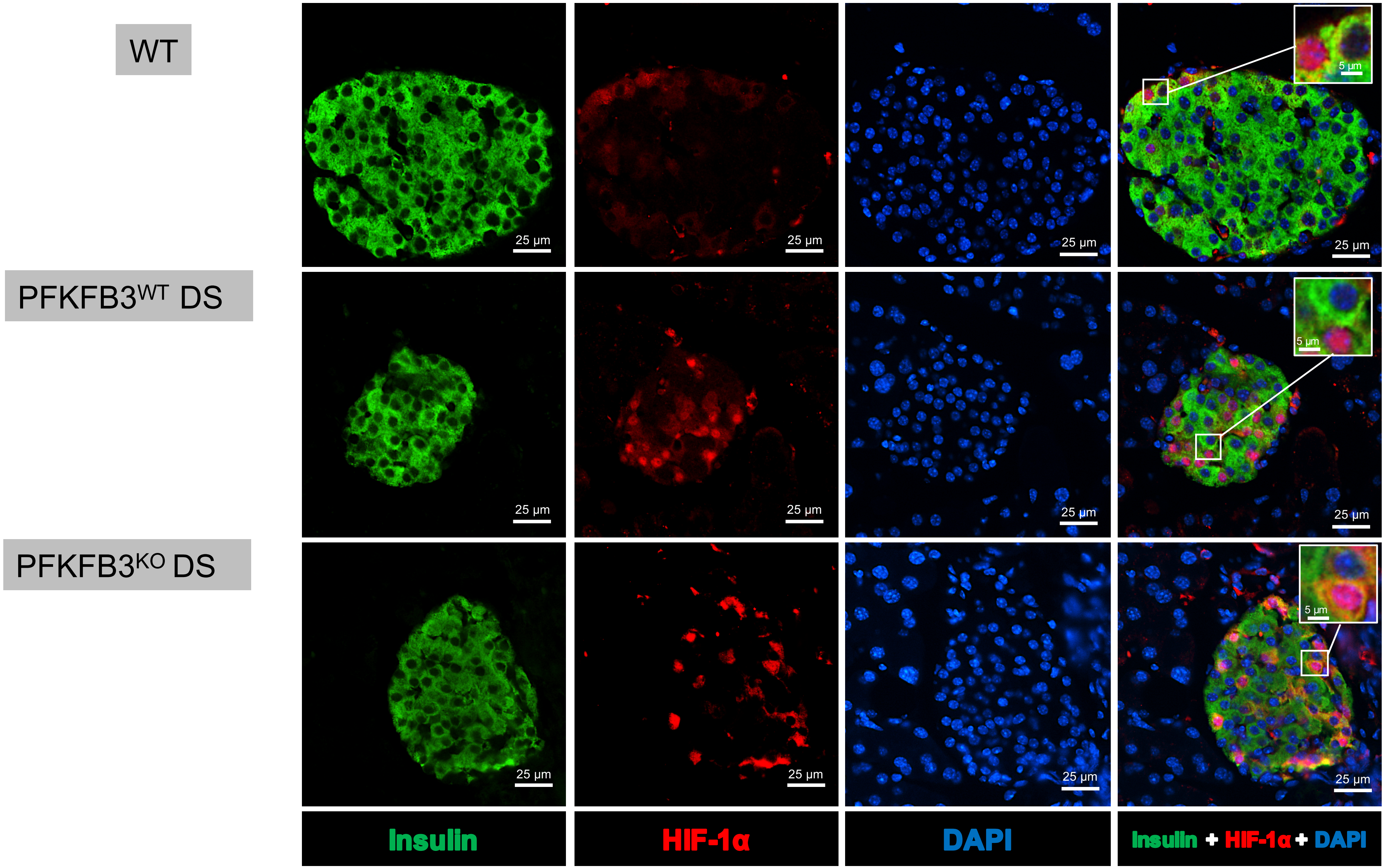

### Supplemental Data

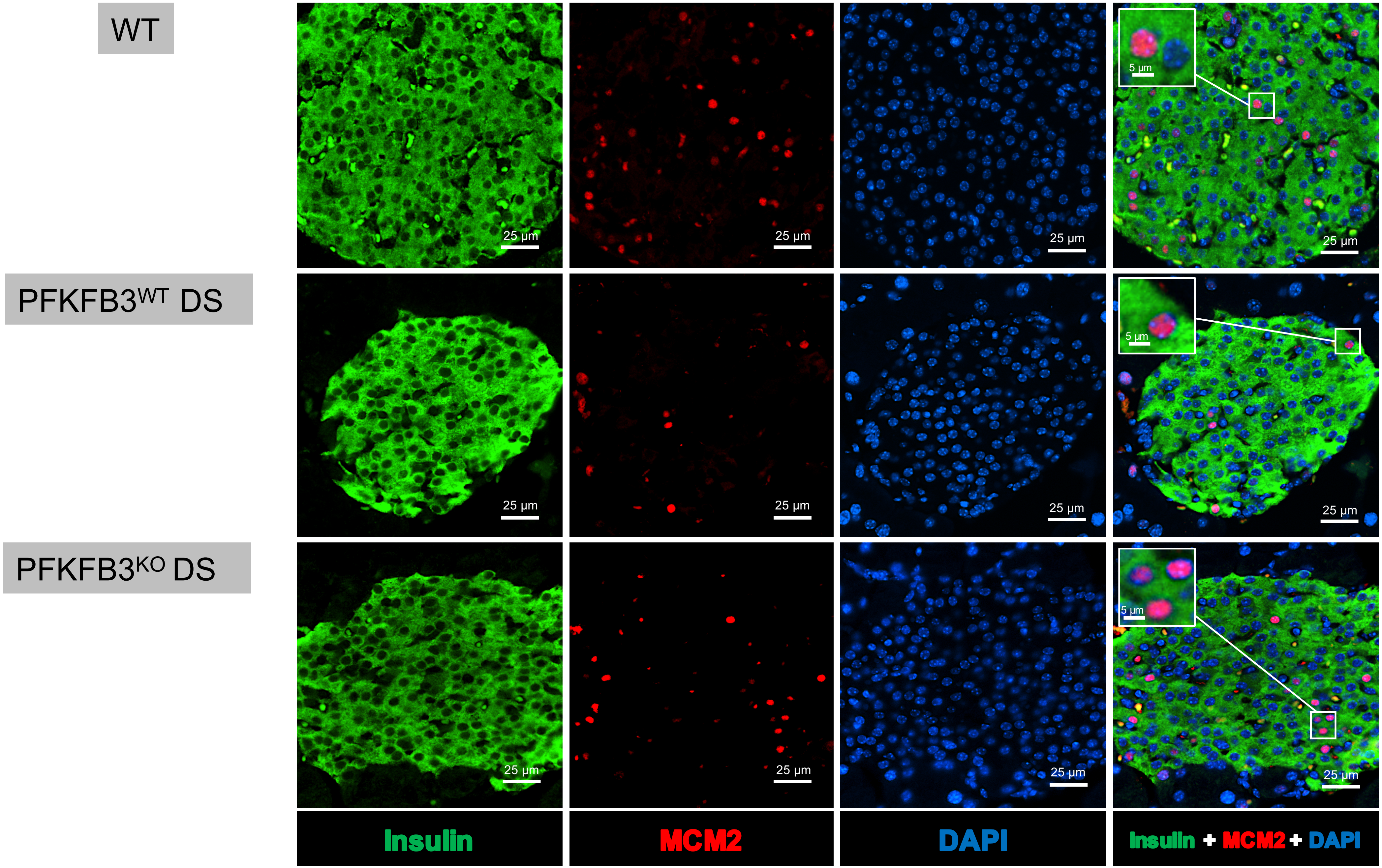

### Supplemental Data

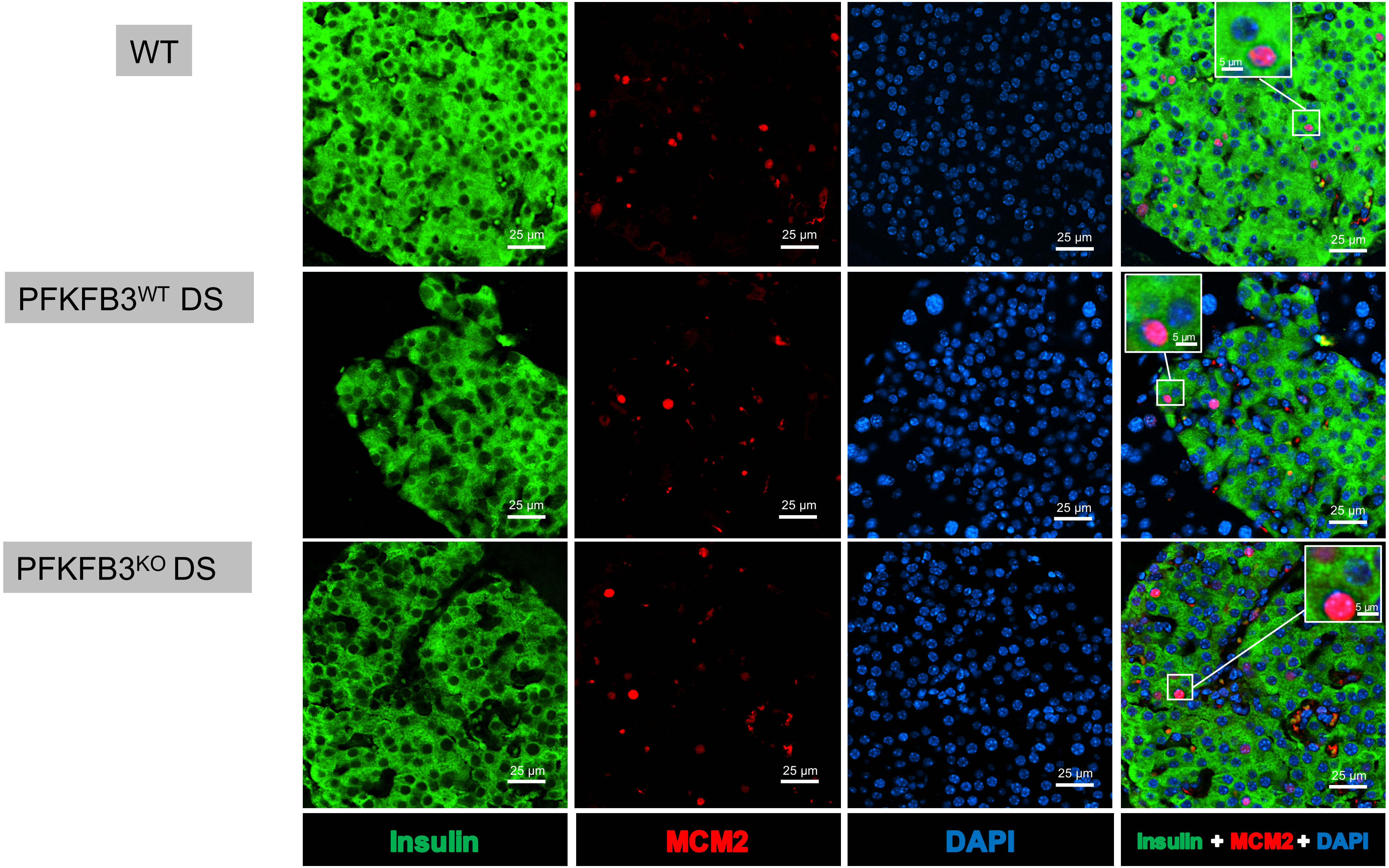

### Supplemental Data

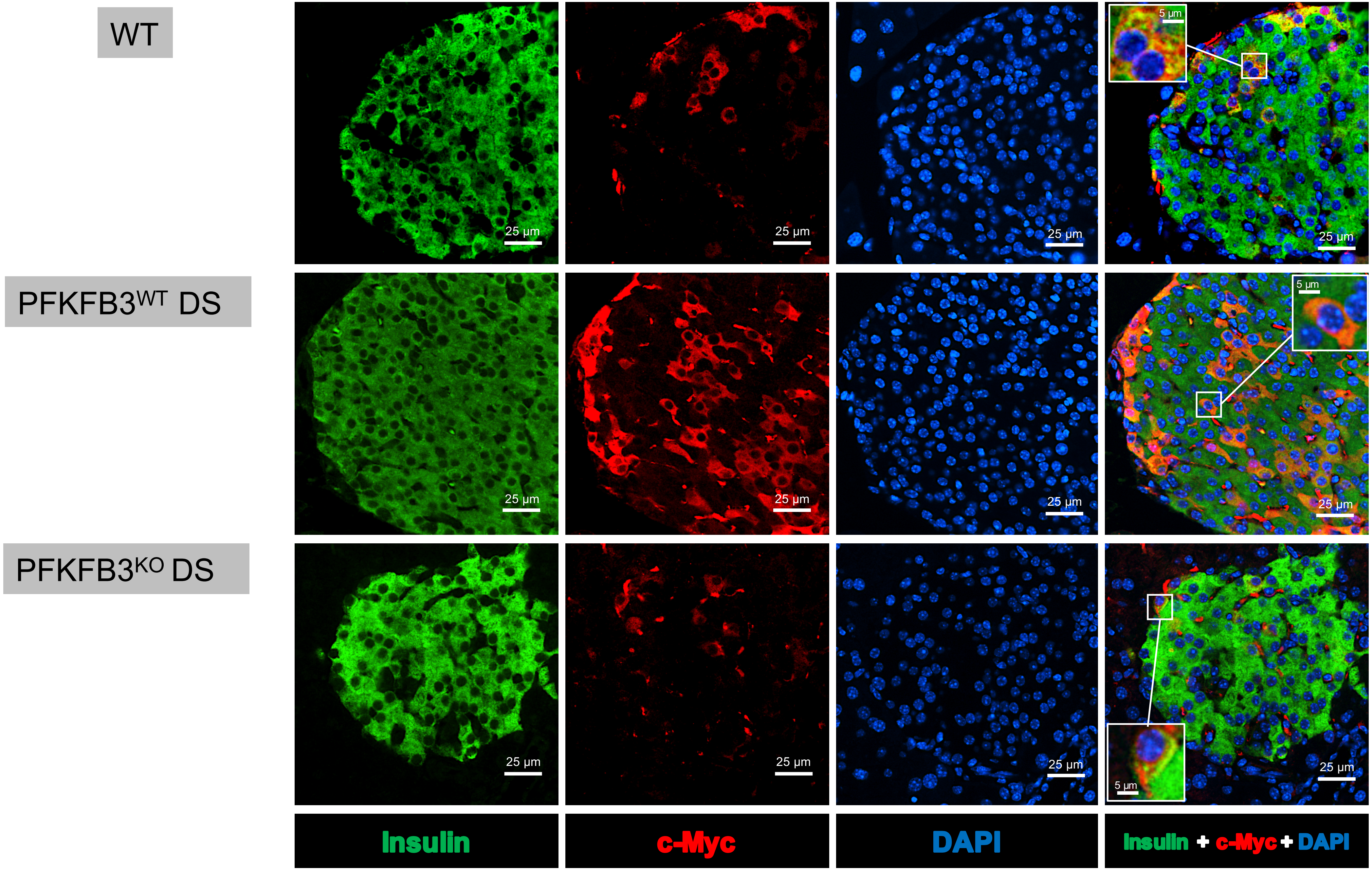

### Supplemental Data

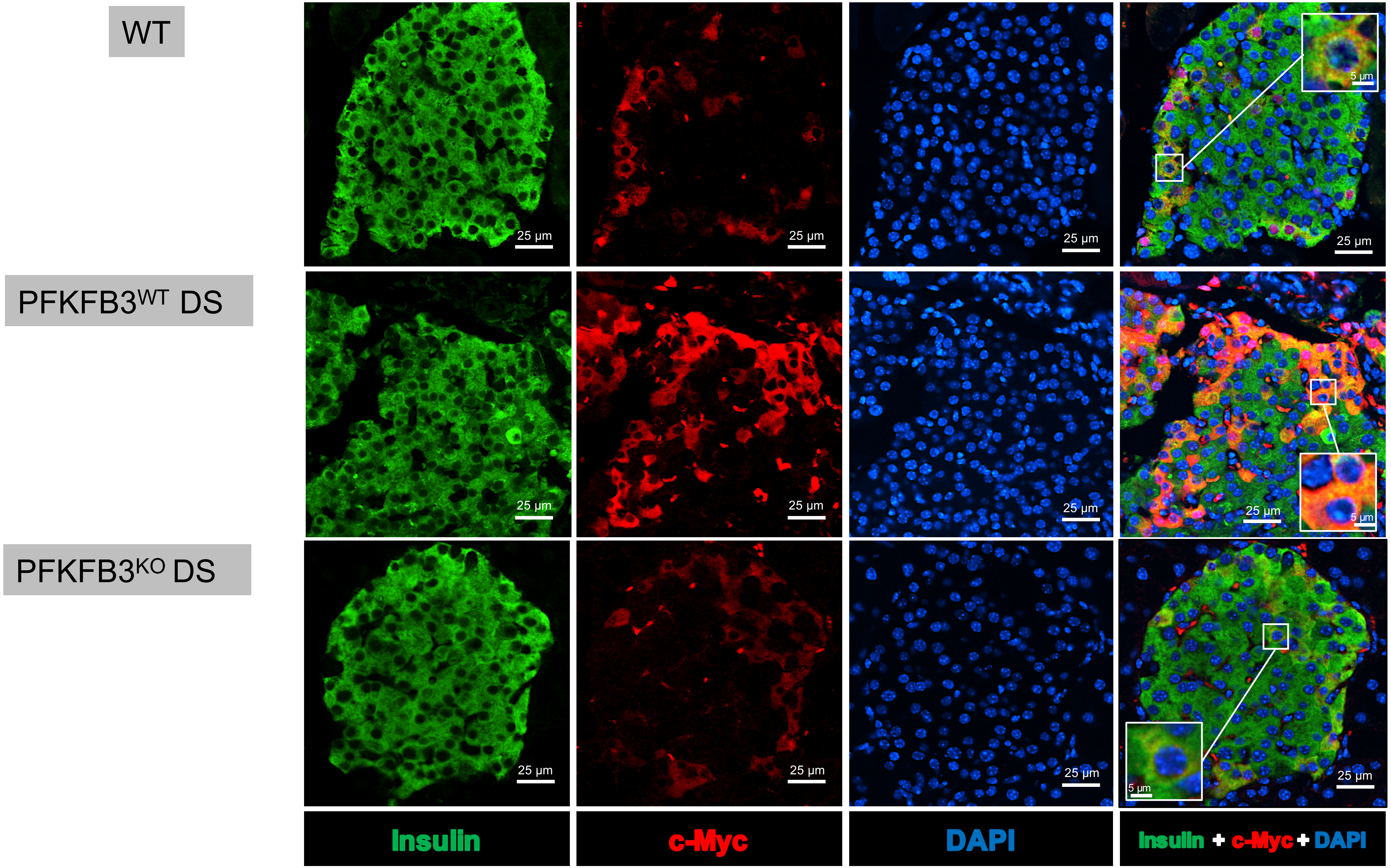

### Supplemental Data

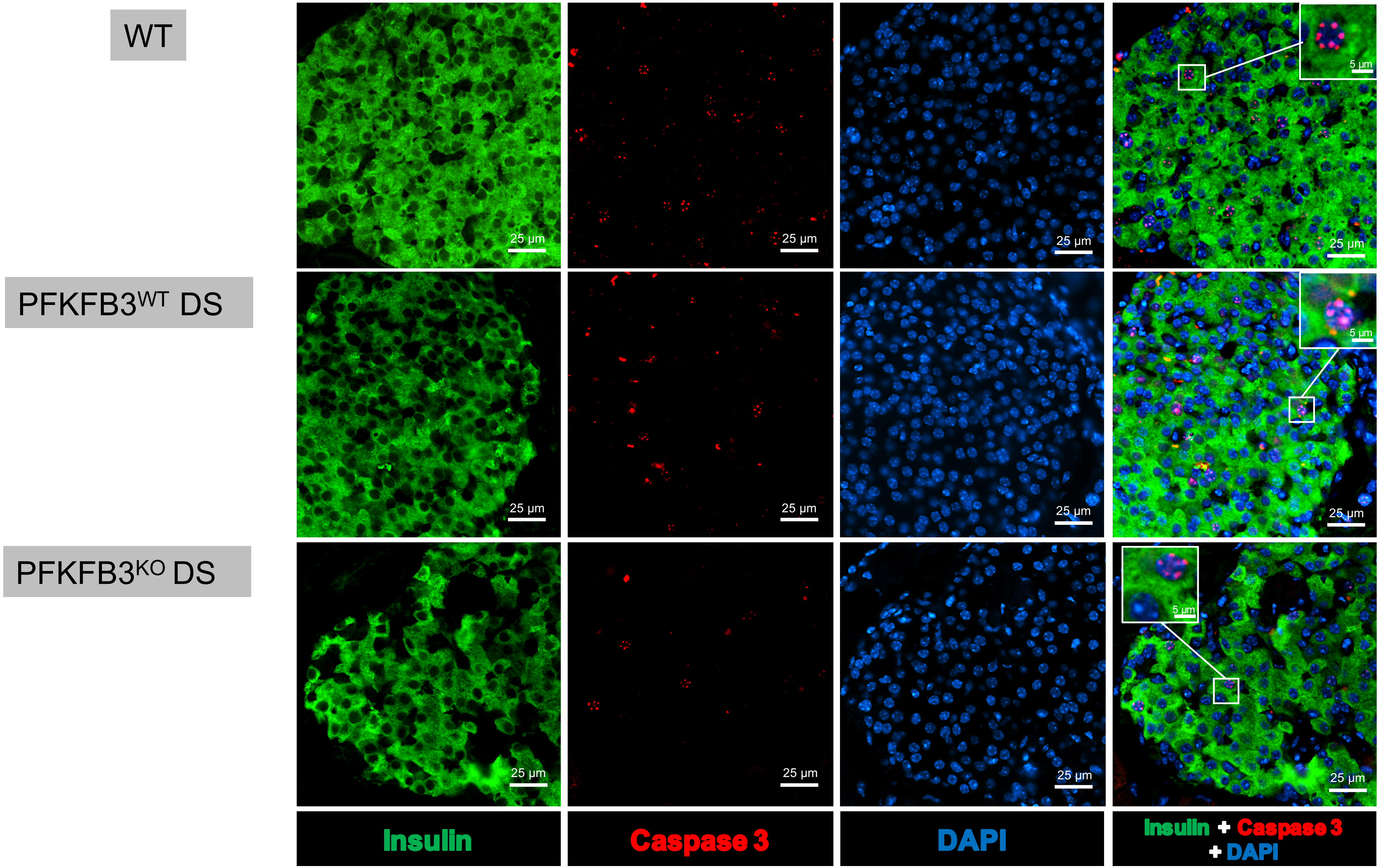

### Supplemental Data

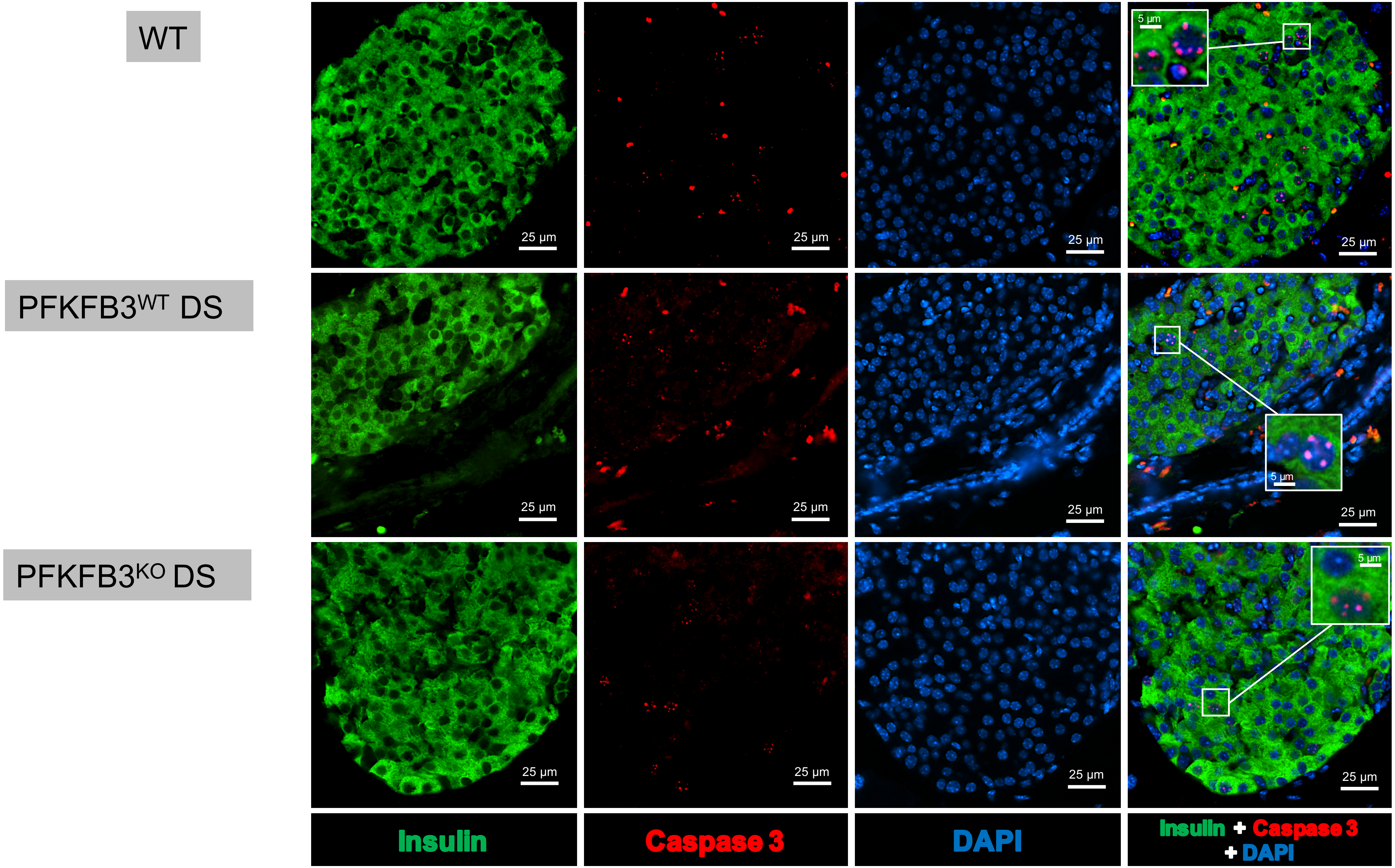

### Supplemental Data

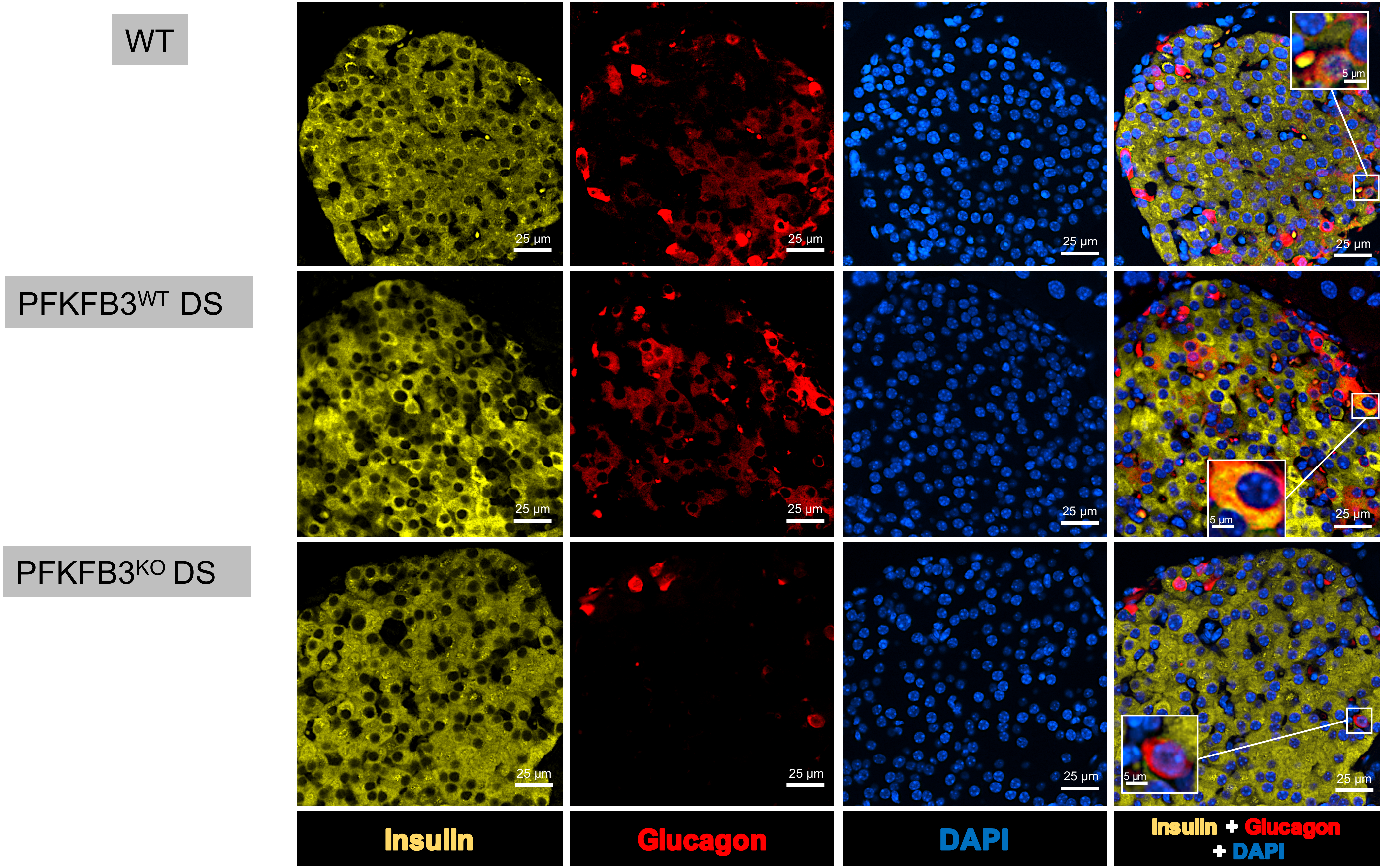

### Supplemental Data

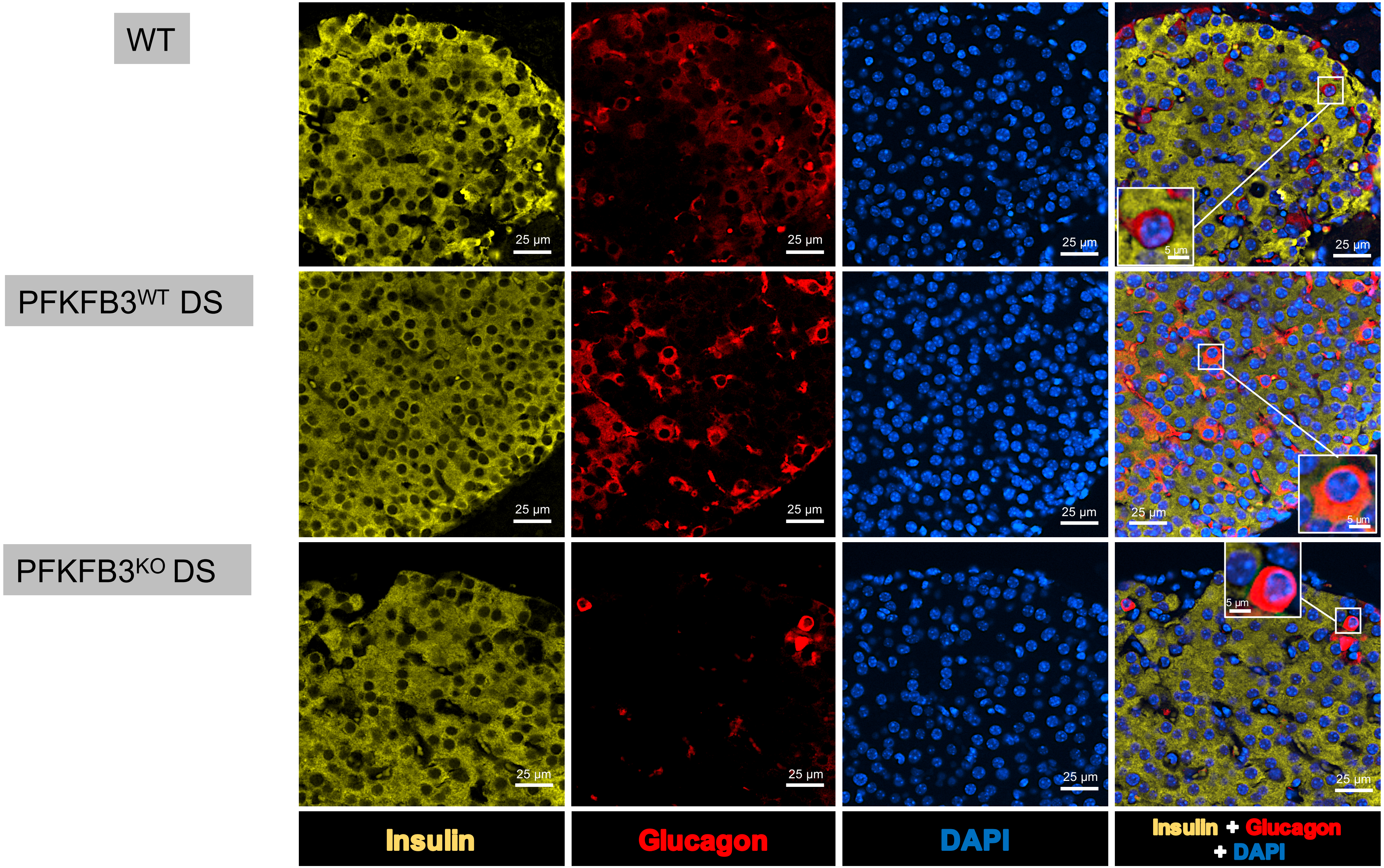

#### SUPPLEMENTARY FIGURE LEGENDS

***Supplementary Figure 1.*** ***Bodyweight of mice from experimental groups.*** *Bodyweight in indicated experimental groups (**A**) at the baseline (t=0) (**B**) 1 week before onset of HFD (**C**) at 4 weeks HFD, and (**D**) at 13 weeks HFD.*

***Supplementary Figure 2.*** ***Organ weight of mice from experimental groups.*** *Organ weight (g) in indicated experimental groups (**A**) of pancreas (**B**) of liver and (**C**) of spleen.*

***Supplementary Figure 3.*** ***Intraperitoneal-glucose tolerance test (IP-GTT) and Insulin tolerance test (ITT).*** *(**A**) IP-GTT after 7 weeks post HFD (**B**) ITT in mice after 8 weeks post HFD in mice 21-25 weeks of age.*

***Supplementary Figure 4.*** ***Quality of the cells from each donor.*** *Violin plots showing the distribution of number of genes (**A**), number of transcripts (**B**) and percentage of mitochondrial expression (**C**) in the cells from each donor.*

***Supplementary Figure 5.*** ***Quality of the cells from each cluster.*** *Violin plots showing the distribution of number of genes (**A**) number of transcripts (**B**) and percentage of mitochondrial expression (**C**) in the cells from each cluster.*

***Supplementary Figure 6.*** ***Distribution of annotated pancreatic cell types in non-diabetic versus T2D donors.*** *Relative contribution of nine annotated pancreatic cell types in non-diabetics (ND) and T2D is presented as a percentage (%).*

***Supplementary Figure 7.*** ***Heatmap showing the top marker genes for each cluster.*** *The marker genes were ranked by expression fold changes comparing the indicated cluster to all the other clusters. The color scale represents the scaled expression of the gene.*

***Supplementary Figure 8.*** ***Dotplot showing the top marker genes for each cluster.*** *The marker genes were ranked by expression fold changes comparing the indicated cluster to all the other clusters. The size of the dots represents percentage of cells the gene was detected in. The color scale represents the scaled expression of the gene.*

***Supplementary Figure 9.*** ***β-cell cluster differences in non-diabetics versus T2D donors***

*STRING analysis on data from [33] was performed to present the relationship between the differentially expressed genes in cluster 7 versus cluster 1 (**A**) in non-diabetics (**B**) in type-2 diabetes.*

***Supplementary Figure 10.*** ***Differences in LDHA-positive versus LDHA-negative β-cells in non-diabetic versus T2D donors.*** *We used STRING analysis to present the* *relationship between the differentially expressed genes [33] in LDHA-positive (cluster 7) versus LDHA-negative β-cells (cluster 1) (**A**) in non-diabetics (**B**) in type-2 diabetes.*

**Supplementary Figure 11.** ***β-cell differences in cluster 1 or LDHA-negative β-cells in non-diabetic versus T2D donors.*** *STRING analysis was performed to present the relationship between the differentially expressed genes [33] in non-diabetics versus T2D in either (**A**) cluster 1 or (**B**) LDHA-negative β-cells. No differences were observed in cluster 7 or LDHA-positive β-cells when non-diabetic- were compared to T2D donors.*

***SUPPLEMENTAL DATA***

***Supplemental Images -*** *single and merged images of the immunostainings performed in the study*

***Supplemental Tables 1-7***

Supplemental Table 1 cluster1_vs_cluster7 comparison

| | p_val | avg_logFC | pct.1 | pct.2 | p_val_adj |
|---|---|---|---|---|---|
| **GC** | 3.80180871976127E-59 | -1.89590400553957 | 0.092 | 1 | 1.77517854551813E-54 |
| **ARX** | 5.47729374696755E-58 | -0.295910419283131 | 0.038 | 0.879 | 2.55751276927156E-53 |
| **GPR119** | 1.05331111541449E-53 | -0.340029067362382 | 0.033 | 0.81 | 4.91822559120489E-49 |
| **LOXL4** | 3.43202865820207E-53 | -2.08270620179524 | 0.115 | 0.983 | 1.60251714137429E-48 |
| **TM4SF4** | 2.69586518243191E-52 | -2.31449898172568 | 0.148 | 1 | 1.25878032963293E-47 |
| **TMEM236** | 1.76873234778241E-51 | -0.499690080845802 | 0.047 | 0.81 | 8.25874195150042E-47 |
| **MRC1** | 2.55881053267494E-48 | -0.778248155533556 | 0.062 | 0.828 | 1.19478540202191E-43 |
| **SPOCK3** | 1.01204834146892E-46 | -0.458834106120558 | 0.036 | 0.724 | 4.72555732082083E-42 |
| **POPDC3** | 1.2459747517772E-45 | -0.274317815051722 | 0.009 | 0.603 | 5.81782990847328E-41 |
| **F10** | 3.4112117534631E-45 | -0.708281019405208 | 0.118 | 0.948 | 1.59279710404453E-40 |
| **CD36** | 3.39107219137448E-44 | -0.622903138482993 | 0.036 | 0.707 | 1.58339333831848E-39 |
| **IRX2** | 4.40652287763774E-44 | -0.374368294736417 | 0.089 | 0.828 | 2.05753772725539E-39 |
| **KCTD12** | 1.49165354502797E-43 | -0.685671173913432 | 0.148 | 0.948 | 6.9649778977991E-39 |
| **MUC13** | 2.85489573975476E-43 | -0.694002302449139 | 0.027 | 0.655 | 1.33303646776369E-38 |
| **FXYD3** | 1.27082092583167E-42 | -0.4458694318051 | 0.098 | 0.862 | 5.93384414898581E-38 |
| **PAPPA2** | 1.53891540824434E-42 | -0.506510505193441 | 0.038 | 0.69 | 7.18565771571528E-38 |
| **SERPINA1** | 3.5520245409303E-41 | -1.57670947945118 | 0.08 | 0.793 | 1.65854681889659E-36 |
| **MAOB** | 1.94951327320569E-40 | -0.373556543441814 | 0.033 | 0.655 | 9.10286232657932E-36 |
| **PTPRT** | 6.12761134679727E-40 | -0.281906123996622 | 0.041 | 0.672 | 2.86116556616005E-35 |
| **DPP4** | 6.2439253441593E-40 | -0.557254351476022 | 0.053 | 0.707 | 2.9154760609483E-35 |
| **DMD** | 7.59023314230943E-40 | -0.165473877374797 | 0.068 | 0.759 | 3.54410756113854E-35 |
| **TMEM176A** | 1.20240090529503E-39 | -1.0437562565174 | 0.269 | 1 | 5.61437054709409E-35 |
| **MYO10** | 3.52189346778314E-39 | -0.641434580231333 | 0.068 | 0.741 | 1.64447771691198E-34 |
| **S100A10** | 5.34132334050138E-39 | -0.566908402440228 | 0.124 | 0.879 | 2.49402410738031E-34 |
| **CRH** | 3.35393915824309E-37 | -0.638884243255757 | 0.03 | 0.603 | 1.56605481115845E-32 |
| **VIM** | 4.54254878173757E-37 | -1.17087672230074 | 0.207 | 0.931 | 2.12105230265672E-32 |
| **TMEM86B** | 6.23921949506403E-37 | -0.214979659021131 | 0.083 | 0.776 | 2.91327875883025E-32 |
| **FEV** | 6.874541814585E-37 | -0.552495165050059 | 0.151 | 0.879 | 3.20992980948417E-32 |
| **APOH** | 8.83874230103876E-36 | -0.502607811156132 | 0.044 | 0.638 | 4.12707394262403E-31 |
| **VIL1** | 7.20477000233462E-35 | -0.540202572674132 | 0.053 | 0.638 | 3.36412325719011E-30 |
| **GPC6** | 2.06376461282658E-34 | -0.315680340994895 | 0.059 | 0.672 | 9.63633610667117E-30 |
| **TMEM176B** | 2.32582229685332E-34 | -1.25576578093544 | 0.364 | 0.983 | 1.08599620506972E-29 |
| **FXYD5** | 4.17609368525379E-34 | -0.569084472213089 | 0.139 | 0.845 | 1.94994342445555E-29 |

| | | | | | |
|---|---|---|---|---|---|
| **TTR** | 2.56701775384417E-33 | -2.40765432379202 | 0.962 | 1 | 1.19861759980246E-28 |
| **SMIM24** | 3.89353754306973E-33 | -0.787737535264275 | 0.172 | 0.879 | 1.81800948498555E-28 |
| **FAP** | 1.55217547369334E-32 | -0.591875979732712 | 0.118 | 0.741 | 7.24757293931631E-28 |
| **FABP5** | 1.59787484923056E-32 | -0.37065850088586 | 0.163 | 0.845 | 7.46095703351225E-28 |
| **GPER1** | 2.36485781356378E-32 | -0.193905024802642 | 0.024 | 0.517 | 1.10422305888733E-27 |
| **GCG** | 3.47752228366952E-32 | -3.84974706949689 | 0.763 | 1 | 1.62375947991381E-27 |
| **ADGRG2** | 3.89368856571169E-31 | -0.526924677528466 | 0.112 | 0.741 | 1.81808000198776E-26 |
| **PPP2R2B** | 4.32413421244322E-31 | -0.243053685292869 | 0.05 | 0.603 | 2.01906798781611E-26 |
| **LDHA** | 1.54798063684808E-30 | -0.80316614074208 | 0.142 | 0.81 | 7.22798598763476E-26 |
| **IGFBP2** | 1.78998426596299E-30 | -0.957707942347793 | 0.299 | 0.966 | 8.35797353306099E-26 |
| **NPNT** | 1.16192494706539E-29 | -0.404835313985191 | 0.062 | 0.621 | 5.42537615533242E-25 |
| **ADGRG5** | 1.43045825021326E-28 | -0.171906928755954 | 0.038 | 0.534 | 6.67923870772079E-24 |
| **USH1C** | 1.84587028802463E-28 | -0.47387126369174 | 0.16 | 0.828 | 8.61892213587338E-24 |
| **LAMB1** | 2.48827854329068E-28 | -0.422807411584514 | 0.189 | 0.81 | 1.16185190021872E-23 |
| **RNF144A** | 3.66865330223363E-28 | -0.134511466265907 | 0.033 | 0.5 | 1.71300428641195E-23 |
| **SEMA3E** | 5.85025934137705E-28 | -0.186144716934954 | 0.036 | 0.5 | 2.73166159426918E-23 |
| **KIF12** | 9.38402772270483E-28 | -0.180625978392371 | 0.13 | 0.759 | 4.38168406456257E-23 |
| **SPTSSB** | 1.47265876773256E-27 | -0.609320911097299 | 0.145 | 0.793 | 6.87628558417363E-23 |
| **MBOAT4** | 2.14648436365604E-27 | -0.347093275894926 | 0.059 | 0.569 | 1.00225794392191E-22 |
| **SLC6A19** | 2.94710776803652E-27 | -0.287243382042475 | 0.03 | 0.483 | 1.37609303012929E-22 |
| **KCNIP1** | 9.69982158286035E-27 | -0.225276583392828 | 0.027 | 0.466 | 4.52913769168498E-22 |
| **ANK2** | 1.14691603782256E-26 | -0.242976425613359 | 0.101 | 0.672 | 5.35529505540488E-22 |
| **RGS4** | 1.84842697114261E-26 | -1.15097198736651 | 0.346 | 0.948 | 8.63086005635617E-22 |
| **CRYBA2** | 7.69728078268479E-25 | -1.61854685808147 | 0.722 | 1 | 3.59409131585901E-20 |
| **REG4** | 1.10540362731695E-24 | -0.841807088592046 | 0.041 | 0.483 | 5.16146115703106E-20 |
| **AOX1** | 1.16498098746446E-24 | -0.259163582285894 | 0.036 | 0.466 | 5.43964572476782E-20 |
| **PYROXD2** | 2.69656063465583E-24 | -0.109617463655448 | 0.038 | 0.483 | 1.25910505713985E-19 |
| **CLU** | 2.98323106106261E-24 | -1.31016422298528 | 0.929 | 1 | 1.39296007934196E-19 |
| **PLPPR1** | 4.07748851539375E-24 | -0.228020971772917 | 0.024 | 0.414 | 1.90390171249281E-19 |
| **SYNDIG1** | 6.93368418059876E-24 | -0.298381418579345 | 0.041 | 0.483 | 3.23754515444698E-19 |
| **CSGALNACT** | 9.20973137162732E-24 | -0.223981878614721 | 0.059 | 0.534 | 4.30029986935394E-19 |
| **PRSS3** | 1.5526547953033E-23 | -0.242113401521143 | 0.053 | 0.5 | 7.24981103570971E-19 |
| **EGFL7** | 2.23949051138785E-23 | -0.253532542030301 | 0.172 | 0.793 | 1.04568530448233E-18 |
| **SLC38A4** | 7.75492707318194E-23 | -0.364168626629373 | 0.228 | 0.862 | 3.62100809828084E-18 |
| **FAM110B** | 1.57480963490794E-22 | -0.253391047696972 | 0.151 | 0.707 | 7.35325862827565E-18 |
| **LDB2** | 1.94665116052713E-22 | -0.181258226741596 | 0.021 | 0.379 | 9.08949826384932E-18 |

| | | | | | |
|---|---|---|---|---|---|
| **MYRF** | 2.41844879105585E-22 | -0.182944702673991 | 0.065 | 0.517 | 1.12924629400771E-17 |
| **PLIN3** | 5.60548666758567E-22 | -0.19781873211174 | 0.109 | 0.655 | 2.61736988969578E-17 |
| **CD82** | 6.40517467378991E-22 | -0.4613707917827 | 0.26 | 0.862 | 2.99076821043272E-17 |
| **HNF1A-AS1** | 1.31089413929297E-21 | -0.140333740418626 | 0.098 | 0.603 | 6.12095800460066E-17 |
| **GLS** | 1.59969301063988E-21 | -0.664855732772313 | 0.536 | 1 | 7.46944657458081E-17 |
| **SLC7A14** | 2.70255155313775E-21 | -0.378089891030948 | 0.207 | 0.793 | 1.26190239670661E-16 |
| **KCNJ3** | 3.37955791889527E-21 | -0.130163556649339 | 0.107 | 0.603 | 1.57801697906977E-16 |
| **CYSTM1** | 3.43601599146509E-21 | -0.679907246645856 | 0.932 | 1 | 1.60437894689479E-16 |
| **SPATS2L** | 5.45667731830059E-21 | -0.303481035451314 | 0.216 | 0.793 | 2.54788634023409E-16 |
| **C1orf21** | 5.66394778405606E-21 | -0.103496644024633 | 0.027 | 0.379 | 2.6446671388093E-16 |
| **CNIH3** | 5.79275579805157E-21 | -0.236169581842531 | 0.104 | 0.603 | 2.70481146478422E-16 |
| **KCNH6** | 6.77373476215036E-21 | -0.258912302318725 | 0.16 | 0.724 | 3.16285997249087E-16 |
| **ERBB3** | 7.45948626289895E-21 | -0.252272333767288 | 0.065 | 0.5 | 3.4830579207354E-16 |
| **TMEM45B** | 7.51342225259228E-21 | -0.194617463791683 | 0.03 | 0.397 | 3.50824225240291E-16 |
| **SERPINA10** | 8.57273718583465E-21 | -0.157347330919362 | 0.021 | 0.362 | 4.00286817418177E-16 |
| **POU6F2** | 1.29597817848255E-20 | -0.112566042287548 | 0.056 | 0.466 | 6.05131090878859E-16 |
| **PRLR** | 2.12016652589937E-20 | -0.166710026366053 | 0.033 | 0.397 | 9.89969355938195E-16 |
| **CEACAM1** | 2.3268779921371E-20 | -0.285219924942378 | 0.104 | 0.603 | 1.08648914086858E-15 |
| **SLC44A3** | 2.50074802518989E-20 | -0.120751453434284 | 0.038 | 0.431 | 1.16767427540192E-15 |
| **HS3ST3B1** | 3.51380367545717E-20 | -0.171662091858596 | 0.041 | 0.431 | 1.64070035018122E-15 |
| **STK32B** | 6.05933560762234E-20 | -0.123160485854962 | 0.003 | 0.259 | 2.8292855752671E-15 |
| **RHOC** | 8.58346888150376E-20 | -0.246762090919656 | 0.095 | 0.552 | 4.00787912484055E-15 |
| **ARFGEF3** | 1.06148772278188E-19 | -0.754178910914446 | 0.855 | 1 | 4.95640462398545E-15 |
| **KLHL41** | 1.08350897740335E-19 | -0.274642179999936 | 0.077 | 0.517 | 5.05922846818946E-15 |
| **INS** | 1.11015214223568E-19 | 0.72969422468799 | 1 | 1 | 5.18363339774105E-15 |
| **IMPA2** | 1.75886965069793E-19 | -0.165164968648153 | 0.056 | 0.466 | 8.21269006000385E-15 |
| **QDPR** | 3.02281255619857E-19 | -0.754469263875519 | 0.787 | 0.983 | 1.4114418668658E-14 |
| **COTL1** | 3.16080971614721E-19 | -0.540100147143886 | 0.364 | 0.931 | 1.47587688076062E-14 |
| **ABHD15** | 5.97714608130366E-19 | -0.242770308660086 | 0.163 | 0.707 | 2.79090881974312E-14 |
| **CPNE8** | 1.37542273981448E-18 | -0.315963761671105 | 0.263 | 0.879 | 6.42226139901575E-14 |
| **HMGB3** | 1.39232995583625E-18 | -0.411120903721079 | 0.293 | 0.845 | 6.50120626278622E-14 |
| **ANXA4** | 2.6846657745393E-18 | -0.313721177414564 | 0.228 | 0.828 | 1.25355099010564E-13 |
| **CDO1** | 6.1202575311721E-18 | -0.199231471220749 | 0.015 | 0.293 | 2.85773184903019E-13 |
| **WIPF3** | 7.68135859844012E-18 | -0.188017485663392 | 0.044 | 0.397 | 3.58665677036964E-13 |
| **TSPAN14** | 1.06786801573446E-17 | -0.217540295079628 | 0.157 | 0.638 | 4.98619612586892E-13 |
| **PLCE1** | 1.16305562873299E-17 | -0.697783746112533 | 0.183 | 0.69 | 5.43065564724296E-13 |

| | | | | | |
|---|---|---|---|---|---|
| **KLB** | 1.41834818038879E-17 | -0.147137709508135 | 0.033 | 0.362 | 6.62269315868938E-13 |
| **ITPR1** | 1.69943737164397E-17 | -0.156600887023552 | 0.133 | 0.621 | 7.93518291941717E-13 |
| **CAMK2G** | 2.20038542229552E-17 | -0.47486814974375 | 0.346 | 0.897 | 1.02742596523245E-12 |
| **COL27A1** | 2.47807935204606E-17 | -0.154377069751298 | 0.139 | 0.621 | 1.15708959185087E-12 |
| **STK32A** | 2.70500083046694E-17 | -0.118890168110059 | 0.021 | 0.31 | 1.26304603776993E-12 |
| **ARRDC4** | 3.6347656496268E-17 | -0.348633527458798 | 0.145 | 0.621 | 1.69718112478024E-12 |
| **FGFR4** | 4.91264865599677E-17 | -0.120068212459394 | 0.03 | 0.345 | 2.29386303694457E-12 |
| **TPD52L1** | 5.4064551214896E-17 | -0.226060831285126 | 0.065 | 0.448 | 2.52443608987714E-12 |
| **DSC2** | 7.50265312937187E-17 | -0.130507772660043 | 0.059 | 0.431 | 3.50321382569761E-12 |
| **GATA6** | 8.28946268357068E-17 | -0.100776470918767 | 0.033 | 0.345 | 3.87059881083966E-12 |
| **ITGB8** | 9.37698312196246E-17 | -0.140502167599553 | 0.121 | 0.586 | 4.37839472913793E-12 |
| **NEDD9** | 9.87083677299949E-17 | -0.268832050911474 | 0.139 | 0.603 | 4.60898981441665E-12 |
| **PDK4** | 1.02720034107338E-16 | -1.06623489136769 | 0.506 | 0.966 | 4.79630655257395E-12 |
| **GRIA3** | 1.08168451255644E-16 | -0.452917014958378 | 0.414 | 0.931 | 5.05070949447978E-12 |
| **DOK4** | 1.37654193943449E-16 | -0.199096912032705 | 0.142 | 0.638 | 6.42748727780145E-12 |
| **LBH** | 2.05980925171943E-16 | -0.149408265481876 | 0.056 | 0.414 | 9.61786733905351E-12 |
| **A1CF** | 2.24396587616068E-16 | -0.297237083070788 | 0.263 | 0.793 | 1.04777498655571E-11 |
| **PRKCE** | 3.54780068431153E-16 | -0.112635723929541 | 0.186 | 0.69 | 1.65657457352558E-11 |
| **SLC5A1** | 4.61521834093063E-16 | -0.237714445774137 | 0.065 | 0.431 | 2.15498389993074E-11 |
| **EZR** | 4.69717698898403E-16 | -0.614147879970267 | 0.757 | 1 | 2.19325285146631E-11 |
| **FAM131C** | 4.84146348150199E-16 | -0.112742576307627 | 0.216 | 0.741 | 2.26062454341772E-11 |
| **NPHP4** | 4.99664166147935E-16 | -0.152724072639645 | 0.101 | 0.517 | 2.33308189099455E-11 |
| **TSPAN15** | 5.56495904036555E-16 | -0.128594248362879 | 0.009 | 0.241 | 2.59844632471789E-11 |
| **CYP4F3** | 6.05677190328653E-16 | -0.142368535637928 | 0.024 | 0.31 | 2.82808850480158E-11 |
| **SH3RF1** | 6.25927399945803E-16 | -0.189610645243994 | 0.112 | 0.534 | 2.92264280856694E-11 |
| **UPB1** | 1.04789186535213E-15 | -0.103560546093573 | 0.027 | 0.31 | 4.89292148688868E-11 |
| **SLC40A1** | 2.40026802683192E-15 | -0.53608883414356 | 0.263 | 0.776 | 1.12075714976863E-10 |
| **MATN2** | 2.85610539020761E-15 | -0.340962629214117 | 0.219 | 0.741 | 1.33360128984964E-10 |
| **PALLD** | 4.01946953282147E-15 | -0.581212459052514 | 0.506 | 0.948 | 1.87681090896033E-10 |
| **RIIAD1** | 4.65199054883375E-15 | -0.10307098915898 | 0.175 | 0.655 | 2.17215394696694E-10 |
| **VGF** | 4.92283248003882E-15 | -0.281731638394842 | 0.731 | 1 | 2.29861816990453E-10 |
| **TMEM56** | 5.28600159660056E-15 | -0.119713901589685 | 0.074 | 0.431 | 2.4681927255007E-10 |
| **PTGER4** | 7.38323665239003E-15 | -0.204925222321456 | 0.041 | 0.345 | 3.44745469010048E-10 |
| **HGD** | 7.78000572297845E-15 | -0.12479715483546 | 0.059 | 0.414 | 3.63271807223033E-10 |
| **SMOC1** | 8.35487559239854E-15 | -0.183593219526026 | 0.13 | 0.569 | 3.90114206035865E-10 |
| **INHBA** | 1.26635293605278E-14 | -0.339943511334142 | 0.207 | 0.69 | 5.91298176431124E-10 |

| | | | | | |
|---|---|---|---|---|---|
| **CBR3-AS1** | 1.533397831424E-14 | -0.155892824016742 | 0.195 | 0.672 | 7.15989449426808E-10 |
| **ELL2** | 1.55960245950902E-14 | -0.46541416636037 | 0.76 | 1 | 7.28225176418546E-10 |
| **ZFP36** | 1.98402463325092E-14 | -0.232015228901594 | 0.213 | 0.707 | 9.26400622003853E-10 |
| **TJP2** | 2.19675013727742E-14 | -0.141199795630552 | 0.077 | 0.448 | 1.02572854159895E-09 |
| **FSTL5** | 2.41381827609991E-14 | -0.124177278703497 | 0.107 | 0.517 | 1.12708416765933E-09 |
| **PLXDC2** | 2.68684318163889E-14 | -0.14071317979755 | 0.059 | 0.379 | 1.25456768680265E-09 |
| **ANG** | 3.14641025972103E-14 | -0.104287309681457 | 0.027 | 0.293 | 1.46915334257154E-09 |
| **SSX2IP** | 3.51432523643216E-14 | -0.324071504094603 | 0.343 | 0.897 | 1.64094388264727E-09 |
| **MFSD3** | 3.52075221111501E-14 | -0.218565950651333 | 0.331 | 0.845 | 1.64394482993593E-09 |
| **PDE3B** | 4.25016571404087E-14 | -0.240922239240638 | 0.385 | 0.914 | 1.9845298768571E-09 |
| **SLC7A8** | 4.28114587368489E-14 | -0.642338877258115 | 0.935 | 1 | 1.99899544279968E-09 |
| **BTBD11** | 4.37980629829332E-14 | -0.114064369508386 | 0.056 | 0.379 | 2.0450629548621E-09 |
| **SERTM1** | 4.42661626269969E-14 | -0.111483630630322 | 0.018 | 0.259 | 2.06691993154236E-09 |
| **PCSK2** | 9.57104724916984E-14 | -0.647839421620055 | 0.932 | 1 | 4.46900909205488E-09 |
| **ELF3** | 1.15191570797899E-13 | -0.347221587976907 | 0.296 | 0.776 | 5.37864001526632E-09 |
| **SLC7A2** | 1.60870578841783E-13 | -0.512839441407065 | 0.657 | 1 | 7.5115299378594E-09 |
| **AGFG2** | 1.82583828481783E-13 | -0.137177522764718 | 0.169 | 0.621 | 8.52538670329988E-09 |
| **MYL6** | 1.9047404024076E-13 | -0.384158890047797 | 1 | 1 | 8.89380436096181E-09 |
| **RPS6KA3** | 2.00472337135664E-13 | -0.343824052331616 | 0.464 | 0.914 | 9.36065483787556E-09 |
| **VSNL1** | 2.17836682632058E-13 | -0.122028572127038 | 0.056 | 0.362 | 1.01714482221387E-08 |
| **LMCD1** | 2.66190187618752E-13 | -0.250196750976684 | 0.124 | 0.534 | 1.24292184304824E-08 |
| **ALDH1A1** | 2.73134275610881E-13 | -0.732603188473602 | 0.962 | 1 | 1.27534587310989E-08 |
| **IFIH1** | 2.91512779094316E-13 | -0.131374013940563 | 0.16 | 0.621 | 1.36116061942509E-08 |
| **SNTG1** | 2.96346737450126E-13 | -0.132867636220502 | 0.186 | 0.638 | 1.38373182117587E-08 |
| **BID** | 3.66412058644442E-13 | -0.245982470501195 | 0.204 | 0.621 | 1.71088782542849E-08 |
| **TMSB4X** | 4.06126195109697E-13 | -0.660358187894644 | 0.87 | 1 | 1.89632504282571E-08 |
| **CHGA** | 6.52990946222069E-13 | -0.675955512023227 | 0.985 | 1 | 3.04901062519471E-08 |
| **GALNT13** | 8.4562172145241E-13 | -0.206143351848593 | 0.098 | 0.448 | 3.94846150397774E-08 |
| **TMPRSS2** | 1.23738837469753E-12 | -0.110501494535216 | 0.015 | 0.224 | 5.77773753797519E-08 |
| **GNG5** | 1.27930324429886E-12 | -0.334392014235897 | 0.63 | 1 | 5.97345063860467E-08 |
| **KIAA1217** | 1.5033233412325E-12 | -0.117419938173064 | 0.18 | 0.621 | 7.01946767721691E-08 |
| **SMIM5** | 1.98126455229831E-12 | -0.150017000687261 | 0.095 | 0.448 | 9.25111857404652E-08 |
| **HPS1** | 2.26925502002612E-12 | -0.515803656907961 | 0.58 | 0.914 | 1.05958324650079E-07 |
| **IAPP** | 2.27683387319997E-12 | 1.4361529534897 | 0.976 | 0.966 | 1.06312204041326E-07 |
| **MPC2** | 2.37214517483037E-12 | -0.455059714855242 | 0.814 | 0.983 | 1.10762574648354E-07 |
| **TCN2** | 2.50992968916924E-12 | -0.103320761981907 | 0.109 | 0.483 | 1.17196146976379E-07 |

| | | | | | |
|---|---|---|---|---|---|
| **SLC6A4** | 2.70173496242615E-12 | -0.25637299935001 | 0.201 | 0.638 | 1.26152110600564E-07 |
| **FAM84A** | 2.74113414150257E-12 | -0.231621055635555 | 0.213 | 0.655 | 1.2799177646918E-07 |
| **KCNA5** | 3.47134863637079E-12 | -0.170598181308537 | 0.189 | 0.655 | 1.62087681878061E-07 |
| **SNED1** | 3.59777197533481E-12 | -0.107543575313822 | 0.178 | 0.586 | 1.67990766844309E-07 |
| **TMEM178B** | 4.12063632235838E-12 | -0.148916924150524 | 0.284 | 0.741 | 1.9240487179988E-07 |
| **IRF6** | 4.51218424186857E-12 | -0.175295156919494 | 0.225 | 0.655 | 2.10687418805569E-07 |
| **MACC1** | 5.17741820525375E-12 | -0.10233770336672 | 0.118 | 0.466 | 2.41749188257913E-07 |
| **RBM47** | 6.2676563572955E-12 | -0.278427111859994 | 0.5 | 0.862 | 2.92655678291199E-07 |
| **HIGD1A** | 6.35492504791932E-12 | -0.511938723231059 | 0.725 | 0.966 | 2.96730515262497E-07 |
| **PDZK1** | 7.0966120627768E-12 | -0.152014285766821 | 0.089 | 0.414 | 3.31362107047237E-07 |
| **SELENBP1** | 8.30277016798396E-12 | -0.115143716926136 | 0.033 | 0.276 | 3.87681247453675E-07 |
| **ITPK1** | 8.77440490886894E-12 | -0.162534545833591 | 0.37 | 0.845 | 4.09703288409817E-07 |
| **AC006369.2** | 9.81631020825841E-12 | -0.235227046917101 | 0.45 | 0.879 | 4.5835297255421E-07 |
| **LPP** | 1.0707558034182E-11 | -0.276590907767687 | 0.459 | 0.897 | 4.9996800729006E-07 |
| **HACD1** | 1.07522903624532E-11 | -0.168803632985545 | 0.275 | 0.793 | 5.02056693894029E-07 |
| **STXBP6** | 1.12836453574524E-11 | -0.121211619167425 | 0.104 | 0.448 | 5.26867252675523E-07 |
| **LUZP1** | 1.78947225157635E-11 | -0.162713403003445 | 0.121 | 0.483 | 8.35558278428546E-07 |
| **LINC00473** | 1.82021386279543E-11 | -0.343969199602023 | 0.272 | 0.707 | 8.4991245895507E-07 |
| **FAM83F** | 1.96534119498213E-11 | -0.114266280172955 | 0.21 | 0.638 | 9.17676764173007E-07 |
| **CNN3** | 2.66136659034625E-11 | -0.146941332827129 | 0.145 | 0.534 | 1.24267190203037E-06 |
| **EGFEM1P** | 2.86642619971196E-11 | -0.309524114195873 | 0.222 | 0.621 | 1.3384203854315E-06 |
| **EFNA1** | 2.93224327789598E-11 | -0.142588726847427 | 0.059 | 0.345 | 1.36915235374797E-06 |
| **GPX3** | 2.94828900743003E-11 | -0.576617566477536 | 0.817 | 1 | 1.3766445862393E-06 |
| **SAMD5** | 3.09571718932056E-11 | -0.127077470585908 | 0.213 | 0.638 | 1.44548322720945E-06 |
| **TMEM8A** | 3.4092265547954E-11 | -0.209359353448672 | 0.382 | 0.845 | 1.59187015523062E-06 |
| **VSTM2A** | 3.4394468451429E-11 | -0.201513383168101 | 0.021 | 0.224 | 1.60598091540257E-06 |
| **CLSTN2** | 3.47494786529077E-11 | -0.150290280633804 | 0.098 | 0.431 | 1.62255740674022E-06 |
| **HADH** | 4.45826486072762E-11 | 0.696725872800164 | 0.976 | 0.966 | 2.08169761141955E-06 |
| **SAMD4A** | 5.91144280151473E-11 | -0.107500715306047 | 0.107 | 0.431 | 2.76022998731127E-06 |
| **ATP2B4** | 6.32423187605132E-11 | -0.144331189172768 | 0.175 | 0.569 | 2.95297358988464E-06 |
| **REEP4** | 6.56034688436536E-11 | -0.184452560332196 | 0.325 | 0.845 | 3.06322277071672E-06 |
| **GDF15** | 7.61113612730117E-11 | -0.220426414996274 | 0.157 | 0.552 | 3.55386779192073E-06 |
| **ADCY6** | 8.40075173953128E-11 | -0.121350357395423 | 0.083 | 0.397 | 3.92256300973934E-06 |
| **KCNJ6** | 8.60637512401028E-11 | -0.374846583535666 | 0.678 | 0.983 | 4.01857473665412E-06 |
| **NRP1** | 8.66627860674077E-11 | -0.218758851899479 | 0.18 | 0.569 | 4.04654546984547E-06 |
| **PFKP** | 8.74174569673549E-11 | -0.291410890886657 | 0.417 | 0.862 | 4.0817833181767E-06 |

| | | | | | |
|---|---|---|---|---|---|
| **MARCKSL1** | 9.54968376070063E-11 | -0.142141750867642 | 0.148 | 0.5 | 4.45903383838394E-06 |
| **SERINC5** | 1.10454040242552E-10 | -0.167423224524634 | 0.249 | 0.638 | 5.15743050104548E-06 |
| **FGF12** | 1.10536887040564E-10 | -0.261144759738786 | 0.388 | 0.897 | 5.16129886658507E-06 |
| **B2M** | 1.12692285672589E-10 | -0.451970056615722 | 1 | 1 | 5.26194089491021E-06 |
| **AC009014.3** | 1.19000935386796E-10 | -0.11315977777839 | 0.112 | 0.466 | 5.55651067601565E-06 |
| **CACNB2** | 1.44287647730348E-10 | -0.372125370922079 | 0.538 | 0.862 | 6.73722313547314E-06 |
| **CNTN1** | 1.60419891678244E-10 | -0.339638727168569 | 0.553 | 0.966 | 7.49048600213227E-06 |
| **CCDC86** | 2.21243158450035E-10 | -0.24934161860261 | 0.385 | 0.862 | 1.03305067975075E-05 |
| **TAGLN2** | 2.3783787145709E-10 | -0.538462161692373 | 0.876 | 1 | 1.11053637319459E-05 |
| **GLRX** | 2.52548945344678E-10 | -0.233020132201726 | 0.281 | 0.69 | 1.1792267904979E-05 |
| **LRRFIP1** | 2.52976325283895E-10 | -0.387743012572566 | 0.586 | 0.948 | 1.18122235564809E-05 |
| **WWC1** | 2.55320287400863E-10 | -0.165841948511034 | 0.266 | 0.69 | 1.19216701796085E-05 |
| **STARD3** | 3.30607408405589E-10 | -0.215991244081393 | 0.355 | 0.776 | 1.54370517206822E-05 |
| **SERPINE2** | 3.39094844166738E-10 | -0.329942633497817 | 0.399 | 0.879 | 1.58333555586775E-05 |
| **CCDC150** | 3.72387442453804E-10 | -0.109031047400979 | 0.083 | 0.379 | 1.73878868504955E-05 |
| **SYDE2** | 3.79631570175365E-10 | -0.152614129664882 | 0.071 | 0.345 | 1.77261369061983E-05 |
| **ASS1** | 3.88981777426243E-10 | -0.215336859445896 | 0.269 | 0.707 | 1.81627261333636E-05 |
| **USP25** | 4.13318061668086E-10 | -0.107849207347359 | 0.222 | 0.638 | 1.92990602534679E-05 |
| **HLA-A** | 4.33304735756765E-10 | -0.482380246335455 | 0.973 | 1 | 2.02322980266906E-05 |
| **FOSL2** | 4.59101704744447E-10 | -0.174190495075589 | 0.272 | 0.69 | 2.14368358996325E-05 |
| **LIMA1** | 4.84450290287611E-10 | -0.174471476634676 | 0.349 | 0.81 | 2.26204374043994E-05 |
| **SAMD11** | 4.87616807496453E-10 | 0.394236117503495 | 0.964 | 0.948 | 2.27682915924319E-05 |
| **ARHGEF28** | 5.04555076608175E-10 | -0.136995689847945 | 0.275 | 0.69 | 2.35591901920655E-05 |
| **GS1-358P8.4** | 5.12514570064475E-10 | -0.147167409286517 | 0.186 | 0.552 | 2.39308428200205E-05 |
| **SQRDL** | 5.20400918388983E-10 | -0.134019326745571 | 0.178 | 0.569 | 2.42990800823368E-05 |
| **CNKSR3** | 5.24858378627382E-10 | -0.202999267435809 | 0.32 | 0.776 | 2.45072122732483E-05 |
| **RAP2B** | 6.19002107884137E-10 | -0.24620214318526 | 0.349 | 0.793 | 2.8903065423434E-05 |
| **LAT2** | 6.33692925650045E-10 | -0.109540392585601 | 0.05 | 0.293 | 2.95890237773775E-05 |
| **TM4SF1** | 6.59284770915622E-10 | -0.244293051057995 | 0.068 | 0.328 | 3.07839838083631E-05 |
| **CD99L2** | 6.84618927442363E-10 | -0.247609515866979 | 0.47 | 0.897 | 3.19669115790662E-05 |
| **TTLL12** | 7.08230865709162E-10 | -0.115938560911736 | 0.207 | 0.603 | 3.30694238125579E-05 |
| **MAGIX** | 7.90932914853701E-10 | -0.102000081173696 | 0.228 | 0.655 | 3.69310305932638E-05 |
| **PCSK1** | 7.98064778683927E-10 | 0.825023076159184 | 0.973 | 0.983 | 3.72640387110886E-05 |
| **COX8A** | 8.20280502345633E-10 | -0.267535230208855 | 0.973 | 1 | 3.83013574960246E-05 |
| **HMGCS2** | 8.23340211312731E-10 | -0.176462904874732 | 0.033 | 0.241 | 3.84442244868254E-05 |
| **HSPB1** | 8.86254110628251E-10 | -0.270134173831373 | 0.586 | 0.966 | 4.13818631875649E-05 |

| | | | | | |
|---|---|---|---|---|---|
| **TRNP1** | 9.1793431360427E-10 | -0.143247484973597 | 0.34 | 0.724 | 4.28611069051242E-05 |
| **CNNM1** | 9.85393033734205E-10 | -0.189584002222509 | 0.382 | 0.828 | 4.60109569241512E-05 |
| **TPM4** | 1.07949760808121E-09 | -0.413212269533318 | 0.615 | 0.966 | 5.0404981814136E-05 |
| **NBEAL2** | 1.15333211789837E-09 | -0.183640134545618 | 0.254 | 0.621 | 5.38525365810288E-05 |
| **PCDH17** | 1.1657413657821E-09 | -0.270175352388178 | 0.598 | 0.914 | 5.44319615924636E-05 |
| **MLIP** | 1.21326174643726E-09 | -0.123430011113372 | 0.109 | 0.431 | 5.66508307263951E-05 |
| **CYP3A5** | 1.5609049574593E-09 | -0.167058212763321 | 0.077 | 0.345 | 7.28833351786471E-05 |
| **CD63** | 1.57382850560337E-09 | -0.325340730957166 | 1 | 1 | 7.34867744121381E-05 |
| **TAP1** | 1.64987444939749E-09 | -0.14330809498489 | 0.157 | 0.517 | 7.70375876657171E-05 |
| **SLC12A8** | 1.70640054350521E-09 | -0.192014567207158 | 0.21 | 0.586 | 7.96769605778886E-05 |
| **VWA5B2** | 1.7772109256027E-09 | -0.291553495618608 | 0.509 | 0.914 | 8.2983309749167E-05 |
| **MYC** | 1.78638881349251E-09 | -0.107932688457727 | 0.024 | 0.207 | 8.34118528684059E-05 |
| **CTC-444N24** | 1.91087515170736E-09 | -0.125858469700856 | 0.462 | 0.914 | 8.92244934586718E-05 |
| **H2AFJ** | 2.00711554112285E-09 | -0.211863174537317 | 0.754 | 1 | 9.37182459616491E-05 |
| **SH3KBP1** | 2.09175666271885E-09 | -0.183756351206314 | 0.293 | 0.638 | 9.76703938523314E-05 |
| **C15orf48** | 2.21516512476217E-09 | -0.291522944877844 | 0.731 | 0.931 | 0.00010343270517052 |
| **MYH10** | 2.28742576094869E-09 | -0.286108715006257 | 0.506 | 0.879 | 0.000106806771055977 |
| **PPP1R14C** | 2.84509782591482E-09 | -0.106508219555452 | 0.151 | 0.517 | 0.000132846152785441 |
| **GALE** | 2.89664811485462E-09 | -0.149565651760803 | 0.157 | 0.5 | 0.000135253190426907 |
| **HLA-E** | 3.10847460770423E-09 | -0.522511813863464 | 0.76 | 0.983 | 0.000145144004857534 |
| **BCAR3** | 3.22603828468189E-09 | -0.100978786978122 | 0.077 | 0.345 | 0.000150633405626652 |
| **STON2** | 3.37764575960027E-09 | -0.123740864352567 | 0.154 | 0.5 | 0.000157712413453015 |
| **GLUL** | 3.42311472638085E-09 | -0.343776082060121 | 0.456 | 0.879 | 0.000159835495918901 |
| **ARPC1B** | 3.95723482425472E-09 | -0.249041309724347 | 0.68 | 0.983 | 0.000184775165648926 |
| **PLSCR4** | 4.03175353956845E-09 | -0.151121219593357 | 0.062 | 0.31 | 0.00018825466802307 |
| **GSTA1** | 4.16472074658998E-09 | -0.20528436551133 | 0.015 | 0.172 | 0.000194463305820526 |
| **FXYD6** | 4.60413094256416E-09 | -0.328434000091115 | 0.654 | 0.966 | 0.000214980686101148 |
| **EFHD2** | 4.87984423332979E-09 | -0.142414981221338 | 0.373 | 0.793 | 0.000227854566786868 |
| **ARHGEF12** | 5.05948615621549E-09 | -0.136947307204131 | 0.459 | 0.828 | 0.00023624258709217 |
| **PAPSS2** | 5.09403128611394E-09 | -0.467863002148991 | 0.772 | 0.948 | 0.000237855602842518 |
| **TSPAN33** | 5.36953422754553E-09 | -0.11879325870261 | 0.178 | 0.552 | 0.000250719661686783 |
| **RHOB** | 5.41560450520794E-09 | -0.138636857044874 | 0.414 | 0.828 | 0.000252870821161674 |
| **PLP2** | 5.44417609390632E-09 | -0.304274091788441 | 0.849 | 1 | 0.000254204914352768 |
| **AGT** | 5.86579740471473E-09 | -0.594137920582938 | 0.154 | 0.483 | 0.000273891678218345 |
| **MYADM** | 6.44372938352311E-09 | -0.157881954458017 | 0.47 | 0.879 | 0.000300877056104844 |
| **ASXL3** | 6.93794905040986E-09 | -0.190046791177513 | 0.192 | 0.517 | 0.000323953655010787 |

| | | | | | |
|---|---|---|---|---|---|
| **BACE1** | 7.12565672365548E-09 | -0.224520717947887 | 0.314 | 0.707 | 0.000332718289397645 |
| **OCRL** | 7.2760513428205E-09 | -0.170299358160705 | 0.263 | 0.638 | 0.000339740665350318 |
| **FCGRT** | 7.70031414674598E-09 | -0.150409992470174 | 0.393 | 0.845 | 0.00035955076845401 |
| **HIST2H2BE** | 7.77217207250992E-09 | -0.216264159082041 | 0.189 | 0.552 | 0.000362906030581706 |
| **PYGB** | 8.40621845755447E-09 | -0.1996330968829 | 0.429 | 0.897 | 0.000392511558438591 |
| **FAM46A** | 8.53385056536447E-09 | -0.406244824181195 | 0.71 | 1 | 0.000398471084448563 |
| **EHHADH** | 8.56989601084682E-09 | -0.103360155989569 | 0.142 | 0.466 | 0.00040015415443447 |
| **MARVELD3** | 8.70861266602031E-09 | -0.112183022191395 | 0.136 | 0.448 | 0.000406631251214487 |
| **GLYCTK** | 9.64430765191419E-09 | -0.108765726468924 | 0.142 | 0.466 | 0.000450321657190829 |
| **LASP1** | 9.7340684266036E-09 | -0.144968327233973 | 0.417 | 0.862 | 0.000454512857043402 |
| **TIMM21** | 1.01831632723772E-08 | -0.12363884205571 | 0.225 | 0.621 | 0.000475482442677108 |
| **GK** | 1.08982966916375E-08 | -0.102128078999683 | 0.148 | 0.483 | 0.000508874167422629 |
| **FKBP1B** | 1.35091202053972E-08 | -0.178242063310946 | 0.246 | 0.621 | 0.000630781349750613 |
| **DIAPH2** | 1.44386451713142E-08 | -0.103884711519695 | 0.201 | 0.552 | 0.000674183658984173 |
| **NR4A3** | 1.59580435892053E-08 | -0.102576585817068 | 0.115 | 0.414 | 0.000745128929310765 |
| **TMEM164** | 1.70494719775068E-08 | -0.182943714544743 | 0.444 | 0.862 | 0.000796090995045724 |
| **CTNND2** | 1.77400992858302E-08 | -0.10669673453044 | 0.13 | 0.431 | 0.000828338455953268 |
| **DNPH1** | 1.95543182136907E-08 | -0.200463405768276 | 0.55 | 0.914 | 0.000913049780351861 |
| **CDS1** | 1.96636401327111E-08 | -0.125356241771267 | 0.092 | 0.362 | 0.000918154348716681 |
| **DAPK2** | 2.05656792884554E-08 | -0.15174898057899 | 0.34 | 0.759 | 0.000960273263015849 |
| **RASD1** | 2.11209871503004E-08 | 0.522398925990479 | 0.941 | 0.948 | 0.000986202253008976 |
| **NR4A1** | 2.21259395766025E-08 | -0.486406006683489 | 0.467 | 0.845 | 0.0010331264966503 |
| **TMED8** | 2.2884796657327E-08 | -0.177054943231653 | 0.45 | 0.845 | 0.00106855981032057 |
| **LMF2** | 2.31589829519958E-08 | -0.123815830394399 | 0.444 | 0.828 | 0.00108136239097754 |
| **NPTN** | 2.36079156494529E-08 | -0.123985701265639 | 0.769 | 0.983 | 0.0011023244054199 |
| **NFKB1** | 2.73071779751994E-08 | -0.102519885005753 | 0.118 | 0.414 | 0.00127505406119599 |
| **EMC10** | 2.80295297459452E-08 | -0.173532899139642 | 0.775 | 0.948 | 0.00130878283242742 |
| **PRR26** | 2.90916136859234E-08 | -0.134915270001568 | 0.163 | 0.5 | 0.00135837471783682 |
| **TNFSF10** | 3.24326112380958E-08 | -0.147183072385873 | 0.101 | 0.362 | 0.00151437591654041 |
| **KIAA1671** | 3.63488430141587E-08 | -0.129567234667114 | 0.166 | 0.483 | 0.00169723652686011 |
| **DOPEY2** | 3.70129049628068E-08 | -0.175613290993827 | 0.151 | 0.448 | 0.00172824357142834 |
| **UBR5** | 3.8669306125893E-08 | -0.194151588222289 | 0.42 | 0.793 | 0.00180558591093632 |
| **KCTD14** | 4.00777786282885E-08 | -0.138990409327964 | 0.089 | 0.362 | 0.00187135171749068 |
| **TP53I13** | 4.4302303776002E-08 | -0.200551715983051 | 0.512 | 0.879 | 0.00206860747021286 |
| **MCOLN3** | 4.62378507212202E-08 | -0.264808479426642 | 0.358 | 0.741 | 0.00215898396372594 |
| **DSG2** | 4.68780360668563E-08 | -0.156422748464921 | 0.314 | 0.707 | 0.00218887613806972 |

| | | | | | |
|---|---|---|---|---|---|
| **EML5** | 4.79044912938896E-08 | -0.103556827870302 | 0.175 | 0.5 | 0.00223680441198559 |
| **ST3GAL1** | 4.81533509717559E-08 | -0.108303107769654 | 0.204 | 0.552 | 0.0022484244169242 |
| **SESN3** | 4.98181507785888E-08 | -0.171606083355781 | 0.219 | 0.552 | 0.00232615891430465 |
| **PLEKHG3** | 6.00743064173723E-08 | -0.139922035471639 | 0.305 | 0.672 | 0.00280504958954636 |
| **TNFRSF19** | 6.43611713963564E-08 | -0.11362999567267 | 0.154 | 0.483 | 0.00300521617601007 |
| **VAMP8** | 6.64063583717652E-08 | -0.202653255461897 | 0.666 | 1 | 0.00310071209145283 |
| **ARNT2** | 6.90816456362018E-08 | -0.208758648424358 | 0.243 | 0.586 | 0.00322562927969117 |
| **KCNMB2** | 6.98005127938846E-08 | -0.114311891616276 | 0.47 | 0.879 | 0.00325919534388485 |
| **PEG10** | 7.93597662368217E-08 | -0.408847312198185 | 0.74 | 0.897 | 0.00370554556489592 |
| **GRB14** | 8.1984766551191E-08 | -0.138825432304784 | 0.145 | 0.448 | 0.00382811470457476 |
| **HN1L** | 1.03873775826628E-07 | -0.118317644000448 | 0.293 | 0.672 | 0.00485017821467275 |
| **NEURL1** | 1.04995079497661E-07 | -0.23842517130762 | 0.657 | 1 | 0.00490253524698431 |
| **SACM1L** | 1.13417953746446E-07 | -0.132647191079838 | 0.456 | 0.845 | 0.00529582451428282 |
| **GNAZ** | 1.14069395871701E-07 | -0.191426050955328 | 0.441 | 0.793 | 0.00532624230143734 |
| **MCU** | 1.15371307421182E-07 | -0.161900488097221 | 0.281 | 0.638 | 0.00538703245741726 |
| **GNAS** | 1.28026947892289E-07 | 0.347192380960224 | 1 | 1 | 0.00597796227793463 |
| **GRAMD3** | 1.29505051223975E-07 | -0.141523044560302 | 0.367 | 0.776 | 0.00604697935680106 |
| **AMDHD2** | 1.3219678128391E-07 | -0.114426943824836 | 0.314 | 0.741 | 0.00617266430848961 |
| **IGFBP7** | 1.44269257554261E-07 | -0.280569165832853 | 0.651 | 0.966 | 0.00673636444298112 |
| **ABCB1** | 1.44921680986749E-07 | -0.133560116735061 | 0.186 | 0.5 | 0.00676682805031427 |
| **CHKA** | 1.4564063220115E-07 | -0.15406498541087 | 0.328 | 0.741 | 0.00680039803936832 |
| **COX6A1** | 1.47561042083424E-07 | -0.216855813962592 | 0.985 | 1 | 0.00689006773800132 |
| **PPP1R1A** | 1.59969910143257E-07 | -0.291481652134913 | 0.74 | 0.983 | 0.00746947501431911 |
| **SEZ6L2** | 1.6231055971281E-07 | -0.249362471189979 | 0.888 | 1 | 0.00757876696467023 |
| **RSAD2** | 1.63876206722574E-07 | -0.376709486174488 | 0.124 | 0.379 | 0.00765187172049713 |
| **YBX1** | 1.64329719017948E-07 | -0.168753772926506 | 0.973 | 1 | 0.00767304757010506 |
| **UCP2** | 1.78046711747336E-07 | -0.250776432540422 | 0.219 | 0.552 | 0.00831353511161836 |
| **TTC9C** | 1.79788239315323E-07 | -0.114884991172828 | 0.399 | 0.776 | 0.00839485225835038 |
| **CTNNAL1** | 1.87813052285087E-07 | -0.154108594507353 | 0.29 | 0.672 | 0.00876955485034755 |
| **STMN2** | 1.91169160893281E-07 | -0.142071790975694 | 0.444 | 0.862 | 0.00892626162958996 |
| **MBOAT2** | 1.97157230825234E-07 | -0.172318434807909 | 0.719 | 0.931 | 0.00920586257892266 |
| **GRHL1** | 2.0102778409842E-07 | -0.114832582901046 | 0.189 | 0.5 | 0.00938659032290753 |
| **KIF9** | 2.08056746008948E-07 | -0.12145045942345 | 0.296 | 0.638 | 0.0097147936413958 |
| **PLXNB1** | 2.08217207364173E-07 | -0.140204293667604 | 0.24 | 0.569 | 0.00972228606345533 |
| **SMIM22** | 2.15597248869746E-07 | -0.129676935790038 | 0.574 | 0.931 | 0.0100668823414751 |
| **ATP9A** | 2.33180712978227E-07 | -0.277413419359121 | 0.722 | 0.931 | 0.0108879070310924 |

| | | | | | |
|---|---|---|---|---|---|
| **PHLDB2** | 2.41198723244494E-07 | -0.124656876167032 | 0.364 | 0.724 | 0.0112622919844552 |
| **NDUFB2** | 2.45330787866454E-07 | -0.221312668687024 | 0.973 | 1 | 0.0114552304778484 |
| **BIVM** | 2.46493003105568E-07 | -0.114077112862885 | 0.225 | 0.569 | 0.0115094977940083 |
| **MOB1B** | 2.49862024370355E-07 | -0.209421786451926 | 0.482 | 0.879 | 0.011666807503925 |
| **MOV10** | 2.64884015614465E-07 | -0.124291414526098 | 0.151 | 0.448 | 0.0123682293410862 |
| **NT5C3A** | 2.68631883358228E-07 | -0.196923491509114 | 0.538 | 0.897 | 0.0125432285296457 |
| **PTP4A3** | 2.80546964796952E-07 | -0.115563797653088 | 0.254 | 0.621 | 0.0130995794272641 |
| **PRKD1** | 2.81598815811443E-07 | -0.191337667169399 | 0.104 | 0.345 | 0.0131486935066837 |
| **GNB5** | 3.03104865929217E-07 | -0.11140322902744 | 0.417 | 0.81 | 0.0141528755048329 |
| **MBD3** | 3.09557212679126E-07 | -0.12476098592808 | 0.284 | 0.638 | 0.0144541549316264 |
| **P3H4** | 3.1494536150294E-07 | -0.101172697034606 | 0.287 | 0.655 | 0.0147057437646568 |
| **PFN1** | 3.17446806522462E-07 | -0.244853530921768 | 0.982 | 1 | 0.0148225437369533 |
| **NR4A2** | 3.18182906753136E-07 | -0.361390621688193 | 0.426 | 0.741 | 0.0148569144650242 |
| **DIAPH1** | 3.22663531673851E-07 | -0.109787568464374 | 0.45 | 0.828 | 0.0150661282844471 |
| **CADM1** | 3.54764825002678E-07 | 0.497094391692175 | 0.962 | 1 | 0.01656503397385 |
| **WEE1** | 3.56884989080172E-07 | -0.105923305385492 | 0.13 | 0.397 | 0.0166640307951205 |
| **NAA20** | 3.64029925018139E-07 | -0.317060583367259 | 0.953 | 1 | 0.016997649288872 |
| **CFC1** | 3.66569897549492E-07 | -0.304835701816531 | 0.962 | 1 | 0.0171162482262784 |
| **DIP2B** | 3.76244934267778E-07 | -0.168173222383796 | 0.467 | 0.862 | 0.0175680047157654 |
| **CALY** | 3.76655770190819E-07 | -0.141320524385706 | 0.299 | 0.638 | 0.0175871878775199 |
| **ABLIM1** | 3.81265299813698E-07 | -0.325996599663429 | 0.754 | 0.983 | 0.017802420644201 |
| **SPINT2** | 3.86349664229429E-07 | -0.299052974014658 | 0.97 | 1 | 0.0180398248718647 |
| **LINC00888** | 3.97782147389785E-07 | -0.141835566153836 | 0.257 | 0.603 | 0.0185736418080712 |
| **PLOD3** | 4.00633394547102E-07 | -0.252903942253531 | 0.577 | 0.914 | 0.0187067750915878 |
| **CFL1** | 4.09738601933404E-07 | -0.229589535209892 | 0.973 | 1 | 0.0191319245400765 |
| **DMKN** | 4.16254745691163E-07 | -0.168924411688691 | 0.574 | 0.879 | 0.0194361828405575 |
| **NRIP1** | 4.28324814327147E-07 | -0.12787475541324 | 0.293 | 0.655 | 0.0199997705553775 |
| **TMEM131** | 4.41238669711374E-07 | -0.116535920458659 | 0.429 | 0.793 | 0.0206027572048332 |
| **SH3BGRL3** | 4.43449748348807E-07 | -0.22315191325132 | 0.843 | 1 | 0.0207059990996508 |
| **ENPP1** | 4.99715489673902E-07 | -0.231407631511114 | 0.334 | 0.672 | 0.0233332153593435 |
| **GCH1** | 5.10986249889351E-07 | -0.404831966230605 | 0.675 | 0.966 | 0.0238594809660835 |
| **MGAT4B** | 5.43579249142927E-07 | -0.138562318700926 | 0.308 | 0.638 | 0.0253813458802307 |
| **CALM3** | 5.5621229301525E-07 | -0.212591984285135 | 0.911 | 1 | 0.0259712205977611 |
| **AKAP12** | 5.5932388480178E-07 | -0.164857720230966 | 0.234 | 0.569 | 0.0261165101530495 |
| **TXNDC11** | 6.19035963137248E-07 | -0.240243334500087 | 0.476 | 0.897 | 0.0289046462267675 |
| **PLS1** | 6.36308680405771E-07 | -0.189862707557231 | 0.503 | 0.879 | 0.0297111612141867 |

| | | | | | |
|---|---|---|---|---|---|
| **ZDHHC24** | 6.77338923467586E-07 | -0.108329130879533 | 0.432 | 0.81 | 0.031626986353472 |
| **TJP1** | 6.81706683126925E-07 | -0.15427012809945 | 0.58 | 0.862 | 0.0318309301552455 |
| **CAPZB** | 6.88604738137365E-07 | -0.250377688611913 | 0.893 | 0.983 | 0.032153021037848 |
| **SERPINB1** | 7.13819296835243E-07 | -0.200226193788631 | 0.568 | 0.931 | 0.033330364427128 |
| **KLHL18** | 7.29629760829587E-07 | -0.111320051009394 | 0.328 | 0.655 | 0.0340686024224159 |
| **SLC25A33** | 7.42412944514258E-07 | -0.184676558802427 | 0.515 | 0.879 | 0.0346654876182043 |
| **ODF2L** | 7.56265530057886E-07 | -0.146658437018769 | 0.361 | 0.724 | 0.0353123063949929 |
| **XPO4** | 7.74010836152443E-07 | -0.120082218635415 | 0.201 | 0.5 | 0.036140887972466 |
| **FBP1** | 7.89531059264008E-07 | -0.140707114060941 | 0.095 | 0.328 | 0.0368655737502143 |
| **SLC37A4** | 7.94384372200515E-07 | -0.290409094778938 | 0.589 | 0.948 | 0.0370921894911587 |
| **AURKAIP1** | 8.13265795103749E-07 | -0.202845922693643 | 0.873 | 1 | 0.0379738197707793 |
| **FLNB** | 8.23034679319505E-07 | -0.211925697457228 | 0.683 | 0.914 | 0.0384299582814657 |
| **TTC39A** | 8.53194414100877E-07 | -0.282790329804557 | 0.763 | 0.983 | 0.0398382067776122 |
| **ALDH2** | 8.70744224612969E-07 | -0.156203990985199 | 0.42 | 0.81 | 0.0406576600798534 |
| **RHPN1** | 9.16499669684211E-07 | -0.135496716104027 | 0.234 | 0.534 | 0.0427941190765649 |
| **AAK1** | 9.91382101290939E-07 | -0.206756978742736 | 0.743 | 0.914 | 0.0462906044555778 |
| **LPGAT1** | 9.92264650801061E-07 | -0.173321434504753 | 0.385 | 0.741 | 0.046331813339854 |
| **SERF2** | 9.93153045595476E-07 | -0.234175466263683 | 1 | 1 | 0.0463732951579896 |
| **KRT19** | 1.0224960371328E-06 | -0.140835481082231 | 0.18 | 0.466 | 0.0477434074618419 |
| **PPP2R5B** | 1.03687738434844E-06 | -0.105518517857638 | 0.364 | 0.707 | 0.0484149157073819 |
| **AP1S2** | 1.03768116204757E-06 | -0.141814502827749 | 0.263 | 0.603 | 0.0484524464994874 |
| **TTN** | 1.04366996869632E-06 | -0.246123360102898 | 0.456 | 0.707 | 0.0487320818483373 |
| **CDH1** | 1.05426195445564E-06 | -0.288463409056408 | 0.802 | 1 | 0.0492266534393972 |

Supplemental Table 2 - Cluster1_vs_cluster7 in non-diabetic donors

| | p_val | avg_logFC | pct.1 | pct.2 | p_val_adj |
|---|---|---|---|---|---|
| ARX | 1.09197249203384E-46 | -0.277871654929122 | 0.038 | 0.927 | 5.09874715705362E-42 |
| MUC13 | 3.54604027994565E-43 | -0.729527214368204 | 0.023 | 0.78 | 1.65575258791502E-38 |
| GC | 1.17095638508797E-42 | -1.95486001803184 | 0.103 | 1 | 5.46754664889126E-38 |
| TMEM236 | 2.67446207912997E-42 | -0.514893160898948 | 0.046 | 0.854 | 1.24878657860816E-37 |
| GPR119 | 4.56722513249049E-42 | -0.34733736876347 | 0.038 | 0.854 | 2.13257443111378E-37 |
| LOXL4 | 1.18397036046967E-39 | -2.25256357156365 | 0.118 | 0.976 | 5.52831280414102E-35 |
| TM4SF4 | 3.96129076430817E-38 | -2.28304928672284 | 0.156 | 1 | 1.84964549657841E-33 |
| DPP4 | 1.77704958131889E-36 | -0.641675460250022 | 0.046 | 0.78 | 8.29757761005228E-32 |
| MRC1 | 1.96481286363094E-36 | -0.860553353885405 | 0.065 | 0.829 | 9.17430070415195E-32 |
| SPOCK3 | 2.578248510339E-36 | -0.43534840140745 | 0.042 | 0.756 | 1.20386157693259E-31 |
| F10 | 4.77491417264933E-35 | -0.794426230400762 | 0.118 | 0.951 | 2.22955067463515E-30 |
| KCTD12 | 4.31489070069768E-34 | -0.716035160752271 | 0.148 | 0.976 | 2.01475191487677E-29 |
| DMD | 1.3256297072646E-33 | -0.18467728337312 | 0.049 | 0.756 | 6.1897627921306E-29 |
| SERPINA1 | 1.78507026563048E-33 | -1.62645822835248 | 0.068 | 0.805 | 8.33502859130838E-29 |
| POPDC3 | 3.21906293995569E-33 | -0.249985860161138 | 0.011 | 0.585 | 1.50307705855351E-28 |
| PAPPA2 | 3.60396111535254E-33 | -0.507178475175219 | 0.042 | 0.707 | 1.68279756359156E-28 |
| CRH | 4.28722200595811E-33 | -0.794154046558248 | 0.027 | 0.659 | 2.00183257124202E-28 |
| FXYD3 | 1.21644618932753E-31 | -0.435267136539163 | 0.095 | 0.854 | 5.67995219182705E-27 |
| MAOB | 2.16464794578442E-31 | -0.343955793561699 | 0.038 | 0.683 | 1.01073906532512E-26 |
| CD36 | 7.47165561139226E-31 | -0.693879612735007 | 0.042 | 0.683 | 3.48874015462739E-26 |
| APOH | 1.8652322468258E-30 | -0.539436835243995 | 0.042 | 0.683 | 8.7093289301037E-26 |
| NPNT | 4.55084620680307E-30 | -0.462189714048027 | 0.038 | 0.659 | 2.12492661934256E-25 |
| TMEM176A | 8.54662559267864E-30 | -1.11296503419195 | 0.266 | 1 | 3.99067588798944E-25 |
| IRX2 | 1.23048652101768E-29 | -0.361936096780119 | 0.091 | 0.78 | 5.74551071258785E-25 |
| PTPRT | 1.91119979613663E-29 | -0.309516438810408 | 0.034 | 0.634 | 8.92396520810078E-25 |
| VIL1 | 2.79626808246183E-29 | -0.587710173865183 | 0.046 | 0.659 | 1.3056614557439E-24 |
| S100A10 | 1.56300578181929E-28 | -0.512861515473964 | 0.125 | 0.878 | 7.29814289704882E-24 |
| VIM | 1.38001570539838E-27 | -1.27224541585328 | 0.179 | 0.902 | 6.44370733321666E-23 |
| GPC6 | 2.19422195975949E-27 | -0.347962973452563 | 0.057 | 0.683 | 1.0245480596705E-22 |
| TMEM86B | 7.83142104528287E-27 | -0.232712488995468 | 0.084 | 0.756 | 3.65672542867393E-22 |
| MYO10 | 1.05673912063734E-25 | -0.664384987355504 | 0.072 | 0.683 | 4.93423197599195E-21 |
| FEV | 1.36321905322266E-25 | -0.535760984109832 | 0.171 | 0.878 | 6.36527872521257E-21 |
| FAP | 2.64540051270197E-25 | -0.624364897052682 | 0.103 | 0.732 | 1.23521686139593E-20 |
| FABP5 | 5.05170998566815E-25 | -0.399613381296317 | 0.144 | 0.829 | 2.35879494360803E-20 |
| TMEM176B | 7.36660488671629E-25 | -1.2669879821745 | 0.35 | 0.976 | 3.43968881975444E-20 |

| | | | | | |
|---|---|---|---|---|---|
| **TTR** | 2.81138127771011E-24 | -2.35718180735035 | 0.966 | 1 | 1.31271826000118E-19 |
| **LDHA** | 2.35838478817717E-23 | -0.861470690038796 | 0.129 | 0.805 | 1.10120060914356E-18 |
| **LAMB1** | 3.7389943708131E-23 | -0.438796667863001 | 0.163 | 0.805 | 1.74584864156376E-18 |
| **GCG** | 4.55756888415312E-23 | -3.63312085639101 | 0.741 | 1 | 2.12806563907762E-18 |
| **GPER1** | 7.70124288669531E-23 | -0.194744738062598 | 0.03 | 0.512 | 3.59594134108464E-18 |
| **KCNIP1** | 1.87518599033021E-22 | -0.251767701906357 | 0.03 | 0.512 | 8.75580594464883E-18 |
| **MBOAT4** | 7.35021627388465E-22 | -0.388724340891473 | 0.068 | 0.61 | 3.43203648476496E-17 |
| **KIF12** | 7.77698316583141E-22 | -0.208078061317748 | 0.148 | 0.805 | 3.63130674962166E-17 |
| **IGFBP2** | 8.57407241100904E-22 | -0.853028628749056 | 0.3 | 0.976 | 4.00349163087245E-17 |
| **FXYD5** | 1.89002349521137E-21 | -0.55721307104222 | 0.156 | 0.805 | 8.82508670619047E-17 |
| **SPTSSB** | 2.28694227699057E-21 | -0.600845534113068 | 0.129 | 0.78 | 1.06784195739521E-16 |
| **ADGRG2** | 2.28891750319062E-21 | -0.476459931722775 | 0.114 | 0.707 | 1.0687642497648E-16 |
| **PTGER4** | 4.68481532591332E-21 | -0.256835086201887 | 0.015 | 0.415 | 2.18748082012871E-16 |
| **PRSS3** | 4.91334596115702E-21 | -0.265181747243205 | 0.034 | 0.488 | 2.29418862964305E-16 |
| **MYRF** | 5.31142518177663E-21 | -0.2051367722462 | 0.072 | 0.61 | 2.48006376012696E-16 |
| **PPP2R2B** | 5.3272134198433E-21 | -0.188521873569351 | 0.038 | 0.512 | 2.48743576212743E-16 |
| **SMIM24** | 5.64090382355403E-21 | -0.68830597057406 | 0.19 | 0.829 | 2.63390722233208E-16 |
| **ADGRG5** | 1.73978017672173E-20 | -0.157517720634252 | 0.046 | 0.537 | 8.12355557916675E-16 |
| **RNF144A** | 4.87381452289715E-20 | -0.137853504283216 | 0.03 | 0.463 | 2.27573021517637E-15 |
| **PLPPR1** | 5.62573557989445E-20 | -0.245408209874634 | 0.03 | 0.463 | 2.62682471432012E-15 |
| **IMPA2** | 7.36345656818335E-20 | -0.187974772410015 | 0.053 | 0.561 | 3.43821877538185E-15 |
| **C1orf21** | 7.69931579666172E-20 | -0.117048668257994 | 0.027 | 0.439 | 3.59504152493526E-15 |
| **KLHL41** | 1.68935279900089E-19 | -0.352461635099542 | 0.072 | 0.585 | 7.88809502437487E-15 |
| **SERPINA10** | 1.74949793432074E-19 | -0.170945394533981 | 0.019 | 0.415 | 8.16893070472382E-15 |
| **CSGALNACT1** | 2.04151892636399E-19 | -0.252695207075237 | 0.061 | 0.561 | 9.53246432287139E-15 |
| **CDO1** | 4.35484309383551E-19 | -0.198215496487403 | 0.008 | 0.341 | 2.03340688580461E-14 |
| **SLC6A19** | 4.46157571103954E-19 | -0.31780973313348 | 0.027 | 0.439 | 2.08324354675569E-14 |
| **ERBB3** | 5.09122567640748E-19 | -0.281814977897713 | 0.057 | 0.537 | 2.37724600508494E-14 |
| **USH1C** | 5.57237154593763E-19 | -0.501514319677441 | 0.179 | 0.805 | 2.60190744594466E-14 |
| **AOX1** | 5.64455283913893E-19 | -0.255660223510757 | 0.034 | 0.463 | 2.63561105717914E-14 |
| **CLU** | 1.02868659957713E-18 | -1.39015239405022 | 0.924 | 1 | 4.80324633940549E-14 |
| **EGFL7** | 1.05466477504998E-18 | -0.248351161046021 | 0.198 | 0.878 | 4.92454623414089E-14 |
| **ANK2** | 1.30974248382676E-18 | -0.222436792181573 | 0.095 | 0.634 | 6.1155805797323E-14 |
| **BAIAP2L2** | 1.74063097868954E-18 | -0.105192472210766 | 0.019 | 0.39 | 8.12752822879509E-14 |
| **PYROXD2** | 2.47459969259787E-18 | -0.127886089047476 | 0.042 | 0.488 | 1.15546483446472E-13 |
| **SYNDIG1** | 3.77109215615861E-18 | -0.319030338986598 | 0.049 | 0.512 | 1.76083606047514E-13 |
| **RHOC** | 4.87380412222664E-18 | -0.271008573741462 | 0.091 | 0.61 | 2.27572535879129E-13 |

| | | | | | |
|---|---|---|---|---|---|
| **CRYBA2** | 5.17445874218837E-18 | -1.69468159929806 | 0.715 | 1 | 2.41611002049002E-13 |
| **PLIN3** | 6.98068450156183E-18 | -0.203603078058187 | 0.091 | 0.634 | 3.25949101431427E-13 |
| **PLCE1** | 1.53795288380882E-17 | -0.832250626973747 | 0.179 | 0.78 | 7.18116340036854E-13 |
| **LDB2** | 1.65857410797108E-17 | -0.185965063815629 | 0.023 | 0.39 | 7.74438008234937E-13 |
| **SLC38A4** | 1.68546348073132E-17 | -0.38392926417237 | 0.209 | 0.829 | 7.86993463057874E-13 |
| **TPD52L1** | 1.73175862253118E-17 | -0.274965015539831 | 0.057 | 0.512 | 8.08610053618485E-13 |
| **SPATS2L** | 1.01579781097038E-16 | -0.318829393730951 | 0.194 | 0.78 | 4.74306471876398E-12 |
| **FAM110B** | 1.4266057655838E-16 | -0.291656344303343 | 0.163 | 0.707 | 6.66125030124045E-12 |
| **RGS4** | 1.96267377395704E-16 | -0.975282252691519 | 0.38 | 0.927 | 9.16431265273762E-12 |
| **SLC44A3** | 2.18425532874763E-16 | -0.147119230780063 | 0.038 | 0.439 | 1.01989434065213E-11 |
| **CNIH3** | 2.64385376020719E-16 | -0.226128540598591 | 0.099 | 0.61 | 1.23449463625354E-11 |
| **KCNJ3** | 2.6638413413849E-16 | -0.105537831234942 | 0.106 | 0.61 | 1.24382743753285E-11 |
| **CYP4F3** | 3.09092722856259E-16 | -0.158179570254514 | 0.015 | 0.341 | 1.44324665083273E-11 |
| **SEMA3E** | 4.7507884830171E-16 | -0.154247448866643 | 0.038 | 0.415 | 2.21828566637518E-11 |
| **INHBA** | 5.34380143347671E-16 | -0.273838433845322 | 0.167 | 0.756 | 2.49518120333328E-11 |
| **KCNH6** | 6.75162098213997E-16 | -0.26877542956871 | 0.163 | 0.732 | 3.15253438519062E-11 |
| **STC2** | 8.46462050114399E-16 | -0.108697867282178 | 0.023 | 0.366 | 3.95238525059916E-11 |
| **DOK4** | 1.00715121373177E-15 | -0.253561558084824 | 0.125 | 0.683 | 4.70269116227774E-11 |
| **REG4** | 1.19036634707466E-15 | -0.571136653671733 | 0.038 | 0.415 | 5.55817758439572E-11 |
| **CD82** | 1.24860176000098E-15 | -0.496524771468627 | 0.281 | 0.854 | 5.83009619797258E-11 |
| **HS3ST3B1** | 2.45775301542044E-15 | -0.187226587832533 | 0.046 | 0.439 | 1.14759861549026E-10 |
| **TMEM45B** | 3.39332074251564E-15 | -0.189542407689954 | 0.027 | 0.366 | 1.58444325430283E-10 |
| **CYSTM1** | 3.98005347351974E-15 | -0.697709836786659 | 0.928 | 1 | 1.85840636839057E-10 |
| **ANXA4** | 4.24864248672516E-15 | -0.369643656188407 | 0.202 | 0.805 | 1.98381863632658E-10 |
| **ANG** | 6.03437493620277E-15 | -0.128300124972294 | 0.027 | 0.366 | 2.81763068896116E-10 |
| **GLS** | 7.75375847624672E-15 | -0.599130052495756 | 0.51 | 1 | 3.62046244531388E-10 |
| **PHLDA1** | 8.99802980777896E-15 | -0.102973631073992 | 0.042 | 0.415 | 4.20145005814623E-10 |
| **QDPR** | 9.78359868012315E-15 | -0.809386335380637 | 0.764 | 0.976 | 4.5682557317099E-10 |
| **PDK4** | 1.54659814620421E-14 | -1.18727759057498 | 0.483 | 0.976 | 7.2215307240713E-10 |
| **CAMK2G** | 1.75859959199303E-14 | -0.552092262008913 | 0.342 | 0.902 | 8.21142907489305E-10 |
| **CEACAM1** | 2.27404320599728E-14 | -0.297387857486494 | 0.118 | 0.61 | 1.06181899417631E-09 |
| **TSPAN14** | 2.61696485766585E-14 | -0.23255180202725 | 0.171 | 0.683 | 1.22193940098991E-09 |
| **ARFGEF3** | 2.90408816588797E-14 | -0.772149887731609 | 0.833 | 1 | 1.35600588729807E-09 |
| **HMGB3** | 3.84694995092838E-14 | -0.460057677115632 | 0.308 | 0.854 | 1.79625634058699E-09 |
| **INS** | 4.13700359977703E-14 | 0.739562751473335 | 1 | 1 | 1.93169109084389E-09 |
| **ABHD15** | 4.40584383631788E-14 | -0.254458169755519 | 0.179 | 0.732 | 2.05722066249191E-09 |
| **NEDD9** | 4.81866523246377E-14 | -0.307349952495612 | 0.137 | 0.634 | 2.24997935699431E-09 |

| | | | | | |
|---|---|---|---|---|---|
| **NPHP4** | 7.60282257593833E-14 | -0.125520207957072 | 0.091 | 0.537 | 3.54998594538289E-09 |
| **SLC7A14** | 9.3481308609197E-14 | -0.334370139844006 | 0.228 | 0.78 | 4.36492274288924E-09 |
| **COTL1** | 9.61696483325965E-14 | -0.503774465532594 | 0.357 | 0.927 | 4.49044938959393E-09 |
| **COL27A1** | 1.02358629814716E-13 | -0.159327370075338 | 0.129 | 0.61 | 4.77943150193852E-09 |
| **DSC2** | 1.04743651655295E-13 | -0.165656706090891 | 0.065 | 0.463 | 4.89079532674068E-09 |
| **ITPR1** | 1.62330007689576E-13 | -0.160981223441537 | 0.137 | 0.634 | 7.57967504904935E-09 |
| **CPNE8** | 2.41770542164221E-13 | -0.295688025308007 | 0.232 | 0.829 | 1.1288991925274E-08 |
| **SMOC1** | 2.55679383217687E-13 | -0.239898347330402 | 0.141 | 0.634 | 1.19384374405834E-08 |
| **EZR** | 4.42572457213289E-13 | -0.622135546868859 | 0.73 | 1 | 2.06650357446601E-08 |
| **PRLR** | 5.16451862637395E-13 | -0.162926982353382 | 0.038 | 0.366 | 2.41146868221279E-08 |
| **PALLD** | 5.32745978795773E-13 | -0.681051615896128 | 0.483 | 0.951 | 2.4875507987911E-08 |
| **STK32B** | 5.4183420098615E-13 | -0.120449582334327 | 0.004 | 0.22 | 2.52998643466463E-08 |
| **GALNT13** | 5.43460166526879E-13 | -0.238026417027233 | 0.084 | 0.488 | 2.53757855556396E-08 |
| **GRIA3** | 9.46405991760848E-13 | -0.484137312044005 | 0.414 | 0.927 | 4.41905349732893E-08 |
| **TMPRSS2** | 9.80166388520962E-13 | -0.150876553418942 | 0.019 | 0.293 | 4.57669091792093E-08 |
| **MARCKSL1** | 1.08377261506157E-12 | -0.17746993590967 | 0.129 | 0.585 | 5.060459471507E-08 |
| **HNF1A-AS1** | 1.25983460356451E-12 | -0.124204754479375 | 0.095 | 0.512 | 5.88254571442378E-08 |
| **CBR3-AS1** | 1.48260604950623E-12 | -0.166984803188206 | 0.202 | 0.732 | 6.92273242695943E-08 |
| **CRTAC1** | 2.16965673703845E-12 | -0.108879351615308 | 0.027 | 0.317 | 1.01307782022536E-07 |
| **ELL2** | 2.28276105889272E-12 | -0.50730952289018 | 0.753 | 1 | 1.06588962122878E-07 |
| **ARRDC4** | 2.30831985299838E-12 | -0.291878064326422 | 0.133 | 0.585 | 1.07782378896053E-07 |
| **PLXDC2** | 2.48609234270672E-12 | -0.166407991521953 | 0.061 | 0.415 | 1.16083109758005E-07 |
| **SLC5A1** | 2.75472950179752E-12 | -0.191910572257982 | 0.065 | 0.439 | 1.28626584627432E-07 |
| **SMIM5** | 2.79068742606682E-12 | -0.157428448718269 | 0.084 | 0.488 | 1.30305567985338E-07 |
| **SH3RF1** | 3.22376079743949E-12 | -0.193210902497198 | 0.11 | 0.537 | 1.50527062914842E-07 |
| **AC009014.3** | 3.42472638830219E-12 | -0.146196585957469 | 0.08 | 0.488 | 1.59910749248994E-07 |
| **MYC** | 4.8644381571417E-12 | -0.143860327051963 | 0.011 | 0.244 | 2.27135210871417E-07 |
| **SAMD4A** | 5.52649014160829E-12 | -0.102019422112676 | 0.091 | 0.488 | 2.58048404182116E-07 |
| **UPB1** | 5.94862103671606E-12 | -0.106701188712368 | 0.023 | 0.293 | 2.77758962067383E-07 |
| **TM4SF1** | 6.19545216065289E-12 | -0.318158271545189 | 0.046 | 0.366 | 2.89284247737365E-07 |
| **ZFP36** | 6.82403601110886E-12 | -0.272542488606496 | 0.221 | 0.732 | 3.18634713466706E-07 |
| **ELF3** | 9.01901605623717E-12 | -0.392737846984139 | 0.316 | 0.829 | 4.21124916713882E-07 |
| **BID** | 9.1317574112332E-12 | -0.301023583595586 | 0.221 | 0.683 | 4.26389148802712E-07 |
| **FAM131C** | 1.07540174095248E-11 | -0.119665714897166 | 0.24 | 0.756 | 5.02137334902943E-07 |
| **TSPAN15** | 1.34232885955541E-11 | -0.109932911886641 | 0 | 0.171 | 6.26773614392208E-07 |
| **CACNB2** | 1.37075861646438E-11 | -0.463643714669892 | 0.51 | 0.927 | 6.40048320785713E-07 |
| **LBH** | 1.46321145454476E-11 | -0.15985711838181 | 0.065 | 0.415 | 6.83217324470585E-07 |

| | | | | | |
|---|---|---|---|---|---|
| **LMCD1** | 1.72734646916787E-11 | -0.324151833035828 | 0.114 | 0.537 | 8.06549886848555E-07 |
| **HIGD1A** | 1.7462741495809E-11 | -0.59090668994872 | 0.749 | 1 | 8.15387788663811E-07 |
| **CLSTN2** | 2.28823699655807E-11 | -0.12945275724206 | 0.087 | 0.488 | 1.06844650080286E-06 |
| **MFSD3** | 2.28950795807326E-11 | -0.231873660512986 | 0.323 | 0.854 | 1.06903995086315E-06 |
| **FOSL2** | 3.33703616313877E-11 | -0.227230145480033 | 0.262 | 0.78 | 1.55816229565439E-06 |
| **SLC6A4** | 3.50607642810009E-11 | -0.318120820548224 | 0.175 | 0.634 | 1.63709226657277E-06 |
| **SERTM1** | 3.59742075194379E-11 | -0.106705433139636 | 0.019 | 0.268 | 1.67974367170512E-06 |
| **FAM83F** | 3.83927413997256E-11 | -0.137569363361111 | 0.209 | 0.707 | 1.79267227417739E-06 |
| **SSX2IP** | 4.83116981480152E-11 | -0.236896907177278 | 0.312 | 0.902 | 2.25581812162527E-06 |
| **TMEM56** | 4.91411608852075E-11 | -0.114198859268524 | 0.072 | 0.415 | 2.29454822521299E-06 |
| **SLC40A1** | 5.62082025086462E-11 | -0.574822027585317 | 0.27 | 0.756 | 2.62452959973622E-06 |
| **GLRX** | 7.05230380866557E-11 | -0.238813583624139 | 0.259 | 0.756 | 3.29293221738021E-06 |
| **VGF** | 7.50412997130591E-11 | -0.282416309530914 | 0.76 | 1 | 3.50390340750187E-06 |
| **ANKS4B** | 8.70975120853141E-11 | -0.102363349004849 | 0.015 | 0.244 | 4.06684413179957E-06 |
| **SNED1** | 8.78767390096972E-11 | -0.119107540771639 | 0.179 | 0.634 | 4.10322857457979E-06 |
| **SLC7A8** | 9.5370134583673E-11 | -0.666932215981683 | 0.935 | 1 | 4.45311769411544E-06 |
| **ALDH1A1** | 9.93525855556748E-11 | -0.787924828669962 | 0.962 | 1 | 4.63907027735113E-06 |
| **KCNA5** | 1.06018565200326E-10 | -0.23950315466893 | 0.167 | 0.634 | 4.9503248648988E-06 |
| **HACD1** | 1.1016780053355E-10 | -0.207828186729001 | 0.278 | 0.854 | 5.14406511031305E-06 |
| **HGD** | 1.13492646567457E-10 | -0.120089157237575 | 0.065 | 0.415 | 5.29931214617427E-06 |
| **TCN2** | 1.45580697763876E-10 | -0.10202075703251 | 0.087 | 0.463 | 6.79759952068865E-06 |
| **GSTA1** | 1.51443438075118E-10 | -0.26264643539116 | 0.011 | 0.22 | 7.07134845404148E-06 |
| **GPX3** | 1.57528320441873E-10 | -0.674738708339716 | 0.795 | 1 | 7.35546986639239E-06 |
| **AGFG2** | 1.77484297843218E-10 | -0.117365091744067 | 0.19 | 0.683 | 8.28727431919338E-06 |
| **PDE3B** | 1.96446259026642E-10 | -0.252714682512443 | 0.392 | 0.902 | 9.172665172731E-06 |
| **GNG5** | 1.99633928566615E-10 | -0.344019038262573 | 0.608 | 1 | 9.32150702656094E-06 |
| **CASP7** | 2.25243158921532E-10 | -0.103070453084135 | 0.065 | 0.415 | 1.05172788195231E-05 |
| **ITGB8** | 2.26125060133523E-10 | -0.11459460115588 | 0.114 | 0.512 | 1.05584574328146E-05 |
| **RPS6KA3** | 2.28952591605475E-10 | -0.393508399791795 | 0.452 | 0.902 | 1.06904833598344E-05 |
| **KLB** | 2.41542292261994E-10 | -0.135869550622502 | 0.038 | 0.317 | 1.12783342525893E-05 |
| **DNPH1** | 3.15846633484246E-10 | -0.29849253940177 | 0.525 | 0.951 | 1.47478268572799E-05 |
| **STXBP6** | 3.38062221884351E-10 | -0.147316344072321 | 0.087 | 0.439 | 1.5785139326446E-05 |
| **FSTL5** | 3.99019043218871E-10 | -0.125066002620556 | 0.114 | 0.512 | 1.86313961850187E-05 |
| **LINC00473** | 4.80352963362671E-10 | -0.428271405527336 | 0.266 | 0.732 | 2.24291209182932E-05 |
| **NT5E** | 6.44915867126096E-10 | -0.102134608187602 | 0.019 | 0.244 | 3.01130565837188E-05 |
| **HES6** | 6.68868314877194E-10 | -0.104183833891095 | 0.137 | 0.561 | 3.12314682265608E-05 |
| **AC004381.6** | 7.32394819009802E-10 | -0.110132225097592 | 0.049 | 0.341 | 3.41977112840247E-05 |

| | | | | | |
|---|---|---|---|---|---|
| **MATN2** | 7.34504376739596E-10 | -0.22686740952072 | 0.198 | 0.659 | 3.4296212863102E-05 |
| **MYL6** | 7.43879926709944E-10 | -0.368676803209358 | 1 | 1 | 3.47339854178674E-05 |
| **WIPF3** | 7.44511697479047E-10 | -0.138066834757602 | 0.042 | 0.317 | 3.47634846903891E-05 |
| **MPC2** | 8.08594532028106E-10 | -0.475892415330123 | 0.829 | 0.976 | 3.77557044839883E-05 |
| **LASP1** | 8.31642579085138E-10 | -0.183338212334831 | 0.38 | 0.927 | 3.88318869452223E-05 |
| **TTLL12** | 1.00303697768157E-09 | -0.144229455668505 | 0.198 | 0.659 | 4.68348055988856E-05 |
| **NR4A1** | 1.08802584540565E-09 | -0.580735701788959 | 0.441 | 0.927 | 5.0803190799526E-05 |
| **TAGLN2** | 1.10705218782842E-09 | -0.600801564174882 | 0.878 | 1 | 5.16915878062726E-05 |
| **PCSK2** | 1.20927804919031E-09 | -0.611034120973776 | 0.924 | 1 | 5.64648199508433E-05 |
| **HPS1** | 1.30784382305569E-09 | -0.505859204980939 | 0.582 | 0.902 | 6.10671516299392E-05 |
| **TMSB4X** | 1.42154865348093E-09 | -0.653223343061127 | 0.878 | 1 | 6.63763712769849E-05 |
| **CHGA** | 1.48782130308949E-09 | -0.605802344703298 | 0.985 | 1 | 6.94708401051575E-05 |
| **LUZP1** | 1.50505222806826E-09 | -0.170692642683637 | 0.114 | 0.488 | 7.02754036851914E-05 |
| **MYCBP** | 1.56488828559099E-09 | -0.132240451291295 | 0.148 | 0.561 | 7.30693287191003E-05 |
| **BTBD11** | 1.62397872474454E-09 | -0.125991251628373 | 0.072 | 0.39 | 7.58284385944968E-05 |
| **RHOB** | 2.06857111512555E-09 | -0.182117132648783 | 0.411 | 0.902 | 9.65877910785574E-05 |
| **SERPINE2** | 2.13000711445795E-09 | -0.391418478657302 | 0.38 | 0.902 | 9.94564221953852E-05 |
| **LIMA1** | 2.25536736586543E-09 | -0.19155168383418 | 0.338 | 0.854 | 0.000105309868414355 |
| **A1CF** | 2.46363753374984E-09 | -0.260863116863529 | 0.3 | 0.756 | 0.000115034627363381 |
| **SNTG1** | 2.46484525982859E-09 | -0.106589973162229 | 0.183 | 0.61 | 0.000115091019717176 |
| **SERINC5** | 2.58131431047257E-09 | -0.143002231162553 | 0.213 | 0.634 | 0.000120529309098896 |
| **EGFEM1P** | 3.07849195015679E-09 | -0.298882359691475 | 0.19 | 0.61 | 0.000143744024628671 |
| **ITPK1** | 3.2419673654201E-09 | -0.161176386917858 | 0.369 | 0.854 | 0.000151377182193561 |
| **CNKSR3** | 3.26493274941645E-09 | -0.228785577790522 | 0.281 | 0.78 | 0.000152449504868502 |
| **GLUL** | 3.38280081468391E-09 | -0.408498048595728 | 0.468 | 0.951 | 0.000157953118440036 |
| **DBNDD1** | 3.77021742049483E-09 | -0.131893382517435 | 0.129 | 0.512 | 0.000176042762015165 |
| **HADH** | 3.97443625343418E-09 | 0.738283948384165 | 0.977 | 0.976 | 0.000185578351981602 |
| **IFIH1** | 3.98708128257194E-09 | -0.100640800633441 | 0.179 | 0.634 | 0.000186168786327132 |
| **IAPP** | 4.33398259592275E-09 | 1.37655206770199 | 0.981 | 0.976 | 0.000202366649351421 |
| **CYP3A5** | 4.37798087722224E-09 | -0.201286449313931 | 0.057 | 0.341 | 0.000204421061100138 |
| **RBM47** | 4.65953648219671E-09 | -0.295751309896554 | 0.46 | 0.829 | 0.000217567736963211 |
| **STARD3** | 4.89882945829082E-09 | -0.234644028524092 | 0.354 | 0.829 | 0.000228741043895973 |
| **PPP1R14C** | 5.65144799390751E-09 | -0.14783510199743 | 0.137 | 0.537 | 0.000263883061179523 |
| **PLP2** | 6.22501414015368E-09 | -0.365060443857344 | 0.829 | 1 | 0.000290664585246196 |
| **IRF6** | 6.96524644059392E-09 | -0.152353007755608 | 0.224 | 0.659 | 0.000325228252050652 |
| **GDF15** | 7.10620732190246E-09 | -0.199546772466943 | 0.16 | 0.585 | 0.000331810138481591 |
| **HIST2H2BE** | 7.3292121735427E-09 | -0.177944928785137 | 0.194 | 0.634 | 0.00034222290401923 |

| | | | | | |
|---|---|---|---|---|---|
| **SPATA2L** | 7.3396328719582E-09 | -0.124427666339213 | 0.278 | 0.78 | 0.000342709477690344 |
| **PCSK1** | 7.58655538239073E-09 | 0.948613105374904 | 0.966 | 0.976 | 0.000354239030469971 |
| **GALE** | 7.90176836193064E-09 | -0.177341911118739 | 0.16 | 0.561 | 0.000368957270123628 |
| **C2orf54** | 8.62950343758296E-09 | -0.112420632626283 | 0.038 | 0.293 | 0.000402937404011061 |
| **PAFAH1B1** | 8.67512586376656E-09 | -0.257650614077808 | 0.605 | 0.976 | 0.000405067651956852 |
| **KIAA1217** | 8.81189214091369E-09 | -0.103722405286552 | 0.183 | 0.61 | 0.000411453679735683 |
| **AC006369.2** | 1.0625592148995E-08 | -0.241333728768447 | 0.46 | 0.902 | 0.000496140774213024 |
| **SELENBP1** | 1.09234867884196E-08 | -0.125349510379927 | 0.042 | 0.293 | 0.000510050368611677 |
| **ACOX3** | 1.12169275490296E-08 | -0.124317426623185 | 0.122 | 0.488 | 0.00052375199804684 |
| **SAMD5** | 1.12784146411401E-08 | -0.119720806593334 | 0.194 | 0.61 | 0.000526623014838753 |
| **FAM84A** | 1.134272148547E-08 | -0.20982506900988 | 0.213 | 0.634 | 0.000529625694321052 |
| **NR4A2** | 1.20535994813806E-08 | -0.467014125474248 | 0.411 | 0.829 | 0.000562818720584104 |
| **CCDC86** | 1.2442246728223E-08 | -0.273231196024162 | 0.388 | 0.902 | 0.000580965826480915 |
| **STON2** | 1.26221456800816E-08 | -0.141721735760033 | 0.148 | 0.537 | 0.000589365848240048 |
| **HSPB1** | 1.35883200709795E-08 | -0.334836135423 | 0.578 | 0.976 | 0.000634479429074245 |
| **VSNL1** | 1.45067989019188E-08 | -0.109887401106146 | 0.053 | 0.317 | 0.000677365961127292 |
| **TPM4** | 1.50107224830011E-08 | -0.472373556104234 | 0.616 | 1 | 0.000700895664898769 |
| **PDZK1** | 1.71986070995038E-08 | -0.169746072917075 | 0.087 | 0.39 | 0.00080305456129713 |
| **HLA-E** | 1.92159889166398E-08 | -0.621977047581513 | 0.753 | 0.976 | 0.00089725217048466 |
| **B2M** | 2.02145676588688E-08 | -0.471747228155497 | 1 | 1 | 0.000943878807695563 |
| **GS1-358P8.4** | 2.06749497869758E-08 | -0.116802545607801 | 0.171 | 0.561 | 0.00096537543040326 |
| **ASS1** | 2.08672920914358E-08 | -0.246850787653879 | 0.278 | 0.732 | 0.00097435646962541 |
| **FCGRT** | 2.22324341662016E-08 | -0.163457789105446 | 0.403 | 0.927 | 0.00103809904852245 |
| **AMBP** | 2.56582194414504E-08 | -0.142331606192386 | 0.015 | 0.195 | 0.00119805924037964 |
| **HMGCS2** | 3.09480565976185E-08 | -0.198868908198224 | 0.03 | 0.244 | 0.0014450576067126 |
| **SQRDL** | 3.13480163834251E-08 | -0.155518065579625 | 0.179 | 0.585 | 0.00146373292899127 |
| **PYGB** | 3.17501965576258E-08 | -0.23584297846244 | 0.414 | 0.927 | 0.00148251192786522 |
| **RAP2B** | 3.18034056312493E-08 | -0.265062454496136 | 0.342 | 0.805 | 0.00148499641913992 |
| **PFKP** | 3.28225561414851E-08 | -0.273863760153452 | 0.399 | 0.854 | 0.00153258361391436 |
| **CNN3** | 3.82161211597898E-08 | -0.137675008372701 | 0.152 | 0.537 | 0.00178442534531406 |
| **MARVELD3** | 4.32734898398479E-08 | -0.122506337577707 | 0.137 | 0.488 | 0.00202056906109202 |
| **MAGIX** | 4.44998560899934E-08 | -0.128330412789065 | 0.217 | 0.634 | 0.00207783178041006 |
| **HLA-A** | 4.48051919660493E-08 | -0.511624802407558 | 0.966 | 1 | 0.00209208882847074 |
| **CD63** | 4.67964365026249E-08 | -0.355350185653301 | 1 | 1 | 0.00218506600961706 |
| **CD99L2** | 4.69904683333364E-08 | -0.247346532279699 | 0.445 | 0.927 | 0.00219412593788848 |
| **CNTN1** | 4.77963795217897E-08 | -0.333642539553273 | 0.555 | 0.951 | 0.00223175634901093 |
| **TNFRSF19** | 4.86297149475667E-08 | -0.145071160231571 | 0.148 | 0.537 | 0.00227066728004673 |

| | | | | | |
|---|---|---|---|---|---|
| **ST3GAL1** | 5.06814699560003E-08 | -0.104961005274164 | 0.205 | 0.634 | 0.00236646987665552 |
| **COX8A** | 5.55568823323892E-08 | -0.261515087297299 | 0.973 | 1 | 0.00259411750674625 |
| **OCRL** | 5.82058406287334E-08 | -0.193274216505601 | 0.255 | 0.659 | 0.00271780531647745 |
| **RAB27A** | 5.85672042548484E-08 | -0.12397781106214 | 0.24 | 0.659 | 0.00273467846827164 |
| **CCDC150** | 5.92413761605274E-08 | -0.13113190014392 | 0.087 | 0.39 | 0.00276615757706351 |
| **DIAPH2** | 6.33480436517296E-08 | -0.134490144726423 | 0.205 | 0.585 | 0.00295791020223021 |
| **C15orf48** | 6.68870032021381E-08 | -0.323973792863738 | 0.734 | 0.927 | 0.00312315484051744 |
| **CTD-2368P22.1** | 7.31782103216294E-08 | -0.10319131625586 | 0.228 | 0.61 | 0.00341691017454784 |
| **TMEM8A** | 7.45261682088694E-08 | -0.185040593009975 | 0.392 | 0.854 | 0.00347985037217674 |
| **ATP2B4** | 7.8336585228431E-08 | -0.14971561503855 | 0.163 | 0.537 | 0.00365777017407113 |
| **SESN3** | 8.32113037330128E-08 | -0.204781858152468 | 0.221 | 0.61 | 0.00388538540520557 |
| **NFKB1** | 9.21110844883714E-08 | -0.112230323730427 | 0.122 | 0.463 | 0.00430094286801552 |
| **NPTN** | 9.9281767336315E-08 | -0.147717230771846 | 0.749 | 1 | 0.00463576356223456 |
| **WEE1** | 1.00717314237757E-07 | -0.106658472903933 | 0.125 | 0.463 | 0.00470279355370361 |
| **SLC7A2** | 1.0531664998398E-07 | -0.399715204016039 | 0.673 | 1 | 0.00491755033770198 |
| **FSD1L** | 1.08368089572399E-07 | -0.139305244666998 | 0.198 | 0.585 | 0.00506003120640401 |
| **KCNJ6** | 1.12089549601283E-07 | -0.349160465439164 | 0.673 | 0.976 | 0.00523379733953271 |
| **AGT** | 1.18043289026403E-07 | -0.617739464473691 | 0.141 | 0.488 | 0.00551179529450985 |
| **ASXL3** | 1.23421541530956E-07 | -0.18875741676862 | 0.19 | 0.537 | 0.00576292203870494 |
| **MYO15B** | 1.25322443574871E-07 | -0.105277533697623 | 0.369 | 0.732 | 0.00585168085784144 |
| **LPP** | 1.26870108739966E-07 | -0.262446610665425 | 0.475 | 0.878 | 0.00592394598739523 |
| **FGF12** | 1.64340182885284E-07 | -0.179027428263209 | 0.376 | 0.902 | 0.00767353615946254 |
| **TACC2** | 1.68110017466696E-07 | -0.151750853153569 | 0.312 | 0.756 | 0.00784956104557242 |
| **LMNA** | 1.6902186430599E-07 | -0.393398571764191 | 0.798 | 1 | 0.00789213791003957 |
| **FAM46A** | 1.78516993230898E-07 | -0.431812412657705 | 0.692 | 1 | 0.00833549396493034 |
| **KIF9** | 1.87300000216573E-07 | -0.144861250794088 | 0.293 | 0.707 | 0.00874559891011246 |
| **SPINT2** | 1.95721056647586E-07 | -0.373186260645016 | 0.966 | 1 | 0.00913880329804573 |
| **ADCY1** | 1.98096863883858E-07 | -0.134092958005278 | 0.251 | 0.634 | 0.009249736865329 |
| **KIAA1671** | 2.1232156536929E-07 | -0.133497461217409 | 0.148 | 0.488 | 0.00991393085178824 |
| **ARNT2** | 2.39247753881151E-07 | -0.265731798237621 | 0.228 | 0.61 | 0.0111711953719726 |
| **NECAB2** | 2.48222004822834E-07 | -0.100254591926609 | 0.141 | 0.488 | 0.0115902300711926 |
| **SH3KBP1** | 2.58151990366675E-07 | -0.203259073307797 | 0.274 | 0.61 | 0.0120538908861911 |
| **TIMM21** | 2.65856706024765E-07 | -0.154707308889663 | 0.213 | 0.61 | 0.0124136471744144 |
| **RFX6** | 2.90100203626687E-07 | -0.247438083250708 | 0.681 | 0.976 | 0.0135456488079409 |
| **CALY** | 3.06168764354115E-07 | -0.189031586003503 | 0.281 | 0.659 | 0.0142959381139867 |
| **TMEM120A** | 3.06213843065082E-07 | -0.189467854373835 | 0.376 | 0.829 | 0.0142980429742379 |
| **BACE2** | 3.08409329971513E-07 | -0.162877009798988 | 0.19 | 0.585 | 0.0144005568443599 |

| | | | | | |
|---|---|---|---|---|---|
| **RSAD2** | 3.19749686137765E-07 | -0.415544271047567 | 0.129 | 0.439 | 0.0149300720948307 |
| **RASD1** | 3.28354847461563E-07 | 0.569230778890019 | 0.939 | 0.951 | 0.0153318728925227 |
| **RRP1** | 3.40617113896633E-07 | -0.12956793540251 | 0.281 | 0.707 | 0.0159044348991755 |
| **NBEAL2** | 3.49834897737663E-07 | -0.177814786361377 | 0.243 | 0.61 | 0.0163348408800647 |
| **LMF2** | 3.54685690360129E-07 | -0.12871121792804 | 0.43 | 0.854 | 0.0165613389399855 |
| **CFL1** | 3.71529751469948E-07 | -0.267479305670813 | 0.97 | 1 | 0.0173478386853863 |
| **REEP4** | 3.87767105768608E-07 | -0.181266217609924 | 0.346 | 0.805 | 0.0181060094696536 |
| **MLPH** | 3.94302639309347E-07 | -0.112141141864872 | 0.065 | 0.317 | 0.0184111731372713 |
| **SLC12A8** | 4.06078741639464E-07 | -0.182357812086416 | 0.224 | 0.61 | 0.0189610346833715 |
| **NEURL1** | 4.17294296785468E-07 | -0.293042424871972 | 0.643 | 1 | 0.0194847225998038 |
| **MCOLN3** | 4.33993595886029E-07 | -0.308246275632129 | 0.354 | 0.756 | 0.0202644629727063 |
| **CMPK2** | 4.476515898827E-07 | -0.12853030252445 | 0.137 | 0.488 | 0.0209021956863929 |
| **RP5-1158E12.3** | 4.59439170318744E-07 | -0.115514127039964 | 0.148 | 0.463 | 0.0214525931796931 |
| **TRNP1** | 4.80756563270901E-07 | -0.114612350923267 | 0.35 | 0.732 | 0.0224479662088082 |
| **MYH10** | 5.15120194666163E-07 | -0.298286083184947 | 0.525 | 0.878 | 0.0240525072495472 |
| **REEP3** | 5.38794178396491E-07 | -0.275040422805634 | 0.684 | 1 | 0.0251579165718673 |
| **CD44** | 5.67825194623887E-07 | -0.134755963460512 | 0.221 | 0.634 | 0.0265134618125732 |
| **TSPAN33** | 5.68753382674057E-07 | -0.12083722102222 | 0.19 | 0.585 | 0.0265568016971997 |
| **TXNDC11** | 5.8881972803306E-07 | -0.321180582103556 | 0.437 | 0.902 | 0.0274937595610477 |
| **HTR1F** | 5.93025502056321E-07 | -0.132448597963932 | 0.103 | 0.39 | 0.0276901397675158 |
| **ARSE** | 6.45057519799064E-07 | -0.105821444758515 | 0.095 | 0.39 | 0.0301196707719777 |
| **CNNM1** | 6.69174652750394E-07 | -0.175290412440158 | 0.38 | 0.829 | 0.0312457720608741 |
| **DECR2** | 6.82244780152228E-07 | -0.131325996146954 | 0.259 | 0.683 | 0.031856055519648 |
| **HN1L** | 6.85203302319844E-07 | -0.160041305710893 | 0.293 | 0.683 | 0.0319941977952205 |
| **TMEM136** | 7.34404212497882E-07 | -0.10266490735961 | 0.114 | 0.415 | 0.0342915358941636 |
| **VSTM2A** | 7.34626291046718E-07 | -0.194686952442804 | 0.023 | 0.195 | 0.0343019054078444 |
| **GAPDH** | 7.37770131097038E-07 | -0.36367119715831 | 1 | 1 | 0.034448700731314 |
| **TAP1** | 7.41155356831926E-07 | -0.127693855783545 | 0.163 | 0.512 | 0.0346067670765531 |
| **PLS3** | 7.44751996322794E-07 | -0.103219090756 | 0.08 | 0.341 | 0.0347747049643002 |
| **PFN1** | 7.59736799208726E-07 | -0.287892711206075 | 0.977 | 1 | 0.035474390365453 |
| **SLC25A33** | 8.26661639829697E-07 | -0.223162575580097 | 0.513 | 0.927 | 0.038599311948568 |
| **CADM1** | 8.70651317093809E-07 | 0.551192355837539 | 0.954 | 1 | 0.0406533219490612 |
| **C16orf45** | 8.74456752822893E-07 | -0.248390596508034 | 0.513 | 0.902 | 0.0408310091595594 |
| **AMDHD2** | 8.94244466935147E-07 | -0.135178139913216 | 0.327 | 0.805 | 0.0417549568946028 |
| **ODF2L** | 8.9959975679822E-07 | -0.143261255127791 | 0.342 | 0.756 | 0.0420050114441793 |
| **PCDH17** | 9.15423929557437E-07 | -0.261227294827343 | 0.555 | 0.878 | 0.0427438895428254 |
| **CHKA** | 9.26403394114631E-07 | -0.147682467636444 | 0.323 | 0.78 | 0.0432565536813945 |

| | | | | | |
|---|---|---|---|---|---|
| **SAMD11** | 9.3624052421446E-07 | 0.385961705355652 | 0.958 | 0.927 | 0.0437158787971458 |
| **GCH1** | 9.67716540028913E-07 | -0.479161571814384 | 0.688 | 0.976 | 0.0451855884035701 |
| **NMNAT3** | 9.68214521970206E-07 | -0.110577801261496 | 0.19 | 0.561 | 0.0452088406743548 |
| **PAPSS2** | 9.8356617344066E-07 | -0.468488906378998 | 0.764 | 0.927 | 0.0459256553364647 |
| **HN1** | 9.94086754639166E-07 | -0.140384005426195 | 0.304 | 0.683 | 0.0464168928343666 |
| **LINC00888** | 1.05787298657453E-06 | -0.147545785567581 | 0.24 | 0.634 | 0.0493952633621243 |

Supplemental Table 3 - LDHA_pos_vs_neg β-cells in non-diabetics

| | p_val | avg_logFC | pct.1 | pct.2 | p_val_adj |
|---|---|---|---|---|---|
| **LDHA** | 1.64114750334685E-66 | 0.951996903185496 | 1 | 0 | 7.66301003737743E-62 |
| **GC** | 3.66840982684287E-22 | 1.33405277882449 | 0.642 | 0.105 | 1.71289060044774E-17 |
| **TM4SF4** | 7.99423058724265E-22 | 1.73220356479636 | 0.687 | 0.152 | 3.73274608810121E-17 |
| **MUC13** | 8.86763613418312E-22 | 0.459149792986577 | 0.463 | 0.03 | 4.14056534013412E-17 |
| **LOXL4** | 5.54217541380335E-21 | 1.75516013006808 | 0.627 | 0.122 | 2.5878079659672E-16 |
| **GPR119** | 6.60561821435195E-21 | 0.215699676338634 | 0.507 | 0.046 | 3.08436131282735E-16 |
| **ARX** | 1.8581619153234E-19 | 0.170481378480828 | 0.507 | 0.059 | 8.67631543121956E-15 |
| **TMEM236** | 6.86173139992789E-19 | 0.305113525090999 | 0.493 | 0.059 | 3.20394824256833E-14 |
| **DMD** | 8.29635474194723E-19 | 0.115513893585533 | 0.478 | 0.051 | 3.87381691965742E-14 |
| **DPP4** | 1.06882460251026E-18 | 0.352358690635707 | 0.478 | 0.051 | 4.99066271650117E-14 |
| **POPDC3** | 2.38412789861245E-18 | 0.158358610890089 | 0.358 | 0.013 | 1.11322083969911E-13 |
| **VIM** | 8.31220457108224E-18 | 1.00826504307307 | 0.642 | 0.173 | 3.88121768037543E-13 |
| **F10** | 1.41402496093471E-17 | 0.526170197889639 | 0.597 | 0.127 | 6.60250675009246E-13 |
| **FXYD3** | 3.18091815472483E-17 | 0.246523656800718 | 0.552 | 0.097 | 1.48526611398566E-12 |
| **PAPPA2** | 4.05584339698155E-17 | 0.313437138234199 | 0.433 | 0.046 | 1.89379495735259E-12 |
| **TMEM176A** | 1.24216107981542E-16 | 0.800326719280848 | 0.701 | 0.27 | 5.80002272998212E-12 |
| **MAOB** | 1.33852760826462E-16 | 0.219019488273688 | 0.418 | 0.042 | 6.24998696127001E-12 |
| **SERPINA1** | 2.62693669830977E-16 | 1.11674228185663 | 0.493 | 0.076 | 1.22659555254178E-11 |
| **PTPRT** | 1.38664718648904E-15 | 0.187692470520594 | 0.388 | 0.038 | 6.47467170787328E-11 |
| **CD36** | 2.79576042360803E-15 | 0.503209918523335 | 0.403 | 0.051 | 1.3054244145953E-10 |
| **MBOAT4** | 3.22882409280515E-15 | 0.221909532541586 | 0.433 | 0.059 | 1.50763483365351E-10 |
| **MRC1** | 4.13642463344731E-15 | 0.536369177713974 | 0.478 | 0.08 | 1.93142075409555E-10 |
| **APOH** | 8.75684745538921E-15 | 0.366128541262288 | 0.403 | 0.051 | 4.08883478234489E-10 |
| **VIL1** | 1.01666489143129E-14 | 0.341890697389346 | 0.403 | 0.051 | 4.74711337756011E-10 |
| **S100A10** | 1.09750435677189E-14 | 0.30498962453292 | 0.567 | 0.131 | 5.12457709307499E-10 |
| **ADGRG5** | 1.12667782361848E-14 | 0.130948281708067 | 0.373 | 0.038 | 5.26079676182178E-10 |
| **KCTD12** | 2.52538484150071E-14 | 0.312679022480841 | 0.597 | 0.165 | 1.17917794404192E-09 |
| **SYNDIG1** | 2.76212417138286E-14 | 0.206114023843853 | 0.373 | 0.038 | 1.2897186393438E-09 |
| **TMEM86B** | 4.45257683638303E-14 | 0.14758773261736 | 0.478 | 0.089 | 2.07904170221233E-09 |
| **TMEM176B** | 5.41590958116192E-14 | 0.934765569589977 | 0.731 | 0.35 | 2.52885066073194E-09 |
| **SERPINA10** | 5.77279248794898E-14 | 0.123720163292361 | 0.284 | 0.013 | 2.69548999639802E-09 |
| **GPC6** | 7.78677359133967E-14 | 0.214798390146107 | 0.418 | 0.063 | 3.63587819300423E-09 |
| **PRSS3** | 1.12979328621072E-13 | 0.166211725142355 | 0.328 | 0.03 | 5.27534379130372E-09 |
| **MYRF** | 1.65357279825866E-13 | 0.136491028398933 | 0.418 | 0.068 | 7.72102746690918E-09 |
| **SH3RF1** | 1.84253046542458E-13 | 0.130927254425657 | 0.463 | 0.084 | 8.60332750220698E-09 |

| | | | | | |
|---|---|---|---|---|---|
| **CRH** | 2.04557901779133E-13 | 0.547004919130399 | 0.358 | 0.042 | 9.55142210777304E-09 |
| **PTGER4** | 2.12926067440617E-13 | 0.174211293547291 | 0.269 | 0.013 | 9.94215686700471E-09 |
| **FABP5** | 2.85945514354763E-13 | 0.220201119028045 | 0.552 | 0.148 | 1.33516539017669E-08 |
| **FAP** | 1.04167666172828E-12 | 0.349682707349114 | 0.463 | 0.11 | 4.86390083660788E-08 |
| **GCG** | 1.04720709537804E-12 | 2.58363091929506 | 0.896 | 0.743 | 4.8897240904487E-08 |
| **TTR** | 1.42176238039753E-12 | 1.61092386529203 | 1 | 0.962 | 6.63863508279021E-08 |
| **DOK4** | 1.54141123508717E-12 | 0.251264019944414 | 0.507 | 0.114 | 7.19731147999252E-08 |
| **TTLL12** | 2.11733191130405E-12 | 0.186668909495672 | 0.582 | 0.169 | 9.88645789345202E-08 |
| **ELF3** | 3.91457797745302E-12 | 0.347054670650329 | 0.731 | 0.287 | 1.82783389501214E-07 |
| **QDPR** | 6.56187451204669E-12 | 0.584728843512733 | 0.97 | 0.743 | 3.06393606590996E-07 |
| **CSGALNACT1** | 9.90079163005366E-12 | 0.219412610721899 | 0.373 | 0.059 | 4.62297663582096E-07 |
| **NPNT** | 9.92288555766884E-12 | 0.226916080330443 | 0.358 | 0.055 | 4.63329295344231E-07 |
| **INHBA** | 1.22236507081108E-11 | 0.145653968276781 | 0.567 | 0.156 | 5.70758922513817E-07 |
| **FXYD5** | 1.550044084009E-11 | 0.363984674230653 | 0.537 | 0.16 | 7.23762084146323E-07 |
| **CLU** | 1.79832050283602E-11 | 1.08826863034836 | 0.955 | 0.928 | 8.39689792389224E-07 |
| **KIF12** | 2.77995103714043E-11 | 0.116841587950741 | 0.537 | 0.152 | 1.29804253777198E-06 |
| **INS** | 2.79815513796716E-11 | -0.499926663126052 | 1 | 1 | 1.30654257857101E-06 |
| **SMIM24** | 3.23053938236567E-11 | 0.419683992087003 | 0.582 | 0.19 | 1.508435753808E-06 |
| **FEV** | 3.58958493376168E-11 | 0.30217065200645 | 0.552 | 0.186 | 1.67608489312134E-06 |
| **PLS1** | 5.75334087128064E-11 | 0.30376454947276 | 0.896 | 0.456 | 2.68640745302707E-06 |
| **TMEM45B** | 6.19520021837491E-11 | 0.128489357599557 | 0.254 | 0.021 | 2.8927248379658E-06 |
| **RHOC** | 6.3280787685727E-11 | 0.161919850265011 | 0.418 | 0.089 | 2.95476981940965E-06 |
| **SPOCK3** | 6.63975103339341E-11 | 0.208603286258222 | 0.373 | 0.072 | 3.10029895002239E-06 |
| **USH1C** | 7.15516825999112E-11 | 0.342250810219937 | 0.552 | 0.181 | 3.34096271563765E-06 |
| **MYO10** | 7.50421294018839E-11 | 0.341100547988331 | 0.403 | 0.084 | 3.50394214816216E-06 |
| **ADGRG2** | 1.09662394137265E-10 | 0.314507266786 | 0.448 | 0.122 | 5.12046616945131E-06 |
| **GPER1** | 1.1742724287507E-10 | 0.103386785774238 | 0.299 | 0.038 | 5.48303025156566E-06 |
| **NEDD9** | 1.33487322667676E-10 | 0.230683740615813 | 0.478 | 0.127 | 6.23292355732178E-06 |
| **REG4** | 1.46721655948658E-10 | 0.384047426352696 | 0.284 | 0.034 | 6.85087428121068E-06 |
| **IGFBP2** | 1.8191590185967E-10 | 0.485376709823095 | 0.687 | 0.308 | 8.49419920553355E-06 |
| **ERBB3** | 2.03627281389866E-10 | 0.164988928186603 | 0.343 | 0.059 | 9.50796864993701E-06 |
| **ZFP36** | 2.67513785971716E-10 | 0.204941569785833 | 0.597 | 0.203 | 1.24910212083773E-05 |
| **SPTSSB** | 3.17201836011391E-10 | 0.391648923772127 | 0.493 | 0.139 | 1.48111053288799E-05 |
| **RGS4** | 3.55487279757076E-10 | 0.794384331352851 | 0.701 | 0.384 | 1.65987675536971E-05 |
| **CNIH3** | 5.02955350334959E-10 | 0.133321528160397 | 0.418 | 0.097 | 2.34844941731903E-05 |
| **LMCD1** | 8.06672643291743E-10 | 0.240885308302282 | 0.418 | 0.101 | 3.76659657332213E-05 |
| **WDR20** | 1.31008635686438E-09 | 0.204072776133041 | 0.657 | 0.245 | 6.11718622610686E-05 |

| | | | | | |
|---|---|---|---|---|---|
| **LDB2** | 1.79816121018092E-09 | 0.114565993580094 | 0.239 | 0.025 | 8.39615413869777E-05 |
| **KCNIP1** | 2.19827183161351E-09 | 0.153968182030495 | 0.284 | 0.042 | 0.00010264390663353 |
| **KLHL41** | 2.74159328190477E-09 | 0.178075782307023 | 0.358 | 0.08 | 0.000128013215111979 |
| **PLIN3** | 2.85053166677621E-09 | 0.12136044751498 | 0.403 | 0.097 | 0.000133099875116782 |
| **HMGB3** | 3.44570649112334E-09 | 0.302334788102151 | 0.642 | 0.308 | 0.000160890373190022 |
| **KCNA5** | 3.72446050684695E-09 | 0.193058197282608 | 0.507 | 0.152 | 0.000173906234446204 |
| **IRX2** | 3.87780658699967E-09 | 0.133233170265599 | 0.418 | 0.118 | 0.000181066422966775 |
| **CDO1** | 4.14877996282593E-09 | 0.114813871339144 | 0.194 | 0.013 | 0.000193718982804231 |
| **PDK4** | 4.59861821359507E-09 | 0.855075799601585 | 0.776 | 0.485 | 0.000214723280247395 |
| **HIGD1A** | 5.07610772340449E-09 | 0.445306640002348 | 0.97 | 0.73 | 0.000237018697928926 |
| **FAM110B** | 5.30520314492346E-09 | 0.172184439118193 | 0.493 | 0.165 | 0.000247715850445911 |
| **IMPA2** | 6.79564108038939E-09 | 0.104722262270594 | 0.328 | 0.063 | 0.000317308868966622 |
| **ANXA4** | 8.00690471541952E-09 | 0.228834972117107 | 0.567 | 0.203 | 0.000373866401877083 |
| **SLC6A19** | 8.14532098484088E-09 | 0.153841373579795 | 0.254 | 0.034 | 0.000380329472745175 |
| **CAMK2G** | 8.43854894273494E-09 | 0.381228715053018 | 0.672 | 0.346 | 0.000394021165783122 |
| **SLC38A4** | 8.47799228818735E-09 | 0.203207015824901 | 0.567 | 0.215 | 0.000395862893912332 |
| **SPATS2L** | 9.87629472259489E-09 | 0.180466801485688 | 0.537 | 0.198 | 0.000461153829482123 |
| **TPD52L1** | 1.23600506594519E-08 | 0.167485865076117 | 0.313 | 0.063 | 0.000577127845441788 |
| **GALNT13** | 1.42364425264676E-08 | 0.128693841426487 | 0.343 | 0.08 | 0.000664742210888353 |
| **AOX1** | 1.60734337066805E-08 | 0.101709904741844 | 0.269 | 0.042 | 0.000750516840066034 |
| **GLS** | 1.65933071742719E-08 | 0.365805572180302 | 0.821 | 0.506 | 0.000774791291888277 |
| **TAGLN2** | 1.99383825816161E-08 | 0.418628481413672 | 0.985 | 0.869 | 0.000930982897883401 |
| **ITPR1** | 2.19613538955571E-08 | 0.116183387478869 | 0.448 | 0.135 | 0.00102544149744525 |
| **ASS1** | 2.76473434314326E-08 | 0.23229640278424 | 0.627 | 0.257 | 0.00129093740684388 |
| **BID** | 2.78660285960687E-08 | 0.191238648380105 | 0.537 | 0.211 | 0.00130114847323624 |
| **UCP2** | 2.84147792665089E-08 | 0.301577473159693 | 0.463 | 0.152 | 0.0013267712882911 |
| **COTL1** | 2.93216003915544E-08 | 0.268080387119718 | 0.731 | 0.35 | 0.00136911348708285 |
| **CACNB2** | 3.13635790518535E-08 | 0.314983764707493 | 0.806 | 0.498 | 0.0014644595966682 |
| **GRIA3** | 3.70888175197838E-08 | 0.353194899191544 | 0.716 | 0.418 | 0.00173178815645127 |
| **PLPPR1** | 3.80673944226441E-08 | 0.14751715380499 | 0.254 | 0.042 | 0.00177748084777652 |
| **ARFGEF3** | 4.97357052760959E-08 | 0.494501135306167 | 0.925 | 0.835 | 0.00232230928645675 |
| **HPS1** | 5.76100940466723E-08 | 0.363536717970002 | 0.851 | 0.561 | 0.00268998812132127 |
| **BACE2** | 5.89705314488894E-08 | 0.186964562825316 | 0.507 | 0.169 | 0.00275351102494299 |
| **CYSTM1** | 6.67224027712105E-08 | 0.423989748271546 | 1 | 0.92 | 0.00311546915259613 |
| **CRYBA2** | 7.99868229225285E-08 | 1.13707470426333 | 0.896 | 0.713 | 0.00373482472272162 |
| **SPINT2** | 8.70021758417989E-08 | 0.343273494899119 | 0.985 | 0.966 | 0.00406239259658111 |
| **CD82** | 1.08735839650519E-07 | 0.287063624429949 | 0.597 | 0.291 | 0.0050772025608017 |

| | | | | | |
|---|---|---|---|---|---|
| **CNN3** | 1.13301028459825E-07 | 0.153052522022853 | 0.433 | 0.139 | 0.00529036492187462 |
| **HS3ST3B1** | 1.29916884868576E-07 | 0.107214600155916 | 0.269 | 0.051 | 0.00606620910516841 |
| **GLUL** | 1.44401639332374E-07 | 0.33468890727444 | 0.776 | 0.464 | 0.00674254574534652 |
| **RBM47** | 1.46353810003605E-07 | 0.182862526035769 | 0.746 | 0.443 | 0.00683369845049833 |
| **VSTM2A** | 1.6426573973484E-07 | 0.12143740690612 | 0.164 | 0.013 | 0.0076700601854389 |
| **ELL2** | 1.6561267709899E-07 | 0.2880267487257 | 0.925 | 0.747 | 0.00773295273178316 |
| **GYG2** | 1.89273009113081E-07 | 0.13054704685759 | 0.313 | 0.076 | 0.0088377246145171 |
| **MFSD3** | 3.14322656128429E-07 | 0.140794256413199 | 0.657 | 0.321 | 0.0146766677826047 |
| **SLC40A1** | 3.15476195886564E-07 | 0.356932338347556 | 0.582 | 0.266 | 0.0147305300145313 |
| **CPNE8** | 3.19655970925759E-07 | 0.164067805750085 | 0.582 | 0.236 | 0.0149256962504365 |
| **ATG12** | 3.2145587005042E-07 | 0.204126100304274 | 0.896 | 0.586 | 0.0150097389402643 |
| **NECAB2** | 3.40063513297578E-07 | 0.101256299151911 | 0.403 | 0.127 | 0.0158785856264038 |
| **DUSP3** | 3.40422700667742E-07 | 0.294019906884102 | 0.97 | 0.709 | 0.0158953571622789 |
| **PLCE1** | 3.79375401676202E-07 | 0.461997362330395 | 0.493 | 0.194 | 0.0177141756304669 |
| **ZSWIM7** | 4.30024990041344E-07 | 0.219276977948921 | 0.806 | 0.477 | 0.0200791568600005 |
| **EZR** | 4.44414528188467E-07 | 0.553985206470077 | 0.866 | 0.738 | 0.0207510475647041 |
| **PRELID1** | 4.824661493907E-07 | 0.239199376056823 | 0.955 | 0.776 | 0.0225277919135 |
| **TM4SF1** | 4.88799268124329E-07 | 0.323453553566645 | 0.239 | 0.046 | 0.0228235042265293 |
| **GPX3** | 5.07039338969541E-07 | 0.50103404597535 | 0.925 | 0.793 | 0.0236751878545048 |
| **CLIC1** | 5.09513791441142E-07 | 0.293201401440428 | 1 | 0.865 | 0.0237907274637613 |
| **ARRDC4** | 5.99498208455052E-07 | 0.165600788826052 | 0.403 | 0.135 | 0.0279923698473918 |
| **STARD3** | 6.20844793382753E-07 | 0.190245988670871 | 0.672 | 0.346 | 0.0289891059374209 |
| **VGF** | 6.23639061795961E-07 | 0.200092695589264 | 0.896 | 0.764 | 0.0291195787124388 |
| **CEACAM1** | 6.39218751027551E-07 | 0.177324729543472 | 0.388 | 0.127 | 0.0298470411417294 |
| **HLA-E** | 6.43740206284523E-07 | 0.395775730059028 | 0.94 | 0.738 | 0.0300581614520433 |
| **SMOC1** | 8.56256438332743E-07 | 0.141302783658491 | 0.418 | 0.148 | 0.0399811818750708 |
| **KCNH6** | 8.7121236462323E-07 | 0.137630823882766 | 0.463 | 0.177 | 0.0406795189413525 |
| **IDH2** | 8.76717866899378E-07 | 0.137981491777043 | 0.522 | 0.211 | 0.0409365873591327 |
| **RAP1GAP** | 9.00501402794399E-07 | 0.242670882524142 | 0.97 | 0.7 | 0.0420471120006789 |
| **LAMB1** | 9.12942430409065E-07 | 0.219609280075033 | 0.448 | 0.194 | 0.0426280209030905 |
| **KRTCAP3** | 9.68711724782071E-07 | 0.101970247040039 | 0.403 | 0.135 | 0.0452320565652492 |
| **PFKP** | 1.03764768162228E-06 | 0.24244565527215 | 0.701 | 0.392 | 0.0484508831979893 |
| **DAPK2** | 1.0469502654069E-06 | 0.162005745588011 | 0.642 | 0.312 | 0.0488852487426446 |
| **ENTPD6** | 1.0697214181684E-06 | 0.220121084393508 | 0.791 | 0.46 | 0.0499485021785369 |

Supplemental Table 4 - Cluster1_vs_cluster7 in T2D

| | p_val | avg_logFC | pct.1 | pct.2 | p_val_adj |
|---|---|---|---|---|---|
| **GC** | 4.61897173682652E-18 | -1.75384620522329 | 0.053 | 1 | 2.15673647307641E-13 |
| **IRX2** | 1.04019981188085E-15 | -0.407046154225671 | 0.08 | 0.941 | 4.85700498161523E-11 |
| **TM4SF4** | 2.4490644428097E-15 | -2.41122606397764 | 0.12 | 1 | 1.14354166028113E-10 |
| **LOXL4** | 5.85567109195713E-15 | -1.47350348354574 | 0.107 | 1 | 2.73418850296754E-10 |
| **CD36** | 1.61669703947165E-14 | -0.428518777930519 | 0.013 | 0.765 | 7.54884348640496E-10 |
| **MYO10** | 3.46966790131717E-14 | -0.58179604732486 | 0.053 | 0.882 | 1.62009203316202E-09 |
| **FXYD5** | 3.5032715414958E-14 | -0.605071634179043 | 0.08 | 0.941 | 1.63578258087063E-09 |
| **SMIM24** | 1.04849799157386E-13 | -0.986473776232365 | 0.107 | 1 | 4.89575167205585E-09 |
| **GPR119** | 1.45719705692095E-13 | -0.326609147276319 | 0.013 | 0.706 | 6.80409021788097E-09 |
| **POPDC3** | 1.90713604662193E-13 | -0.332002101538673 | 0 | 0.647 | 8.90499034249176E-09 |
| **FEV** | 2.45562830754685E-13 | -0.603707479397353 | 0.08 | 0.882 | 1.14660652564285E-08 |
| **ARX** | 2.66180830401534E-13 | -0.342669214243807 | 0.04 | 0.765 | 1.24287815139388E-08 |
| **MRC1** | 2.89590901314018E-13 | -0.546764556666198 | 0.053 | 0.824 | 1.35218679550554E-08 |
| **SEMA3E** | 2.9994758842511E-12 | -0.255878911343332 | 0.027 | 0.706 | 1.40054527463337E-07 |
| **SPOCK3** | 3.36271460214726E-12 | -0.515184015760364 | 0.013 | 0.647 | 1.57015232918062E-07 |
| **RGS4** | 3.68178660214654E-12 | -1.61630880085541 | 0.227 | 1 | 1.71913661814028E-07 |
| **FXYD3** | 4.97809664690215E-12 | -0.467899353118129 | 0.107 | 0.882 | 2.32442266733802E-07 |
| **USH1C** | 6.94771287418315E-12 | -0.414939685369255 | 0.093 | 0.882 | 3.24409557234234E-07 |
| **PTPRT** | 9.58902568717226E-12 | -0.213990767572244 | 0.067 | 0.765 | 4.47740376411134E-07 |
| **F10** | 1.71518555148785E-11 | -0.462729438082359 | 0.12 | 0.941 | 8.0087158955622E-07 |
| **S100A10** | 1.78604371203009E-11 | -0.679556991492739 | 0.12 | 0.882 | 8.33957390458209E-07 |
| **TMEM86B** | 2.10272476071303E-11 | -0.172691198682541 | 0.08 | 0.824 | 9.81825272519737E-07 |
| **TMEM236** | 3.29508198871083E-11 | -0.464063424961377 | 0.053 | 0.706 | 1.53857263298875E-06 |
| **VIM** | 3.51338875898938E-11 | -0.8728684270882 | 0.307 | 1 | 1.64050661323491E-06 |
| **PAPPA2** | 4.79496825745002E-11 | -0.505786537291023 | 0.027 | 0.647 | 2.23891452845114E-06 |
| **GPER1** | 4.80996004752198E-11 | -0.194577759808868 | 0 | 0.529 | 2.24591464498944E-06 |
| **KCTD12** | 5.34713347806787E-11 | -0.607379523437847 | 0.147 | 0.882 | 2.49673703491423E-06 |
| **TMEM176A** | 6.00631677206238E-11 | -0.851593912092431 | 0.28 | 1 | 2.80452949037909E-06 |
| **TMEM176B** | 9.10964554803416E-11 | -1.22678008347128 | 0.413 | 1 | 4.25356679574359E-06 |
| **ADGRG2** | 9.45331157528546E-11 | -0.640405023805937 | 0.107 | 0.824 | 4.41403477384804E-06 |
| **MAOB** | 1.02963841572922E-10 | -0.440518294040218 | 0.013 | 0.588 | 4.80769065456444E-06 |
| **PPP2R2B** | 1.26097810039361E-10 | -0.357902948122294 | 0.093 | 0.824 | 5.88788504416789E-06 |
| **TTR** | 1.91299526479748E-10 | -2.62357142834504 | 0.947 | 1 | 8.93234878991887E-06 |
| **GCG** | 2.0868521375957E-10 | -4.99027907597035 | 0.84 | 1 | 9.74413868607558E-06 |

| | | | | | |
|---|---|---|---|---|---|
| **A1CF** | 3.70061226642336E-10 | -0.411455012453429 | 0.133 | 0.882 | 1.72792688556106E-05 |
| **IGFBP2** | 4.55084351169166E-10 | -1.19529696156447 | 0.293 | 0.941 | 2.12492536091419E-05 |
| **C3orf80** | 7.05752926027205E-10 | -0.149216698111779 | 0 | 0.471 | 3.29537213749883E-05 |
| **HNF1A-AS1** | 9.02380885435607E-10 | -0.176184190991135 | 0.107 | 0.824 | 4.21348706836448E-05 |
| **SLC7A14** | 1.03649679139766E-09 | -0.508645403231932 | 0.133 | 0.824 | 4.83971446807311E-05 |
| **REG4** | 1.29355670219304E-09 | -1.28371025650491 | 0.053 | 0.647 | 6.04000430954995E-05 |
| **ADGRG5** | 1.61601372878157E-09 | -0.206090071298897 | 0.013 | 0.529 | 7.54565290379979E-05 |
| **SERPINA1** | 2.16955543164779E-09 | -1.4341418778488 | 0.12 | 0.765 | 0.00010130305176993 |
| **RNF144A** | 5.02414425942228E-09 | -0.12677854307854 | 0.04 | 0.588 | 0.000234592367905205 |
| **SLC6A19** | 5.02414425942228E-09 | -0.212716168473076 | 0.04 | 0.588 | 0.000234592367905205 |
| **ANK2** | 5.77337326319652E-09 | -0.292276256298082 | 0.12 | 0.765 | 0.000269576117778435 |
| **FABP5** | 6.23032122324754E-09 | -0.293351630767017 | 0.227 | 0.882 | 0.000290912388877097 |
| **WIPF3** | 7.18159201503102E-09 | -0.303334176214561 | 0.053 | 0.588 | 0.000335330075957844 |
| **DMD** | 9.36001934916503E-09 | -0.116912219766285 | 0.133 | 0.765 | 0.000437047383470563 |
| **FAP** | 9.5998182410938E-09 | -0.510496905358243 | 0.173 | 0.765 | 0.000448244313131393 |
| **PRLR** | 1.35490571076837E-08 | -0.17635825891236 | 0.013 | 0.471 | 0.000632646123529077 |
| **KLB** | 1.51835109483996E-08 | -0.174479140784221 | 0.013 | 0.471 | 0.000708963676713623 |
| **LDHA** | 1.59830865419349E-08 | -0.646819836002201 | 0.187 | 0.824 | 0.000746298259902567 |
| **GLS** | 1.99384819863579E-08 | -0.813506937008475 | 0.627 | 1 | 0.00093098753938901 |
| **GPC6** | 2.08610205659878E-08 | -0.230434900477263 | 0.067 | 0.647 | 0.000974063633287667 |
| **FAM110B** | 2.35838151505979E-08 | -0.158589389636725 | 0.107 | 0.706 | 0.00110119908082687 |
| **CD82** | 2.58681562852818E-08 | -0.381462538234588 | 0.187 | 0.882 | 0.00120786182142866 |
| **KIF12** | 2.71302526412591E-08 | -0.112818271369004 | 0.067 | 0.647 | 0.00126679288657831 |
| **CEACAM1** | 3.8655088498776E-08 | -0.266797782352356 | 0.053 | 0.588 | 0.00180492204727335 |
| **CRYBA2** | 6.09629617253389E-08 | -1.39684856878365 | 0.747 | 1 | 0.00284654357184125 |
| **SLC7A2** | 7.21989315779411E-08 | -0.842574042260619 | 0.6 | 1 | 0.0033711847121688 |
| **CYSTM1** | 7.77824334001041E-08 | -0.653179716145187 | 0.947 | 1 | 0.00363189516275106 |
| **MBOAT4** | 1.17774411921502E-07 | -0.240743699832836 | 0.027 | 0.471 | 0.00549924061585068 |
| **SYNDIG1** | 1.22323361846021E-07 | -0.253417572702418 | 0.013 | 0.412 | 0.00571164473467624 |
| **STK32B** | 1.31489715200691E-07 | -0.129683227822749 | 0 | 0.353 | 0.00613964927186588 |
| **VIL1** | 1.54767395080916E-07 | -0.414994063954574 | 0.08 | 0.588 | 0.00722655397851319 |
| **LRRFIP1** | 1.6497507895564E-07 | -0.482989975644888 | 0.4 | 1 | 0.00770318136167572 |
| **SPTSSB** | 1.98163199785136E-07 | -0.612464245775472 | 0.2 | 0.824 | 0.00925283428756737 |
| **APOH** | 2.19507212671022E-07 | -0.408682358856559 | 0.053 | 0.529 | 0.010249450281248 |
| **INS** | 3.21419889710142E-07 | 0.702139482673282 | 1 | 1 | 0.0150080589102357 |
| **EGFL7** | 3.49674103062842E-07 | -0.283620842332343 | 0.08 | 0.588 | 0.0163273328943133 |

| | | | | | |
|---|---|---|---|---|---|
| **PHLDB2** | 3.52376115148942E-07 | -0.373730419404478 | 0.267 | 0.882 | 0.0164534979446496 |
| **ITGB8** | 3.75499008303765E-07 | -0.194825964819791 | 0.147 | 0.765 | 0.0175331751947277 |
| **TMEM178B** | 3.98690515977937E-07 | -0.236966749370969 | 0.267 | 0.941 | 0.0186160562625578 |
| **PRKCE** | 5.22706877304955E-07 | -0.168853295439471 | 0.24 | 0.882 | 0.0244067522220003 |
| **AOX1** | 6.33242600360397E-07 | -0.268231408288758 | 0.04 | 0.471 | 0.029567996738628 |
| **COTL1** | 7.45774435252715E-07 | -0.631687390685186 | 0.387 | 0.941 | 0.034822445705255 |
| **ABHD15** | 8.03656680623846E-07 | -0.230532914842571 | 0.107 | 0.647 | 0.0375251413883692 |
| **CLU** | 8.24211806056677E-07 | -1.08807902692472 | 0.947 | 1 | 0.0384849218602044 |
| **ARFGEF3** | 9.12558392958818E-07 | -0.701475203694083 | 0.933 | 1 | 0.0426100890424261 |
| **POU6F2** | 9.91706484226383E-07 | -0.143894714081399 | 0.067 | 0.529 | 0.0463057508679825 |

Supplemental Table 5 - LDHA_pos_vs_neg β-cells in T2D

| | p_val | avg_logFC | pct.1 | pct.2 | p_val_adj |
|---|---|---|---|---|---|
| **LDHA** | 1.06337848359989E-20 | 0.736410359051582 | 1 | 0 | 4.96523315347295E-16 |
| **S100A10** | 2.10940452800272E-09 | 0.530047408028363 | 0.679 | 0.078 | 9.8494425626031E-05 |
| **SMIM24** | 3.02392597221245E-08 | 0.595945390537383 | 0.679 | 0.094 | 0.00141196175420516 |
| **TM4SF4** | 3.32354002996006E-08 | 1.41226615308684 | 0.679 | 0.109 | 0.00155186054618925 |
| **F10** | 1.94744497323796E-07 | 0.395163750085695 | 0.643 | 0.109 | 0.00909320481354001 |
| **GC** | 1.99127816305625E-07 | 1.10304171584566 | 0.571 | 0.078 | 0.00929787512675855 |
| **MYO10** | 2.26920678648673E-07 | 0.39430141233161 | 0.536 | 0.062 | 0.0105956072481425 |
| **USH1C** | 5.17680944886665E-07 | 0.324829649353735 | 0.571 | 0.094 | 0.0241720763595931 |
| **FXYD5** | 6.94404590821441E-07 | 0.367562539993246 | 0.571 | 0.094 | 0.0324238335592255 |
| **RGS4** | 7.06217213950261E-07 | 1.29397621483838 | 0.679 | 0.234 | 0.0329754003709795 |
| **ARX** | 7.09227482273617E-07 | 0.207954231531124 | 0.464 | 0.047 | 0.033115958829802 |
| **SPOCK3** | 7.09558822131753E-07 | 0.346446498541144 | 0.393 | 0.016 | 0.033131430081798 |
| **INS** | 9.1592229249322E-07 | -0.518250224744619 | 1 | 1 | 0.0427671596033859 |

Supplemental Table 6 - Cluster1 in non-diabetics vs T2D

| | p_val | avg_logFC | pct.1 | pct.2 | p_val_adj |
|---|---|---|---|---|---|
| **SP100** | 1.83529982769889E-15 | 0.378505445302907 | 1 | 1 | 8.56956548547445E-11 |
| **PSPHP1** | 2.20744542905235E-13 | 0.192776778346709 | 0.494 | 0.013 | 1.03072249418741E-08 |
| **PRSS2** | 4.30831031436233E-12 | -0.162827815744324 | 0.053 | 0.347 | 2.0116793350852E-07 |
| **ANXA2** | 1.17918322353811E-11 | -0.563211460832181 | 0.707 | 0.96 | 5.5059602256665E-07 |
| **HLA-DQB1** | 4.22582007470112E-09 | -0.118541300184139 | 0.015 | 0.187 | 0.000197316216748019 |
| **SELENOM** | 4.57806548755217E-09 | -0.334792388055403 | 0.57 | 0.867 | 0.000213763611810274 |
| **RBP4** | 1.41672401004439E-08 | 0.852515791041687 | 0.654 | 0.28 | 0.000661510942010028 |
| **BSG** | 1.47361893765802E-08 | -0.385386528856361 | 0.981 | 0.987 | 0.00068807689056066 |
| **SPP1** | 3.93465927290205E-08 | -0.925065270629486 | 0.209 | 0.52 | 0.00183721045429615 |
| **SGIP1** | 7.31732691792908E-08 | -0.164998969371357 | 0.202 | 0.52 | 0.00341667945778862 |
| **CBX6** | 1.8497953593377E-07 | 0.309721421676018 | 0.973 | 1 | 0.00863724947135553 |
| **RNF152** | 2.52502071816576E-07 | -0.161457893110474 | 0.163 | 0.427 | 0.0117900792393314 |
| **DLK1** | 2.6498653797298E-07 | -0.850082485081069 | 0.635 | 0.827 | 0.0123730164175724 |
| **SULF1** | 4.26351994219233E-07 | -0.144019985225868 | 0.152 | 0.413 | 0.0199076536660786 |
| **CAPS2** | 4.40191944204415E-07 | -0.197951798845561 | 0.125 | 0.373 | 0.0205538824507367 |
| **NEFL** | 8.47960453055088E-07 | -0.204532655298121 | 0.042 | 0.213 | 0.0395938174345012 |

Supplemental Table 7 - LDHA_pos_β-cells in non-diabetics vs T2D

| | p_val | avg_logFC | pct.1 | pct.2 | p_val_adj |
|---|---|---|---|---|---|
| **RPS2P46** | 5.85730011951328E-09 | -0.1118549378086 | 0.925 | 0.964 | 0.000273494914480434 |
| **PSPHP1** | 2.70419201167337E-08 | 0.24418410322901 | 0.672 | 0 | 0.00126266837601065 |
| **PRSS2** | 1.90296200240872E-07 | -0.322348427835777 | 0.03 | 0.464 | 0.00888550047784702 |
| **CBX6** | 9.49842470252106E-07 | 0.421997298176448 | 1 | 1 | 0.0443509944634816 |
